# Mononuclear myeloid cells mount an adaptive protective response to intracerebral haemorrhage by activation of NRF2

**DOI:** 10.64898/2026.05.17.725738

**Authors:** James J.M. Loan, Jing Qiu, Jack Barrington, Paul Baxter, Sean McKay, Caoimhe Kirby, Xin He, Christine Lerpiniere, Karina McDade, Meryam Beniazza, Jamie McQueen, Friederike Schaper, Yihui Cheng, Bethany Geary, Owen R. Dando, Colin Smith, Neshika Samarasekera, Rustam Al-Shahi Salman, Barry W. McColl, Giles E. Hardingham

## Abstract

Following intracerebral haemorrhage (ICH), secondary injury contributes to negative outcomes. Here, we describe inter-dependent injurious and adaptive-protective pathways that form a therapeutically modifiable ICH response. In perihaematomal tissue from post-mortem ICH patients, transcriptomic analysis pointed to concurrent perihaematomal activation of the cytoprotective stress-induced transcription factor NRF2 and activation of interferon responses, with mononuclear myeloid cells (microglia and monocyte-derived cells, MMCs) and astrocytes contributing to this signature. In vitro, microglia-astrocyte-neuron co-culture studies revealed that microglial NRF2 activation was induced by clot-conditioned medium and could influence the interferon pathway in both microglia themselves as well as astrocytes. In vivo, MMC-specific deletion of NRF2 inhibited NRF2 target-gene induction, exacerbated interferon pathway activation following ICH, and impaired functional recovery. Pharmacological NRF2 activation suppressed interferon pathway induction post-ICH, and direct interferon pathway antagonism rescued neurological recovery deficits caused by MMC-specific NRF2 deficiency. Thus, augmenting NRF2- or inhibiting interferon-signalling may improve outcomes after ICH.

## INTRODUCTION

Intracerebral haemorrhage (ICH) causes approximately 3 million deaths and 69 million disability-adjusted life years globally each year, accounting for the highest burden of disability of all stroke subtypes^1^. Current treatments for ICH aim to mitigate early injury due to mass effect, including rapid blood pressure lowering and surgical evacuation of haematoma in selected cases^2–4^. However, the consequences of bleeding into the brain are complex, and in the days and weeks following ICH secondary injurious and protective responses influence functional recovery^5,6^ Secondary injury was prioritised as a target for therapeutic development by the HEADS academia-industry roundtable given its therapeutically tractable window^6,7^. To enable this, the cellular and molecular pathways that influence secondary injury and neurological outcomes must be identified and targeted.

Multiple factors are thought to influence secondary injury after ICH. These include inflammation and immune cell processes, oxidative stress and metabolic peturbation^8,9^. Mononuclear myeloid cells (MMCs; including microglia and monocyte-derived macrophages) are innate immune cells that accumulate in the perihaematomal area^10^. Whilst these are thought to be the principal cell types involved in haematoma clearance and haem detoxification, they have potential to amplify injury through inflammatory pathway activation^11,12^. Moreover, MMC sensory and secretory functions more generally enable communication with non-immune cell types^13,14^. This contributes to pathophysiological states, including type 1 interferonopathies, where dysregulation of these functions is thought to underpin a destructive microgliopathy^15^. MMCs therefore are spatially and molecularly well-placed to influence secondary injury through cell intrinsic mechanisms and interactions with other cells.

A potential modulator of secondary injury after ICH is the transcription factor NRF2, which controls expression of a wide range of cytoprotective antioxidant, detoxification, metabolic and proteostasis genes^10,16^. Although constitutively translated, under normal conditions, NRF2 is subject to continuous ubiquitin-mediated degradation via Keap1^17^. Stresses associated with secondary injury after ICH, including altered redox balance and iron-containing haem accumulation, inhibit Keap1-dependent degradation of NRF2, allowing it to enter the nucleus and induce genes containing NRF2 binding sites in their promoters, termed antioxidant response elements^17–20^. Mice globally deficient in NRF2 have increased vulnerability to disease models affecting most major organs, a reflection of its widespread expression^21–23^. In ICH, the cell type-specific roles of NRF2 are unclear. In mouse models of ICH, global NRF2 deficiency worsens outcome and administration of NRF2 activators is protective, highlighting therapeutic possibilities^24^. However, the ubiquitous body-wide roles of NRF2 makes the interpretation of such studies difficult. To realise the potential of NRF2-targeted therapy, a more refined cell-type specific understanding is needed. This is because the consequences of manipulating NRF2 activity in different cell types is likely to have different impacts depending on the role of that cell type, the expression level of NRF2, and the nature of the gene-set within each cell-type that is poised to be activated by NRF2 translocation to the nucleus^25^. Importantly, investigation of cell type-related roles of NRF2 needs to be directed by findings from human ICH cases, which is currently lacking^8^.

In this study we show that gene networks and *in situ* markers of NRF2 activity are induced in human ICH brain tissue and largely localised to cells of mononuclear myeloid lineage, close to the haematoma. We then show through cell-type-specific NRF2 deletion and pharmacological NRF2 activation that MMC NRF2 activation forms part of an adaptive-protective response in a preclinical ICH model through a mechanism involving suppression of harmful microglial interferon signalling, promoting functional recovery.

## RESULTS

### The human brain response to ICH involves prominent induction of NRF2 in perihaematomal myeloid cells

Transcriptome-wide profiling of whole brain tissue from patients with ICH over the first weeks of onset has hitherto not been conducted^8^. We reasoned that it may identify putative regulators of secondary injury and protection. We therefore performed RNA-seq on samples collected from perihaematomal brain tissue obtained at autopsy from a cohort of patients who died 7-27 days after their first ever lobar ICH. This was compared to samples from control brain tissue from age/sex-matched controls who died suddenly of non-neurological disease (Cohort 1; Supplementary Table 1, Supplementary Figure 1). 1389 genes were differentially expressed in perihaematomal brain tissue of patients with ICH (p_adj_<0.05) compared with sudden death controls (Figure 1A). Genes associated with inflammatory responses, iron metabolism and responses to oxidative stress, including *HMOX1, S100A8, NQO1, SLC11A1, IL1R2* and *MMP9,* were induced after ICH. In an unbiased Hallmark gene set enrichment analysis (GSEA), the most enriched gene sets after ICH reflected processes of inflammation, including interleukin and STAT-mediated responses, and response to hypoxia, (Figure 1B). These transcriptional changes implicate altered immune cell composition and/or activation as a core element of the human brain’s response to ICH during the subacute phase.

**Figure 1:**
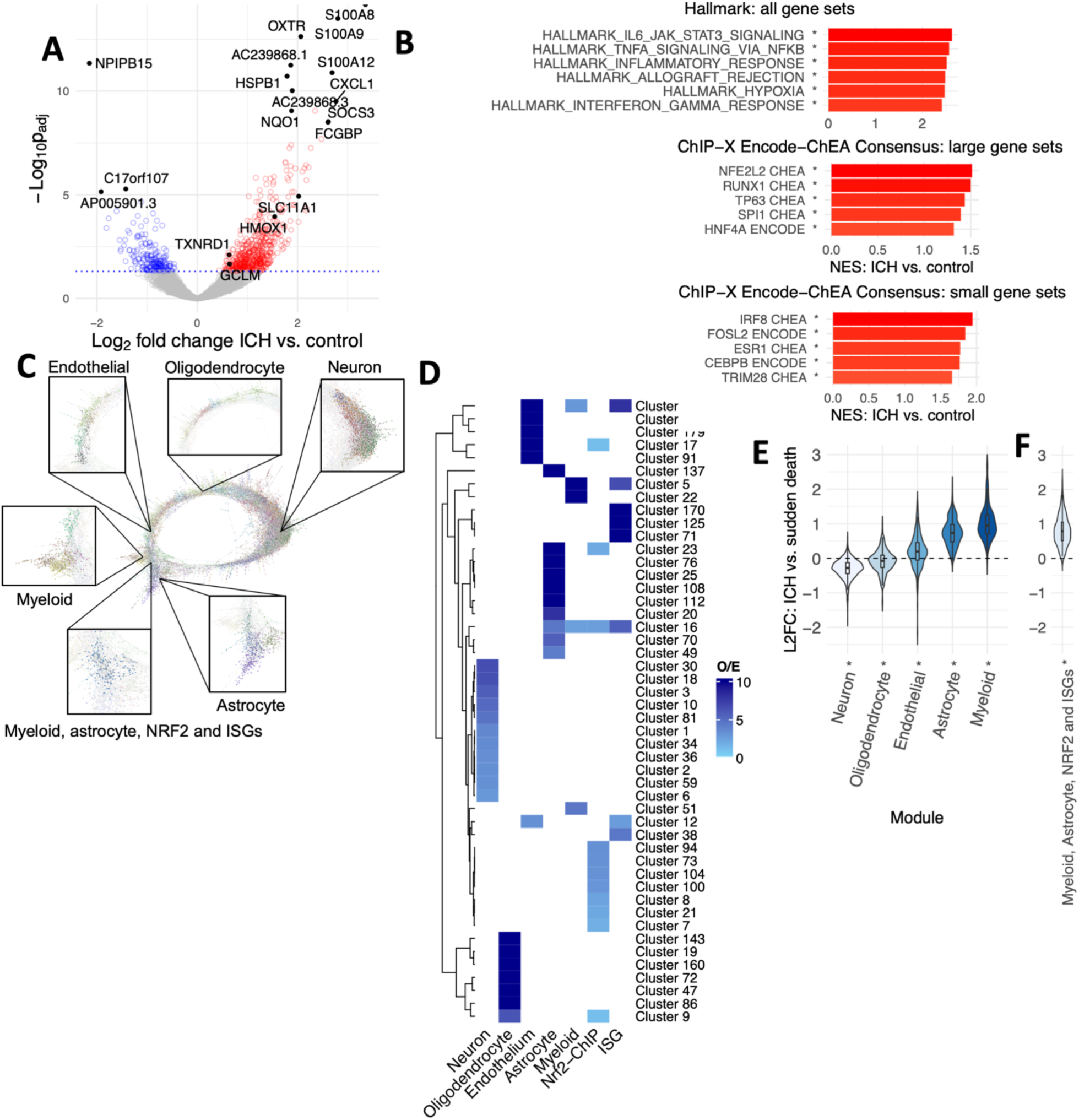
ICH induces the expression of NRF2-regulated genes by myeloid cells and astrocytes in patient brain tissue. **A** Volcano plot demonstrating differential gene expression in perihaematomal brain tissue samples from patients (cohort 1) who died after ICH (n=4) compared with sudden death controls (n=7). Benjamini-Hochberg (B-H) adjusted p-values (p_adj_). **B** Bar chart of Hallmark Gene Set Enrichment Analysis (GSEA) derived normalised Enrichment Score (NES) for ICH vs. sudden death. All *B-H p_adj_<0.05. **C** Gene coexpression network graph. Nodes indicate genes and edges represent coexpression across cohort 1 samples; Pearson R>0.964, R-ranked 𝜅-nearest neighbours (𝜅NN) edge reduction 𝜅=5, Markov Clustering (MCL) inflation value 1.25. Clusters are represented by node colour and gene modules are of groups of clusters enriched for specific cell types (**D)**. **D** heatmap of enrichment (O/E; observed/expected) of clusters (**C**) for cell type specific gene sets as well as NRF2 target genes identified by a previous ChIP-seq analysis (NRF2-ChIP) and Hallmark interferon-stimulated genes (ISGs). Only gene sets showing significant enrichment (Bonferroni-adjusted Fishers p<0.05) displayed. As such, the clusters comprising each module are: neuron – 1-3, 6, 10, 18, 30, 34, 36, 59, 81; oligodendrocyte –19, 47, 72, 86, 143, 160; endothelial – 12, 17, 91, 166, 179; astrocyte – 20, 23, 25, 49, 70, 76, 108, 112, 137; myeloid – 5, 22, 51; myeloid, astrocyte and NRF2 – 16. **E-F** Violin plot of log_2_ fold changes (L2FC) after ICH vs. control for genes in each module (E; GSEA ICH vs. Sudden death control: Neuron NES=-3.5, p_adj_<0.0001; Oligodendrocyte NES=-1.5, p_adj_<0.0001; Endothelial NES 1.9, p_adj_<0.0001; Astrocyte NES 3.5, p_adj_<0.0001; Myeloid NES 3.5, p_adj_<0.0001) or cluster 16 (F; NES=3.13, p_adj_<0.0001).

To explore potential transcriptional regulatory factors contributing to the gene expression signature, we next performed an analysis of the ChIP-X ENCODE-ChEA consensus gene sets, stratified by gene set size^26^. By filtering significantly enriched genes to identify the five largest and smallest sets, we found that the most significantly enriched large gene set was of genes regulated by the transcription factor NRF2, encoded by the gene *NFE2L2* (Figure 1B), and exemplified by the induction of classical NRF2 target genes such as *HMOX1* and *NQO1*. Other highly enriched transcription factors included RUNX1, which regulates myeloid and lymphoid cell differentiation, and IRF8, an interferon regulatory factor which also influences myeloid cell differentiation. These indicate that, in addition to altered immune cell activation/composition, human ICH brain response involves activation of NRF2, known to have the potential to modulate inflammatory processes^27^.

To define changes in gene expression attributable to different cell types, we developed a gene co-expression network comprising all expressed genes in each sample. This revealed clusters of co-expressed genes which are enriched for genes specifically expressed by neurons, cells of myeloid origin, astrocytes, endothelial cells and oligodendrocytes (Figure 1C-D)^28^. By combining clusters selectively enriched for single cell types we created cell type specific “modules” of genes. We found that the overall expression of genes comprising either astrocyte or myeloid cell modules increased after ICH compared with sudden death controls (Figure 1E). This may reflect an increased relative abundance of each of these cell types in perihaematomal tissue and/or their altered reactive status. To investigate whether the NRF2 activation identified previously was associated with a specific cell type, we next tested gene modules for enrichment with NRF2 target genes. We defined NRF2 target genes as those identified by a previous comprehensive NRF2 chromatin immunoprecipitation with parallel sequencing (ChIP-seq) study^22^. This showed that cluster 16, which was elevated after ICH compared with sudden death controls, was enriched for NRF2 target genes and was the only cluster enriched for both myeloid and astrocyte-specific genes (Figure 1C-F, Supplementary table 2). This points to myeloid cells and/or astrocytes as significant contributors to the NRF2 activation signature found in perihaematomal tissue of ICH patients.

To investigate the spatiotemporal evolution of NRF2 activation after ICH, we used GSEA to compare transcriptional responses of perihaematomal brain tissue with contralateral anatomically matched tissue of patients who died either within 48 h of ICH symptom onset (acute) or between 4-12 days of ICH onset (subacute; Cohort 2; Supplementary Table 1 and Supplementary Figure 2). We found that NRF2 target gene sets, and cluster 16 genes (Figure 1D) were enriched in perihaematomal tissue at both acute and subacute time points compared with anatomically-matched contralateral tissue (Supplementary Figure 3). This enrichment was greater in perihaematomal tissue of patients who died between 4-12 days post-ICH than those who died within 48h of symptom onset (Supplementary Figure 3). This demonstrates that in patients with ICH, NRF2 pathway activation is detectable within 48h of ICH onset and becomes more pronounced over the following days. The consistency of differences between perihaematomal and contralateral tissue with those in our case vs. control analysis indicates that these are a response to ICH, rather than underlying conditions such as small vessel disease.

Our differential expression and co-expression network analyses suggested that MMCs (e.g. monocyte-derived cells (MdC), microglia) and astrocytes are cellular sources of NRF2 target gene induction in ICH patients. To corroborate this, we used *in situ* dual mRNA hybridisation-immunohistochemistry to determine if *HMOX1* mRNA is colocalised with the microglial and macrophage marker IBA1 in post-mortem brain tissue from patients who died after ICH and controls who died suddenly of non-neurological disease (Cohort 3; Supplementary Figure 4 and Supplementary Table 3). *HMOX1* is a classical NRF2 target gene that encodes haem oxygenase 1, an inducible enzyme required for the catabolism of haem^29^. We found that in perihaematomal tissue there was a marked increase in IBA1 immunostaining and *HMOX1* expression (figure 2A-F). *HMOX1* transcript puncta were clearly localised to IBA1+ cells in perihaematomal tissue, that had an enlarged amoeboid morphology, consistent with a reactive MMC state. Quantification indicated a ∼10-fold greater abundance of *HMOX1* transcripts in IBA1+ compared to IBA1- cells (Figure 2A-F). In contrast, immunostained OLIG2+ oligodendrocytes and GFAP+ astrocytes were found to express relatively less *HMOX1* (Figure 2G-J).

**Figure 2:**
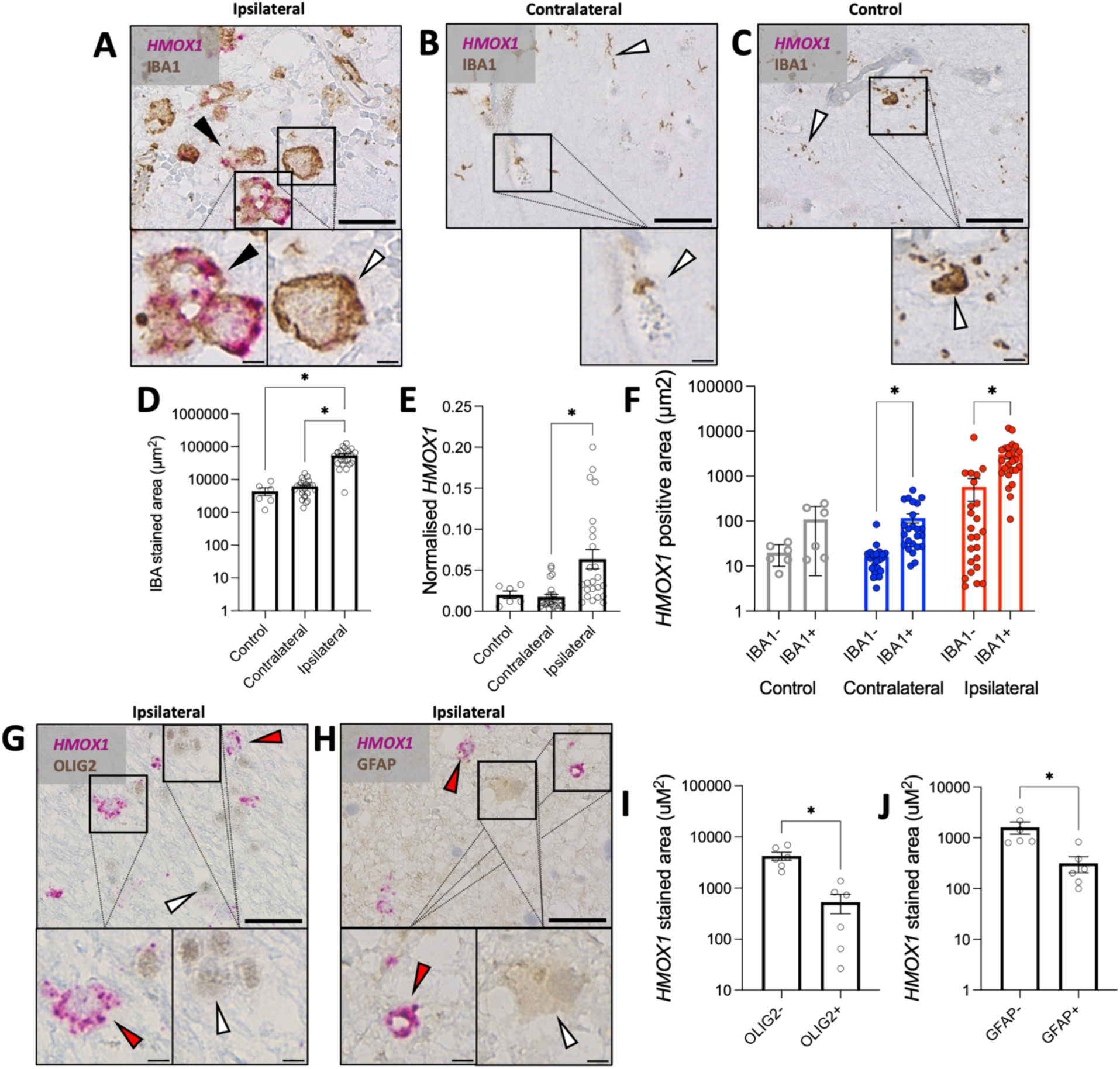
Dual in-situ RNA hybridisation-immunohistochemistry reveals induction of the NRF2-regulated cytoprotective gene *HMOX1* by MMCs in perihaematomal patient brain tissue. A-C Representative images of perihaematomal. (**A**), contralateral (**B**), and control (**C**) brain tissue stained for *HMOX1* mRNA (Fast Red; magenta) and IBA1 protein (DAB; brown) from patients (cohort 3) who died after ICH (n=24) or sudden non-neurological death controls (n=6). **D** Quantification of IBA1 stained area in 10 fields of view (FOV) per patient. Brown-Forsythe one-way ANOVA: F*(2-24.4)=65.9; p<0.0001. Dunnett’s T3 *post-hoc* multiple comparisons control versus contralateral p=0.52, control versus ipsilateral p<0.0001, contralateral versus ipsilateral p<0.0001. **E** Quantification of *HMOX1* stained area normalised to IBA1 stained area for 10 FOV per patient. Brown-Forsythe one-way ANOVA: F*(2-30.3)=15.1; p<0.0001. Dunnett’s T3 *post-hoc* multiple comparisons control versus contralateral p=0.77, control versus ipsilateral p=0.052, contralateral versus ipsilateral p<0.0001. **F** Quantification of *HMOX1* stained area in IBA1 stained (IBA1+) regions and non-IBA1 stained (IBA1-) regions of 10 FOV per patient. Geisser-Greenhouse corrected repeated measures two-way ANOVA comparison of matched perihaematomal and contralateral samples. IBA1 staining: F(1-23)=16.4; p=0.0005. Tissue location: F(1-23)=23.98; p<0.0001. IBA1 x Tissue location: F(1,23)=14.5; p=0.0009. Holm-Šidák *post-hoc* multiple comparisons corrected p-values for comparison of IBA1- versus IBA1 positive regions perihaematomal p=0.0013; contralateral samples p=0.0013. Control sample values shown for illustration but statistical analysis not performed **G-H** Representative images of brain tissue from patients who died after ICH (n=6, cohort 3). For this analysis we selected patients expressing the highest levels of *HMOX1* staining in the previous analysis. Tissue stained for *HMOX1* mRNA (Fast Red; magenta) and either OLIG2 (**G**) or GFAP (**H**) protein (DAB; brown). **I-J** Quantification of *HMOX1* stained area in OLIG2 (**I**) or GFAP (**J**) positive or negatively stained regions of 10 FOV per patient. Paired two-tailed t-test p-values OLIG2 positive versus negative t=4.2, df=5, p=0.008 (**I**); GFAP positive versus negative t=3.5, df=5, p=0.018 (**J**). Main scale bars 30µm, insert 5µm. Black arrows indicate DAB and FastRed double stained cells, white arrows indicate DAB single stained cells and red arrows indicate FastRed positive, DAB negative puncti. All error bars +/- standard error of the mean (SEM)

To further investigate MMC-specific spatiotemporal trends in NRF2 target gene expression we used a probe-based spatial transcriptomics platform (CosMx) to measure the expression of 6175 genes in relation to the haematoma margin in patients with ICH at an acute and a chronic time point, as well as in patients who died suddenly of non-neurological disease (Cohort 4; Supplementary table 4). Counts were assigned to regions that were defined as putative cells according to staining with DAPI, rRNA, histone, GFAP and CD68 and these cells were annotated with cell type using deep generative mapping and a human disease brain taxonomy (Figure 3A, Supplementary figure 5).^30^ Focusing on MMCs, we observed spatial and temporal changes in gene expression that aligned with our bulk RNA sequencing findings, including marked perihaematomal induction of *SPP1* and *HMOX1*, as well as induction of NRF2 target genes (Figure 3B-E, Supplementary figures 6,7) in perihaematomal tissue. Whilst NRF2 target gene expression was submaximal in the acute group, this was further elevated in the chronic group (Figure 3B-E), consistent with the earlier bulk RNA-seq data. Previously annotated sets of microglial genes associated with neurodegenerative disease were markedly induced,^31^ whereas those associated with homeostatic microglia were more modestly influenced by proximity to haematoma (Figure 3B-E).

**Figure 3:**
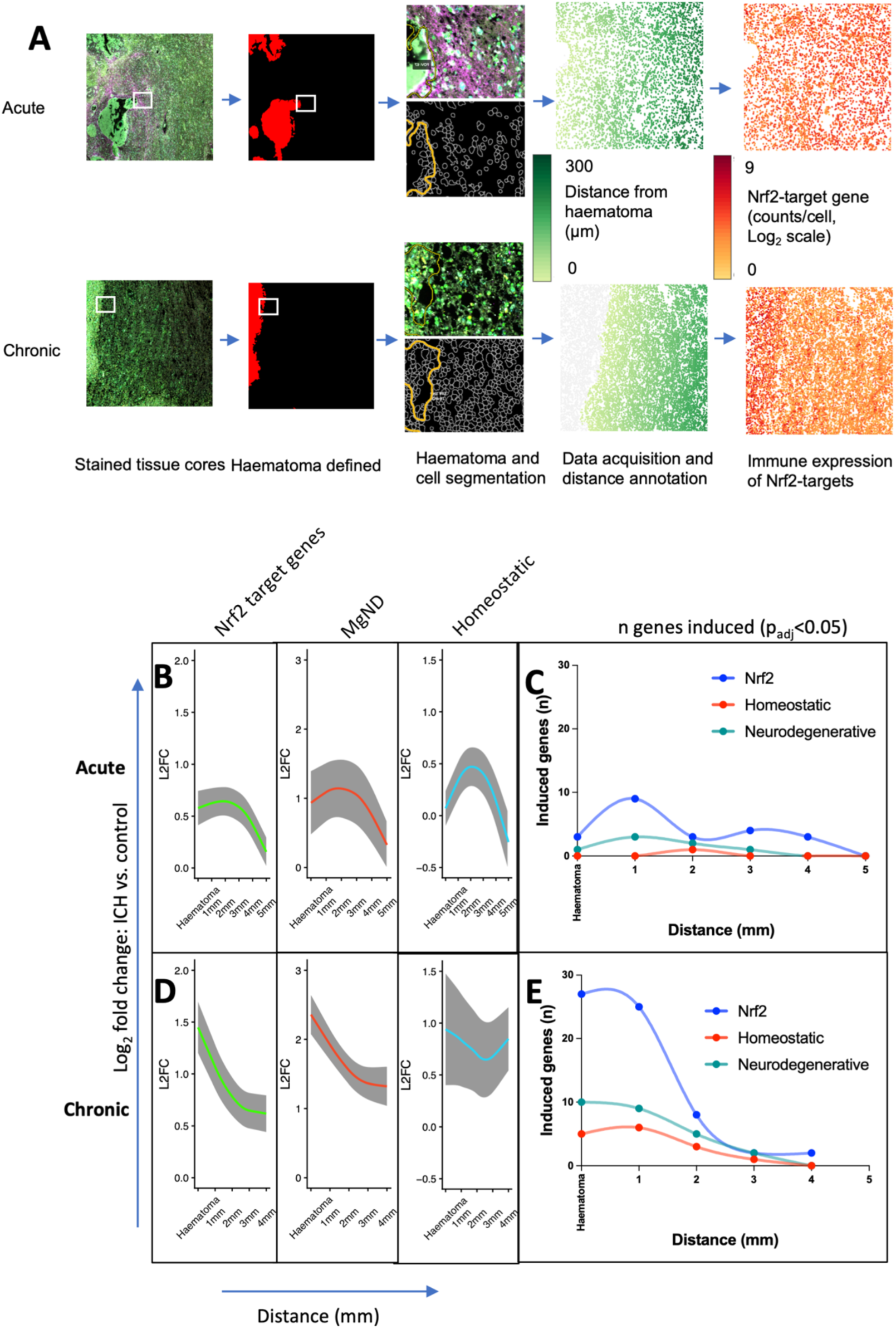
Single cell spatial transcriptomic analysis of patient brain tissue shows perihaematomal co-induction of NRF2 target genes and ISGs by brain immune cells. **A** Analysis pipeline with representative sections from patients who died in the acute (<48h) or chronic (13-41 days) phases after ICH symptom onset. Tissue cores were stained for DAPI (white), rRNA (cyan), Histone (green), GFAP (purple) and CD68 (yellow) and cells and haematoma were segmented. The mRNA expression was then determined within selected fields of view. Segmented cells were annotated with cell type and NRF2 target and ISG expression determined per cell. **B-E** Expression of selected gene sets by segmented cells annotated as containing immune cells in the acute (**B-C**) and chronic (**D-E**) timepoints. **B,E** Log_2_ fold change (L2FC) in expression of differentially expressed (p_adj_<0.05 acute/chronic vs. control) genes in sets of previously annotated NRF2 targets, genes associated with microglia in neurodegenerative disease (MgND) and homeostatic states. Solid line and grey margin indicate mean L2FC with standard error of the mean across genes within a set. Repeated measures mixed-effects analysis of fixed associations of time and distance with gene and subject random effects: NRF2 distance F(4-689.0)=8.1 p<0.001, timepoint F(1-6.0)=4.4 p=0.08, distance*time F(4-689.0)=2.8 p=0.026; MgND distance F(4-215.5)=4.7 p=0.0012, timepoint F(1-5.98)=0.17 p=0.70; distance*timepoint F(4-215.5)=0.43 p=0.79; Homeostatic distance F(4-137.3)=3.5 p=0.0092, timepoint F(1-6.2)=4.9 p=0.067, distance*timepoint F(4-137.3)=3.0 p=0.021. **C,E** Smoothed spline of the number of statistically significantly induced (p_adj_<0.05 vs. control) genes per gene set, distance from haematoma and timepoint. n= 4 acute, 4 chronic, 4 controls.

Overall, our findings indicate that, in perihaematomal brain tissue of patients with ICH, NRF2 target gene activation is prominent and enriched in MMCs. The temporal trajectory indicates a progressive induction during the first days of ICH onset, that remains prominent up to a median of 21 days (Supplementary table 4) of ICH onset, placing the NRF2 response at a critical period of haematoma evolution, and suggesting its potential influence on secondary injury.

### Myelomononuclear NRF2 activation in a preclinical ICH model

Having identified NRF2 pathway activation in MMCs after ICH in humans, we next determined how NRF2 expressed in these cells regulates their responses to ICH. For this, we used a preclinical model combined with MMC-specific deletion of NRF2. We used intrastriatal injection of collagenase to induce ICH in mice, a method which causes deep haemorrhages with similar temporal haematoma evolution to spontaneous ICH in humans (Figure 4A)^32,33^. We initially defined responses to ICH in *Cx3cr1^+/+^:Nfe2l2^fl/fl^* mice, which express normal levels of NRF2. We used fluorescence activated cell sorting (FACS) to separately isolate microglia and monocyte derived cells (MdCs), and profiled their transcriptomes by RNA sequencing (Figure 4; Supplementary Figure 8A; Supplementary table 5). Cytometry data collected during sorting showed that ICH caused a marked accumulation of MdCs 3 days after ICH as well as an increase in numbers of the resident microglial population (Figure 4B-E). The purity of sorted samples was validated by analysis of the expression of microglial specific gene *P2ry12* and MdC specific genes *Ccr2* and *Ly6c2* (Supplementary figure 8B-D). Within three days of ICH onset, microglia showed an altered transcriptome reflective of their reactivity, including suppressed *P2ry12*, which is highly expressed by homeostatic microglia but down-regulated under inflammatory conditions (Figure 4F).^34^ Marked induction of *Hmox1* was observed and enrichment of glycolysis, oxidative phosphorylation, and cell cycle-associated gene sets was evident in ICH microglia (Figure 4F,G). NRF2 target genes (defined using the same previously published ChIP-seq study that we used to annotate human gene expression modules^22^) were overall increased in expression in microglia after ICH compared with sham microglia, indicative of NRF2 activation (Figure 4H). Compared with ICH microglia, ICH-recruited MdCs exhibited relatively greater expression of *Hmox1*, *Il1b*, and *Tgfbi*, as well as enrichment of gene sets associated with immune reactivity and redox regulation (Figure 4I,J and Supplementary Figure 9). Further, genes previously shown to be induced by microglia in neurodegenerative disease contexts^31^ were increased after ICH in microglia and further elevated in MdCs, whilst those which had been previously found to be enriched in homeostatic microglia^31^ were suppressed by ICH in microglia and, as expected, expressed at lower levels in MdCs (Supplementary Figure 10). Using mass spectrometry imaging (MSI) in wild type mice, we identified a trend towards induction of proteins which are highly expressed by astrocytes, MMCs and endothelial cells, which peaked at day 1-3 and returned to baseline by day 28 (Supplementary figure 11).^35^ This supports the use of a day 3 timepoint for analysis of the influence of NRF2 on inflammatory responses after ICH, however, individual protein findings must be interpreted with caution as they did not retain statistical significance following adjustment for multiple testing, and independent replication is needed. Overall, these data from a preclinical ICH model demonstrate that, similarly to the human brain, ICH triggers NRF2-regulated gene network induction in MMCs, raising the possibility that NRF2 in MMCs may influence the trajectory of injury post-ICH in the mouse model.

**Figure 4:**
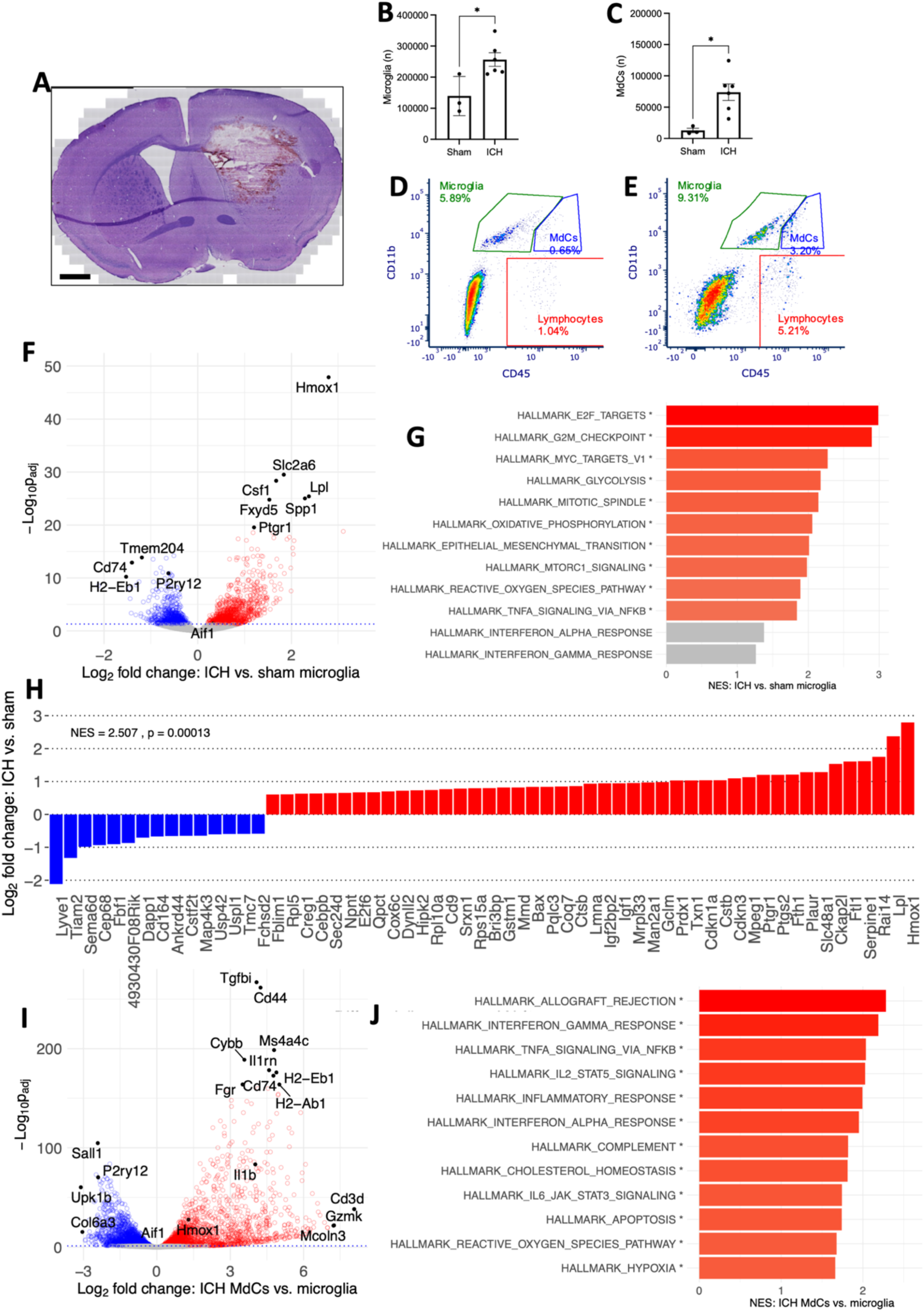
Experimental ICH causes MMC recruitment and activation in mice. **A** Representative haematoxylin and eosin-stained coronal mouse brain section demonstrating a striatal haemorrhage and associated perihaematomal oedema at day three after stereotactic injection of bacterial collagenase. Scale bar 1mm. Absolute numbers of ASCA-2-, O4-, Ly6G-, Cd11b+,Cd45^int^ microglia (**B**; unpaired two-tailed Mann-Whitney U p=0.048) and ASCA-2-, O4-, Ly6G-, Cd11b+,Cd45^hi^ MdCs (**C**; unpaired two-tailed Mann-Whitney U p=0.024) per whole brain of NRF2^LoxP^ mice. Error bars +/- SEM. Unpaired Mann-Whitney U test p-values graphed. **D-E** representative flow cytometry plots of Cd11b versus Cd45 fluorescence from ASCA-2, O4 and Ly6G negative cell populations from sham (**D**) and ICH (**E**) mice. **F** Volcano plots of differential gene expression after ICH vs. sham microglia. DESeq2 log_2_ fold changes and Benjamini-Hochberg adjusted p-values (p_adj_). **G** Hallmark GSEA plots of most positively (red) and negatively (blue) enriched gene sets after ICH compared with sham, in microglia, ranked using NES. Interferon-associated gene sets plotted for illustration (grey, where not statistically significantly enriched). *B-H adjusted p<0.05. **H** Bar chart of log_2_ fold change (L2FC) after ICH versus sham microglia in expression of highly differentially expressed (>1.5 fold change, p_adj_<0.05) NRF2 target genes identified by a previously published ChIP Seq experiment^4^. GSEA test of overall set enrichment with ICH NES=2.5, p=0.0001 **I** Volcano plots of differential gene expression in MdCs versus microglia after ICH. DESeq2 log_2_ fold changes and Benjamini-Hochberg adjusted p-values (p_adj_). **J** Hallmark GSEA plots of most positively (red) and negatively (blue) enriched gene sets in MdCs, compared with microglia after ICH, ranked using NES. n=3 sham and n=5 ICH microglia, n=6 ICH MdCs.

### MMC NRF2 activation facilitates protective adaptive responses to ICH

To determine the functional importance of NRF2 expressed within MMCs on the responses to ICH, we employed the MMC-specific Cx3cr1-Cre-mediated excision of *Nfe2l2* exon 5, which removes the DNA-binding domain of NRF2 as well as disrupting the Neh3 and Neh5 transcriptional activation domains, rendering any translated protein non-functional^36^. This prevents the activation of NRF2-driven gene expression in MMCs of *Cx3cr1*^Cre/+^:*Nfe2l2^f^*^l/fl^ (NRF2^ΔMMC^) mice. *Cx3cr1*^+/+^:*Nfe2l2^f^*^l/fl^ (NRF2^LoxP^) mice were used as controls. Using RT-qPCR and exon differential expression analysis of RNAseq data, we confirmed the selective loss of *Nfe2l2* exon 5 expression in microglia, CNS- and peripheral- MMCs (Supplementary Figure 12) of NRF2^ΔMMC^ mice compared NRF2^LoxP^ controls. Astrocytic expression was unaffected (Supplementary Figure 12).

Using immunohistochemistry, we found that, in control NRF2^LoxP^ mice, IBA1-positive MMCs were recruited to the perihaematomal region and exhibited HMOX1 induction (Figure 5B,F,G,H) in a similar fashion to that observed in human brain tissue (Figure 2, Figure 3) and consistent with mouse transcriptome data (Figure 4). However, although NRF2 deletion did not affect perihaematomal recruitment of Iba1-positive microglia and MdCs, it did suppress microglial transition to a reactive phenotype, indicated by the reduced perihaematomal suppression of the homeostatic microglial marker P2Y12 (Figure 5C-E) and impaired expression of NRF2 target gene HMOX1 (Figure 5F-H). For both HMOX1 and P2Y12, we found that effects of NRF2 deletion were dependent on the distance from the haematoma surface, indicating that NRF2 is induced by ICH-induced stresses and may serve to regulate local responses to ICH rather than influencing the expression of these factors in the resting state (Figure 5C, F). This aligns with the spatial pattern of perihaematomal Nrf2-target gene expression identified in MMCs of patients with ICH (Figure 2, Figure 3). Also consistent with this, we did not observe an effect of NRF2 deletion on HMOX1 or P2Y12 expression in contralateral tissue or that of sham-operated mice (Supplementary Figure 13).

**Figure 5:**
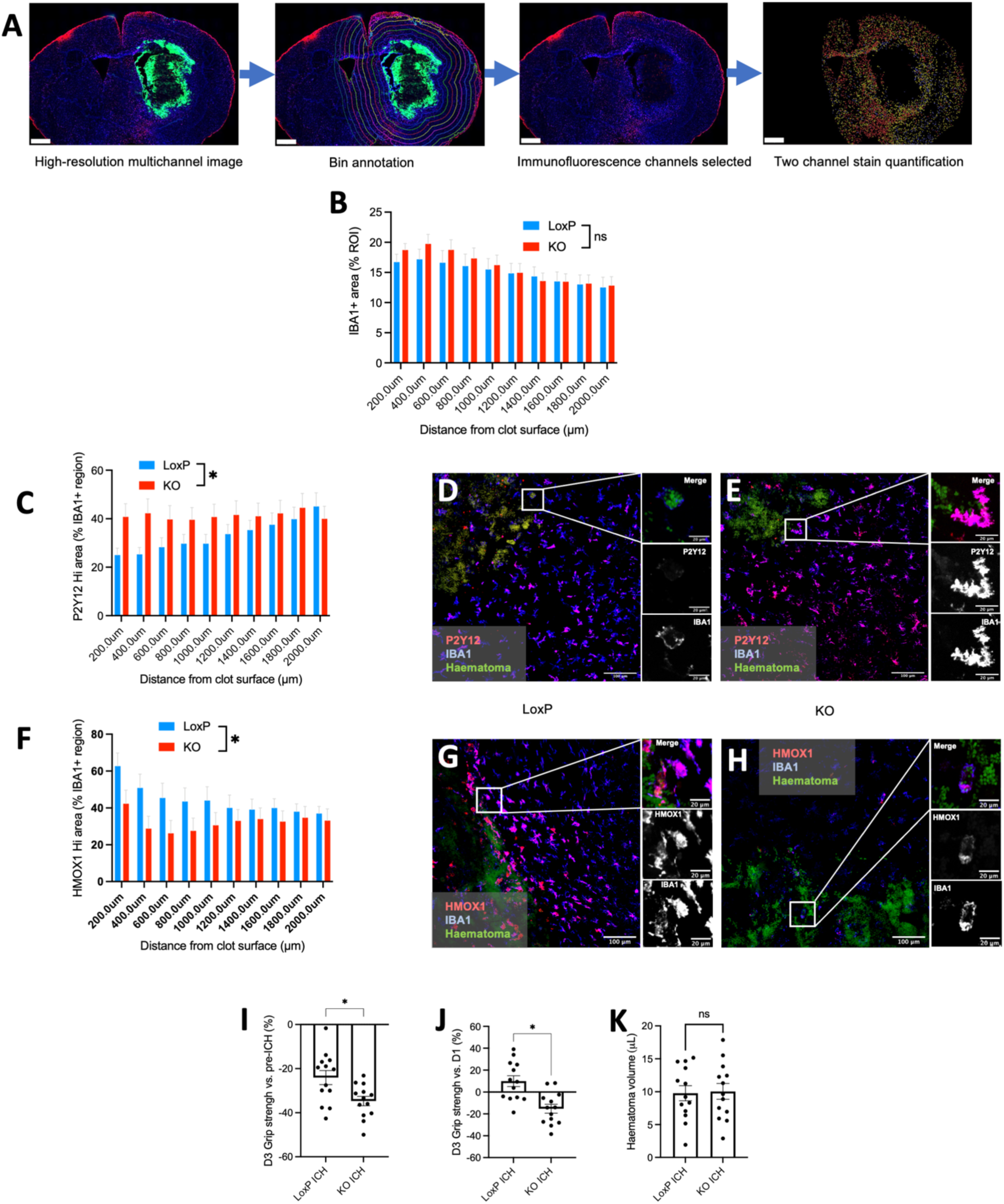
NRF2 expression supports protective MMC responses to haemorrhage. **A** Flow diagram of analysis pipeline for immunohistochemistry. High resolution multichannel immunofluorescence images: haematoma (FITC) green; IBA1 (Alexa 647) blue; P2Y12/HMOX1 (Alexa 555) magenta. Haematoma was defined semiautomatically and 200µm wide regions of interest (ROI) were defined expanding from the haematoma surface. Finally, IBA1 stained regions were segmented and P2Y12 or HMOX1 staining quantified within these. Scale bars 800µm. **B** Percentage of each ROI positively stained for IBA1 according to distance from haematoma surface; n=12 mice/genotype. Two-way repeated measures ANOVA: Distance from haematoma*Genotype F(9-198)=1.7, p=0.010; Genotype F(1-22)=0.15, p=0.70; Distance F(1.1-25.0)=25.0, p<0.0001; Subject F(22-198)=62.8, p<0.0001. Post-hoc comparison of Iba1 staining by genotype at each distance ROI using the Holm-Šidák method yielded no statistically significant findings. **C** P2Y12 stained area as a percentage of the total IBA1 stained area according to distance from haematoma surface; n=12 mice/genotype. Two-way repeated measures ANOVA: Distance*Genotype F(9-198)=2.8, p=0.0042; Genotype F(1-22)=2.0, p=0.17; Distance F(1.5-33.1)=3.5, p=0.055; Subject F(22-198)=24.2, p<0.0001. Nil statistically significant post-hoc differences. **D,E** Representative images of P2RY12 (magenta) and IBA1 immunofluorescence labelled sections from NRF2^LoxP^ (**D**) and NRF2^ΔMMC^ (KO; **E**) mice. **F** HMOX1 stained area as a percentage of the total IBA1 stained area according to distance from haematoma surface; n=13 mice/genotype. Two-way repeated measures ANOVA: Distance*Genotype F(9-216)=4.1, p<0.0001; Genotype F(1-24)=1.8, p=0.18; Distance F(1.2-28.4)=8.4, p=0.0052; Subject F(24-216)=57.16, p<0.0001. Nil statistically significant post-hoc differences. **G-H** Representative images of HMOX1 (magenta) and IBA1 immunofluorescence labelled sections from NRF2^LoxP^ (**G**) and NRF2^ΔMMC^ (**H**) mice. **D,E,G,H** Representative maximum intensity projection of confocal z-stacks with high resolution representative merged and single channel fields of view. Scale bars 100µm, insert 20µm. **I-J** Percentage change in forelimb grip strength at day-three after ICH versus pre-ICH (**I**; Welch’s two-tailed t-test: t=2.8; df=20.9; p=0.011) and versus day-one after ICH (**J**; Welch’s two-tailed t-test: t=3.9; df=23.1.4; p=0.0007). **K** Day-three haematoma volume NRF2^LoxP^ versus WT mice, n=13/group, Welch’s two-tailed t-test: t=0.17; df=24.0; p=0.87. All error bars +/- SEM.

We next studied the impact of MMC-specific NRF2 deletion on the neurological outcome after ICH. Weakened forelimb grip strength (our pre-specified endpoint) caused by ICH was aggravated in NRF2^ΔMMC^ mice compared with their NRF2^LoxP^ littermates (Figure 5I) showing a worsening of performance, rather than the usual recovery that takes place in the first three days after ICH onset seen in NRF2^LoxP^ littermates (Figure 5J and Supplementary Figure 14A). Other measures of wellbeing showed a similar pattern of impaired recovery in NRF2^ΔMMC^ mice compared to NRF2^LoxP^ littermates (Supplementary Figure 14). Whilst haematoma volume and neuronal pyknosis showed no genotype effect (Figure 5K, Supplementary figure 15),) we found a trend of reduced abundance of proteins required for normal neuronal and astrocyte function such as KCNC1, PPP1R12A and FUS close to the haematoma in NRF2^ΔMMC^ mice compared with NRF2^LoxP^ littermates^35^. Conversely, NPL which is induced by lipid-droplet rich microglia in association with neuronal injury^35,37^ displayed a trend towards induction in perihaematomal tissue from NRF2^ΔMMC^ mice but not NRF2^LoxP^ controls (Supplementary figure 16). Overall, these data show that MMC-expressed NRF2 regulates a protective response that limits worsening and/or promotes initial functional recovery after ICH, rather than by influencing initial ICH severity. In line with previous findings, we found that *Cx3cr1* haploinsufficiency and Cre-recombinase expression did not influence either behavioural or histological phenotypes after ICH (Supplementary Figure 17)^38^. Thus, MMC-expressed NRF2 regulates the perihaematomal microglial/macrophage response to ICH and contributes to functional recovery.

### MMC NRF2 deletion suppresses NRF2 target gene induction and drives an exaggerated interferon response

Using RNA-seq of FACS-sorted microglia and MdCs, we found that, after ICH, MMC NRF2 deletion reduced the expression of NRF2 target genes involved in haem detoxification and antioxidant systems (Figure 6A). At a transcriptome-wide level, the induction and repression of genes by microglia in response to ICH was reduced by MMC NRF2 deficiency (Figure 6B-C Supplementary Figure 18). NRF2 target genes that were repressed by myelomononuclear NRF2 deletion in mice with ICH were found to be induced by microglia from NRF2^LoxP^ mice after ICH, compared with those which underwent sham procedures (Figure 6D)^22^. These genes were more highly expressed by MdCs than microglia and they were also repressed in NRF2^ΔMMC^ MdCs compared with those from NRF2^LoxP^ mice after ICH (Figure 6E)^22^. This indicates that in both cell types, ICH induces NRF2 target genes in an NRF2-dependent manner. Because the cell type profile of MdCs after sham and ICH surgery likely differ (Figure 4I-J; Supplementary figure 9), it was not appropriate to directly compare sham and ICH MdCs.

**Figure 6:**
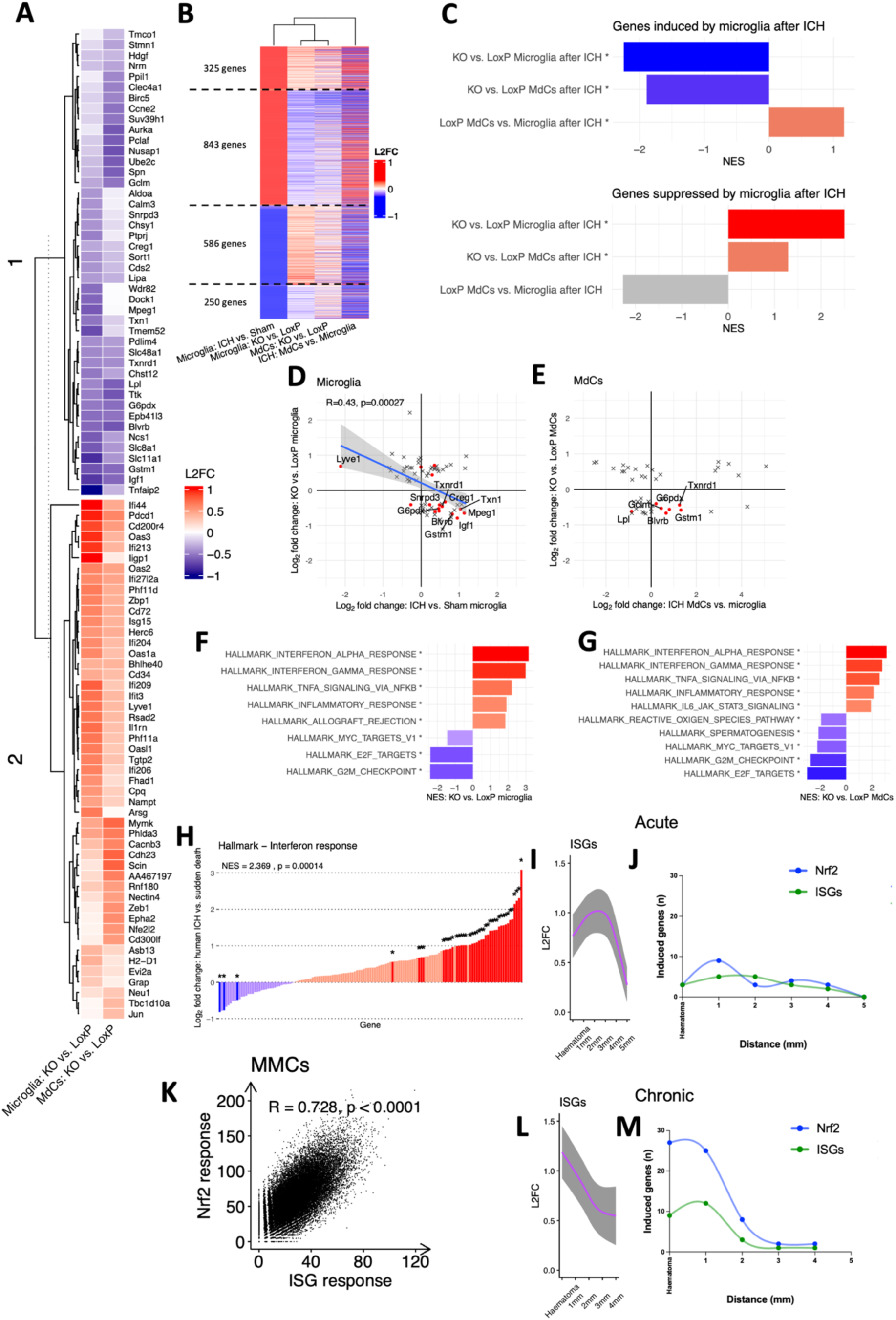
Transcriptome-wide analysis finds that NRF2 deletion blunts adaptive microglial and monocyte derived cell (MdC) responses to ICH and causes the exaggerated induction of ISGs. **A** Heatmap of DESeq2 log_2_ fold change (L2FC) of genes which are differentially expressed (padj<0.05) by microglia and/or MdCs after ICH. Only genes with consistent positive or negative L2FCs plotted. **B** Heatmap of genes that are differentially expressed by microglia after ICH versus Sham. Genes are grouped according to direction of effect of ICH and NRF2 genotype with the majority of genes responding to NRF2 deletion in a direction which attenuates putative adaptive response to ICH. For each gene, the corresponding L2FC for NRF2^ΔMMC^ versus NRF2^LoxP^ and for MdCs versus microglia after ICH is shown. **C** Bar plots of Gene Set Enrichment Analysis (GSEA) Normalised Enrichment Scores (NES) for gene sets (**B**) that are induced or suppressed by microglia after ICH versus Sham in NRF2^ΔMMC^ versus NRF2^LoxP^ microglia, MdCs and in NRF2^LoxP^ MdCs versus Microglia after ICH. *p_adj_<0.05 **D-E** Scatter plots of L2FC in expression of microglial (**D**) and MdC (**E**) differentially expressed genes in NRF2^ΔMMC^ mice versus NRF2^LoxP^ mice with ICH indicate that NRF2 deletion constrains the expression of NRF2 target genes induced by ICH. NRF2 target genes identified by ChIP sequencing are highlighted in red.^4^ For microglial genes (**D**), the effects of NRF2 deletion and ICH are shown with a simple linear regression line (blue) with 95% confidence intervals (gray). For MdC genes (**E**), the effects of NRF2 deletion and the relative expression in NRF2^LoxP^ MdCs compared with NRF2^LoxP^ microglia after ICH are shown. **F-G** Bar plots of GSEA NES for the five most highly positively and negatively enriched Hallmark gene sets in NRF2^ΔMMC^ vs. NRF2^LoxP^ microglia (**F**) and MdCs (**G**) after ICH.^47^ n=5 ICH NRF2^LoxP^ microglia/MdCs, and 6 ICH NRF2^ΔMMC^ microglia, 6 ICH NRF2^ΔMMC^ MdCs, 3 sham NRF2^ΔMMC^/^LoxP^ microglia/MdCs. **H** bar plot of L2FC in hallmark interferon-alpha and -gamma response gene sets in perihaematomal brain tissue from four patients in cohort 1 who died after ICH vs. seven of those who died suddenly of non-neurological disease. Genes increased after ICH are red and those that were decreased are blue. Individually differentially expressed genes (p_adj_<0.05) are indicated by saturated colouring and asterisks. GSEA test of overall enrichment NES=2.4, p=0.0001. Log2 fold change (L2FC; **I,J**) and number of significantly differentially expressed (p_adj_<0.05 Hallmark ISGs in human MMCs identified using spatial transcriptomics after ICH compared with sudden death at acute (**I,J**) and chronic (**L,M**) timepoints. Repeated measures mixed-effects analysis of fixed associations of time and distance with gene and subject random effects. ISGs distance F(4-338.2)=2.5 p=0.042, timepoint F(1-6.1)=0.0005 p=0.98, distance*timepoint F(1-338.2)=0.38 p=0.82. **K** summed expression of Nrf2 target genes and hallmark ISGs per MMC in human brain tissue at all distances and times after ICH. Pearsons R^2^ correlation coefficient displayed. n= 4 acute, 4 chronic, 4 controls.

Despite blunting the overall expression of ICH response genes, we found that NRF2 deletion caused an abnormally elevated MMC interferon-stimulated gene (ISG) response. After ICH, Hallmark interferon-⍺ and interferon-𝛾 response genes were enriched in NRF2^ΔMMC^ microglia and MdCs compared with NRF2^LoxP^ mice (Figure 6F-G). Using genes identified in a dedicated transcriptomic comparison of responses to interferons-⍺ and 𝛾^39^, we found that whilst ICH in NRF2^LoxP^ mice caused some ISGs to be induced and others suppressed, myelomononuclear NRF2 deletion triggered the consistent induction of interferon-⍺ response gene sets after ICH (Supplementary figure 19). Consistent with this, in sham operated mice, NRF2 deletion caused a much smaller change in ISGs and attenuation of NRF2-regulated genes (Supplementary Figure 18C-D). These data indicate that MMC NRF2 deficiency causes an exaggerated type-1 interferon response after ICH^40^. When we interrogated the transcriptomic responses of human patient brain tissue to ICH we found marked ISG induction which was co-expressed with NRF2 target genes as well as myeloid genes in bulk RNA seq (Figure 1D-F, Figure 3B-G, Figure 6H, Supplementary figures 3 and 7). Using spatial transcriptomics we found ISGs were induced early by brain MMCs in humans with ICH and remained raised at a similar level in both acute and chronic subgroups, compared with sudden death control patients, whilst Nrf2 target gene induction became more pronounced with time (Figure 3B-E, Figure 6I,J,L,M). ISG induction was correlated with Nrf2 expression (Figure 6K) with both Nrf2 and ISG target gene expression being strongly associated with proximity to the haematoma, but not with microglial homeostatic genes (Figure 3B-E, Figure 6I,J,L,M, Supplementary figure 20). Thus, early perihaematomal ISG induction is a feature of human ICH that is attenuated by NRF2 activation in our mouse model. The presence of an evolving NRF2 response following ICH in patients is clearly insufficient to block ISG induction, given its persistence up to 3 weeks after ICH onset. This suggests that an endogenous stress-activated NRF2 response following ICH may be too little, too late to fully prevent the inflammatory ISG response.

Together, we found that MMC NRF2 is active and protective after ICH in mice, and suppressed type-1 interferon responses by MMCs. Type-1 interferon responses modulate many cell-cell signalling responses to injurious stimuli and could therefore influence the behaviours of other cell types after ICH^41^, suggesting a mechanism by which NRF2 exerts its protection. To investigate the roles of myelomononuclear NRF2 in modulating cell-autonomous and non-cell-autonomous responses to the consequences of ICH we next developed an *in vitro* system.

### *In vitro* stimulation with blood clot conditioned media or lipopolysaccharide recapitulates distinct elements of the NRF2-dependent microglial *in vivo* response to ICH

At a molecular level, *in vivo* ICH exerts its effects both through the release of haematoma-derived molecules such as haem, iron and fibrin breakdown products as well as release of cytokines and other damage associated with activation of pattern recognition receptors^5^. To model these components *in vitro* we established co-cultures comprising primary mouse microglia, human astrocytes and rat neurons, an approach we have used previously to assess transcriptomic changes in specific cell types using *in silico* RNA-seq read separation according to species^13^. We stimulated these co-cultures with mouse blood clot conditioned media (CCM), to mimic the direct effects of haematoma, or LPS, an activator of TLR4 designed to mimic the inflammatory element of the post-ICH environment (Figure 7A).

**Figure 7:**
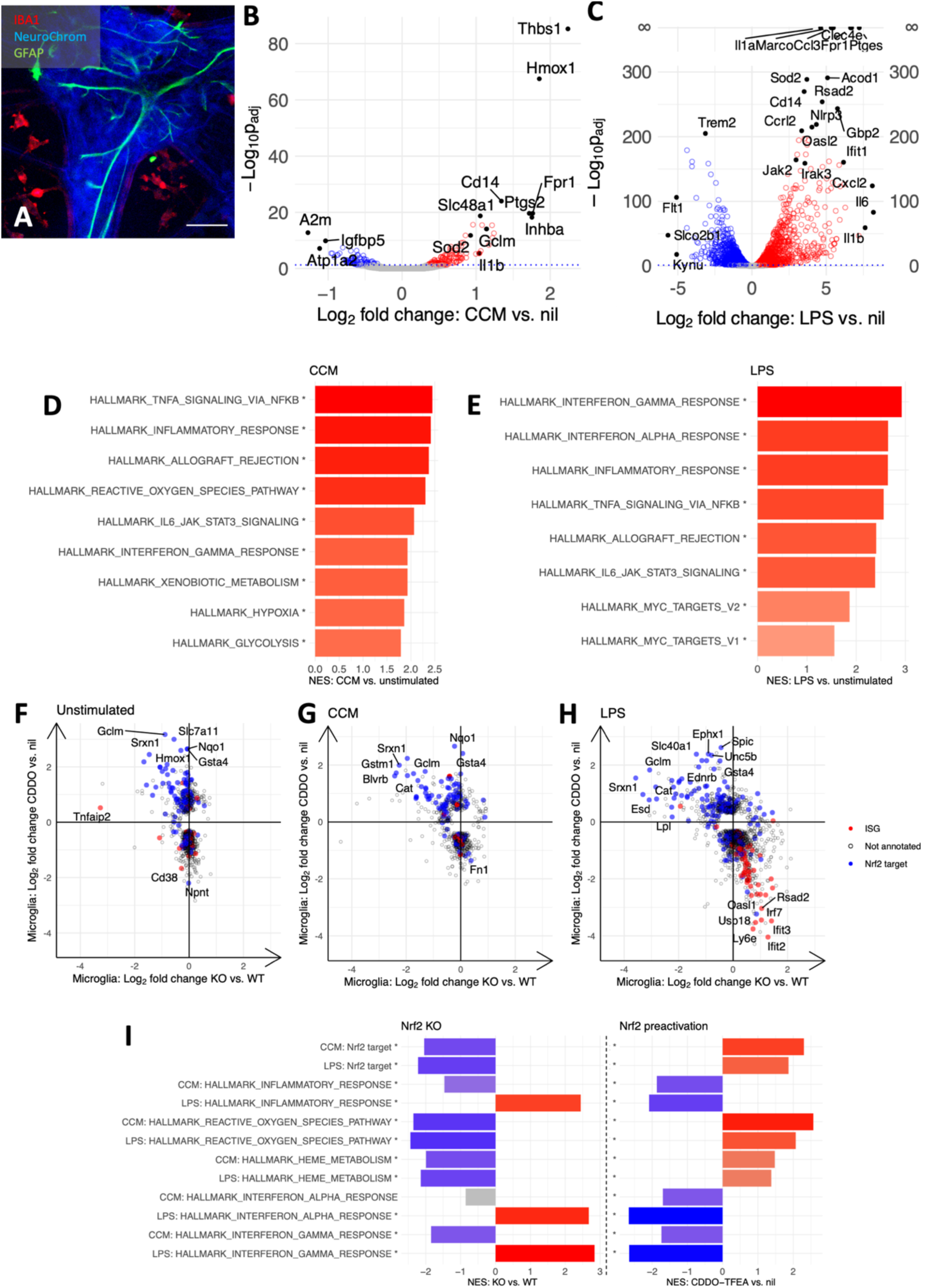
Blood clot conditioned media and lipopolysaccharide exposure *in vitro* recapitulates distinct elements of the *in vivo* microglial responses to ICH, which can be modulated by NRF2 deletion and pharmacological NRF2 activation. A representative immunofluorescence image of an unstimulated co-culture of primary mouse microglia (IBA1; red), rat neurons (NeuroChrom; blue) and human astrocytes (GFAP; green). Scale bar 50µm. **B-C** Volcano plots of the response of WT mouse microglial responses to blood clot conditioned media (CCM; **B**) or lipopolysaccharide (LPS; **C**) exposure *in vitro*. Red points indicate genes with increased expression in the experimental group, blue show genes which are reduced, and grey genes which are not differentially expressed. **D-E** Bar chart of GSEA normalised enrichment score (NES) for significantly enriched or depleted Hallmark gene sets after CCM (**D**) or LPS (**E**) exposure. **F-H** Scatter plots of genes which are differentially expressed by unstimulated (**F**), CCM (**G**) or LPS (**H**) stimulated microglia with NRF2 deletion (x-axis) or NRF2 preactivation using CDDO-TFEA (y-axis). NRF2 target genes described in a previous ChIP sequencing experiment are annotated in blue and Hallmark ISGs in red.^4^. **I** Bar charts of GSEA NES with NRF2 deletion or preactivation using CDDO-TFEA for key Hallmark gene sets and NRF2 target genes defined previously.^4^ Bars representing significantly enriched sets coloured red, sets which are significantly depleted coloured blue and those which are not significantly enriched or depleted are grey. For all analyses, Benjamini-Hochberg adjusted p (p_adj_) <0.05 considered statistically significant. *gene set significantly positively or negatively enriched. WT n=3, KO n=2 independent biological repeats.

Neither CCM nor LPS were cytotoxic and elicited distinct responses in WT microglia (Figure 7B-E, Supplementary figure 21). CCM caused the induction of NRF2 target genes that were also induced by ICH *in vivo* such as *Hmox1, Gclm*, and *Slc48a1*, as well as enrichment of gene sets associated with xenobiotic and reactive oxygen species metabolism (Figures 4F,G and 7B,D). In contrast, LPS caused induction of acute proinflammatory responses including *Il1a, Il6, Ccrl2, Ifit* as well as marked enrichment of ISGs (Figure 7C,E). The induction of ISGs caused by CCM was relatively modest (Figure 7D-H). Thus, with CCM and LPS we were able to, at least in part, mimic the NRF2 and interferon pathway components of the microglial *in vivo* response to ICH.

We next sought to determine how gain- or loss-of-function alterations to microglial NRF2 modulated these responses. Prior to simulation with CCM or LPS, we treated cultures with CDDO-Trifluoethyl Amide (CDDO-TFEA, also known as RTA-404). CDDO-TFEA is a synthetic triterpenoid NRF2-activating drug which is structurally similar to Omaveloxolone (RTA-408), used for the treatment of Friedreich’s Ataxia, a neurodegenerative condition associated with oxidative stress and impaired iron metabolism^42–44^. As expected, CDDO-TFEA treatment increased the expression of NRF2 target genes in WT microglia but had minimal effect on NRF2^-/-^ (KO) microglia (Supplementary Figure 21C-D)^22^. NRF2 pre-activation using CDDO-TFEA and genetic NRF2 deletion had opposing effects on microglial expression of NRF2 target genes in the unstimulated state and after CCM or LPS exposure (Figure 7F-I, Supplementary Figure 21E-J).

We observed elevated ISGs as a result of NRF2 deficiency in the context of LPS stimulation, but not in the context of unstimulated or CCM-stimulated conditions (Figure 7F-H). Therefore, whilst NRF2 suppresses ISG expression under inflammatory conditions (LPS), its loss is not sufficient alone to trigger an ISG response. Further, the lack of an ISG response by CCM-stimulated NRF2 KO microglia indicates that the *in vivo* ISG response is likely to be triggered by a consequence of ICH not attributable to direct actions of clot-released factors on microglia. Rather, NRF2 deficiency may sensitise microglia to ICH-induced interferon pathway stimulators derived from other cell types, such as immune cell production of interferons, and cellular debris, including DNA, released from necrotic cells. Whilst CCM and LPS recapitulate distinct elements of the *in vivo* response to ICH, understanding the significance of the NRF2-activated cytoprotection and ISG-suppressed microglial phenotypes for the wider perihaematomal microenvironment required further investigation.

### Microglial NRF2 activation modulates inflammatory astrocyte responses

Having placed MMC NRF2 as a regulator of antioxidant-haem-catabolic and interferon-stimulated gene expression within MMCs which improves neurological outcomes after ICH (Figure 5I-J, Figure 6B-G), and having also found that NRF2 target gene expression correlated with astrocytic gene expression in human brain tissue after ICH (Figure 1D), we next considered how loss of microglial NRF2 influenced astrocyte responses to CCM and LPS exposure^13^. In wild-type primary co-cultures of human astrocytes, mouse microglia and rat neurons, we found that exposure to CCM and LPS led to the induction of reactive astrocyte transcriptional signatures (Figure 8A-C and Supplementary Figure 22A). Treatment with CCM caused the induction of astrocyte *HMOX1*, enrichment of TNF⍺-NF-𝜅B signalling and MYC associated gene sets, as well as induction of matrix metalloproteinases *TIMP1* and *MMP1* (Figure 8A and Supplementary Figure 22A). Stimulation of astrocytes co-cultured with microglia and neurons with LPS, which acts primarily on astrocytes through its influence on microglia^13^, caused an induction and enrichment of ISGs as well as induction of genes (e.g. *ICAM1* and *VCAM1*) associated with inflammatory astrocyte reactivity (Figure 8B-C). CCM exposure in vitro caused astrocytes to react in a fashion which recapitulated previously curated sets of astrocyte genes induced by in vivo ischaemia by middle cerebral artery occlusion (MCAO), but not in vivo intracerebral LPS injection (Figure 8D).^45,46^ However, LPS exposure induced reactive astrocyte responses that recapitulated both in vivo intracerebral LPS injection and MCAO (Figure 8D).

**Figure 8:**
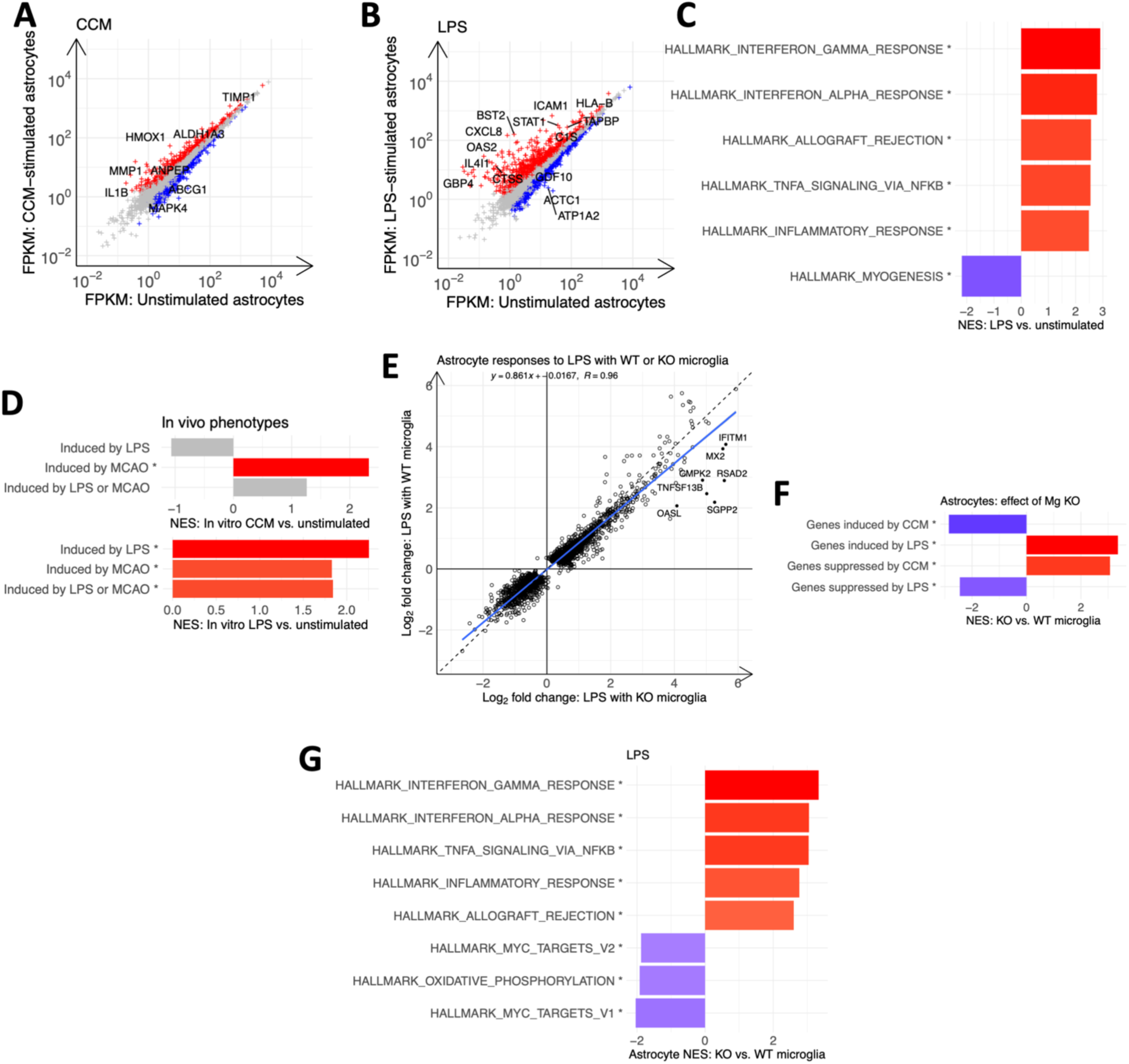
Astrocytes exhibit exaggerated responses to lipopolysaccharide but not blood clot conditioned media when co-cultured with NRF2 deficient microglia and wild-type neurons. A-B. Scatter plots of the response of WT human astrocyte responses to blood clot conditioned media (CCM; **A**) or lipopolysaccharide (LPS; **B**) exposure *in vitro* when co cultured with microglia and neurons. Red points indicate genes with increased expression in the experimental group, blue show genes which are reduced, and grey genes which are not differentially expressed. **C-D** Bar chart of GSEA normalised enrichment score (NES) for the five most highly enriched or depleted Hallmark gene sets after LPS exposure (**C**). and previously curated^45,46^ sets of genes induced by in vivo intracerebral LPS injection and/or middle cerebral artery occlusion (**D**; MCAO; * p_adj_<0.05). **E** Scatter plot of log_2_ fold change in the expression of astrocyte genes which are differentially expressed following exposure to LPS when co-cultured with either WT or KO microglia. Solid blue line indicates fitted linear regression and 95% confidence interval. The formulaic expression of this line and the correlation coefficient are provided. Dashed black line of the form y=x provided for comparison. **F** Bar chart of GSEA NES for sets of genes that are significantly (* p_adj_<0.05) induced or suppressed with CCM or LPS. NES were calculated for astrocytes co-cultured with KO versus WT microglia and stimulated with CCM or LPS, respectively. **G** Bar chart of GSEA NES for the five most highly enriched or depleted (* p_adj_<0.05) Hallmark gene sets in WT astrocytes co-cultured with KO vs. WT microglia after LPS exposure. n=3 independent biological repeats.

Whilst deletion of microglial NRF2 had a minimal effect on the astrocyte response to CCM (Supplementary Figure 22B-C), astrocyte responses to LPS were potentiated when co-cultured with NRF2-deficient KO microglia, compared with WT microglia (Figure 8E-G). This was characterised by enrichment of Hallmark sets of ISGs as well as gene sets associated with TNF⍺-NF-𝜅B signalling (Figure 8G). Examination of sorted astrocytes from mice with ICH revealed concordant results, with increased ISG expression after ICH that was further enriched in NRF2^ΔMMC^ mice (Supplementary Figure 23). Thus, endogenous microglial NRF2 activation status not only influences microglial responses to in vitro LPS and in vivo ICH, but also modulates downstream reactive astrogliosis. In vitro, CDDO-TFEA modulated reactive astrocytosis, with positive enrichment of genes involved in adaptive responses to stressors after both LPS and CCM stimulation and negative enrichment of interferon-stimulated gene sets after LPS stimulation (Supplementary Figure 22D-G). However microglial NRF2 genotype had little effect on CDDO-TFEA’s influence on astrocyte gene expression, indicating that CDDO-TFEA can directly modulate astrocytosis in responses to CCM and LPS through cell-autonomous NRF2 activation (Supplementary figure 22H-K).

### NRF2 activation suppresses ISG expression after ICH, and interferon antagonism restores neurological recovery impaired by MMC NRF2 deficiency

We have determined that deficiency of NRF2 in MMCs causes an exaggerated ISG response to ICH in microglia *in vivo,* and to an inflammatory stimulus *in vitro,* and that patients with ICH also exhibit a marked ISG response. Further, treatment with the NRF2 activator CDDO-TFEA *in vitro* suppressed microglial ISG induction under inflammatory conditions. We therefore determined whether pharmacological targeting of the NRF2 pathway can influence post-ICH ISG induction *in vivo*. We administered CDDO-TFEA by orogastric gavage to mice with ICH and used RT-qPCR to determine how this influenced the expression of ISGs in the perihaematomal ipsilesional brain hemisphere compared with that of mice treated with vehicle only and anatomically-matched contralateral tissue. CDDO-TFEA restrained the ICH-induced expression of *Stat1* in both ipsilesional and contralesional brain hemispheres and ISGs *Ifi27l2a* and *Oas3* in the ipsilesional brain hemisphere in wild-type mice, indicating that the expression of these was antagonised by NRF2 activation (Figure 9A-C)^47^. Reduced post-ICH weight loss and a statistically insignificant trend towards improved sickness score was also observed in mice treated with CDDO-TFEA, suggestive of a potential therapeutic effect (Supplementary Figure 24A-B). We did not observe a significant induction of NRF2 target genes *Hmox1* or *Nqo1* at 24h post-dose (Supplementary Figure 24C-D). We cannot be certain of the mechanism for this. One possibility is a trough effect by which CDDO-TFEA facilitates the catabolism of intracellular haem, which therefore abrogates ICH-induced *Hmox1* expression^5^. Alternatively, since both CDDO-TFEA and ICH-induced stresses can trigger NRF2 activation, it could be that our timepoint chosen reflects the effect of CDDO-TFEA in dampening the consequent ICH-induced stresses that trigger further NRF2 activation.

**Figure 9:**
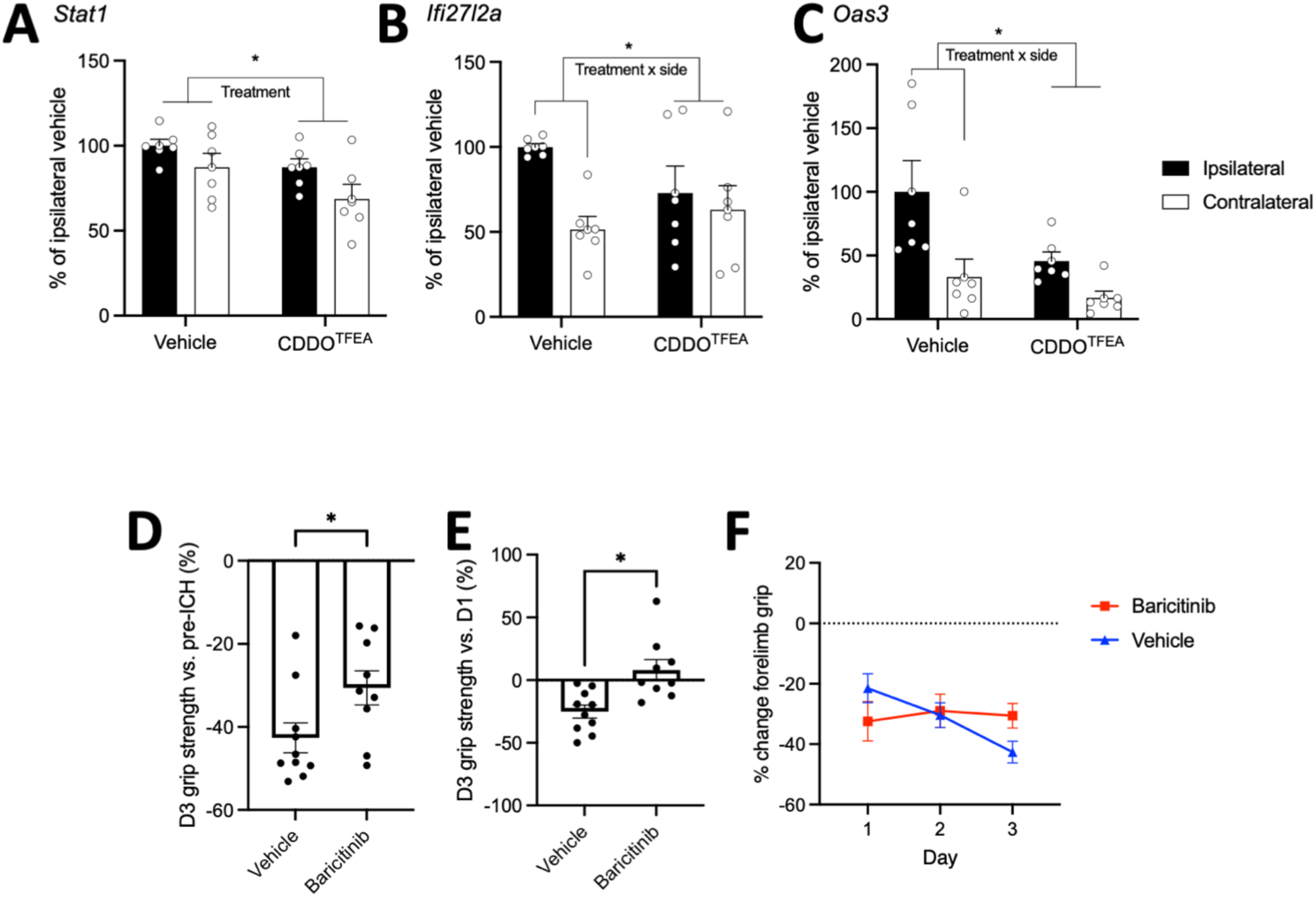
**Pharmacological NRF2 activation suppresses interferon responses and interferon pathway antagonism is protective after ICH *in vivo.*** A-D Bar charts of expression of interferon stimulated genes in brain tissue ipsilateral and contralateral to ICH of WT mice. Samples obtained 24h following simultaneous ICH and enteral administration of CDDO-TFEA or vehicle. Expression of genes were measured by PCR and values normalised to the mean expression in ipsilateral samples of mice treated with vehicle. n=6/group. Two-way repeated measures ANOVA used to determine effects of treatment with CDDO-TFEA versus vehicle (treatment), expression in ipsilateral versus contralateral (side), treatment*side interaction and variance between mice (subject). *Stat1* (**A**; treatment*side F[1-10]=0.1, p=0.71; treatment F[1-10]=7.7, p=0.019; side F[1-10]=4.1, p=0.07; subject F[10- 10]=0.5, p=0.83), *Ifi24l2a* (**B**; treatment*side F[1-10]=5.7, p=0.039; treatment F[1-10]=0.3, p=0.59; side F[1-10]=12.9, p=0.005; subject F[10-10]=3.0, p=0.051) and *Oas3* (**C**; treatment*side F[1-10]=1.3, p=0.28; treatment F[1-10]=7.5, p=0.021; side F[1-10]=8.6, p=0.015; subject F[10-10]=0.6, p=0.77). **D-F** Change in grip strength for NRF2^ΔMMC^ mice with MMC NRF2 deficiency treated with enteral baricitinib or vehicle at ICH onset, day 1 and 2 post- ICH. **D** Grip measured at day three following ICH compared with pre-ICH baseline (Welch’s t-test t=2.2, df=16.4, p=0.041). **E** Grip strength recovery at day 3 compared with day 1 post-ICH (Welch’s t-test t=3.40, df=13.8, p=0.0044). n=10 vehicle, n=9 Baricitinib. Nine NRF2^ΔMMC^ mice were culled following complications of the gavage procedure prior to outcome data collection, and so their data are not presented (n=4 veh, n=5 baricitinib; Supplementary table 4). **F** Daily change in grip strength relative to pre-ICH baseline. All error bars +/- SEM.

As noted above, MMC NRF2 deficiency causes an exaggerated ISG response following ICH and aggravates deficits in ICH grip strength after ICH, suggesting that interferon responses are responsible for this functional deficit. To assess whether direct modulation of interferon signalling is therapeutic after ICH, and understand if interferon signalling mediates the aggravating phenotype in NRF2^ΔMMC^ mice in vivo, we treated NRF2^ΔMMC^ mice with Baricitinib, a small molecule reversible Jak inhibitor that is used for the treatment of rheumatoid arthritis and Covid-19^48^ and can suppress ISG responses.^49^ In NRF2^ΔMMC^ mice, Baricitinib improved functional outcomes by restoring early recovery of grip strength between days one to three post-ICH (Figure 9D-F). Sickness scores were also improved (Supplementary Figure 24E-K).

Together, these findings demonstrate that NRF2 activation suppresses ISG expression and that antagonism of the ISG response which is caused by MMC NRF2 deficiency in the preclinical ICH model, and is observed in patients with ICH, improves early recovery of neurological deficits (Figure 10).

**Figure 10:**
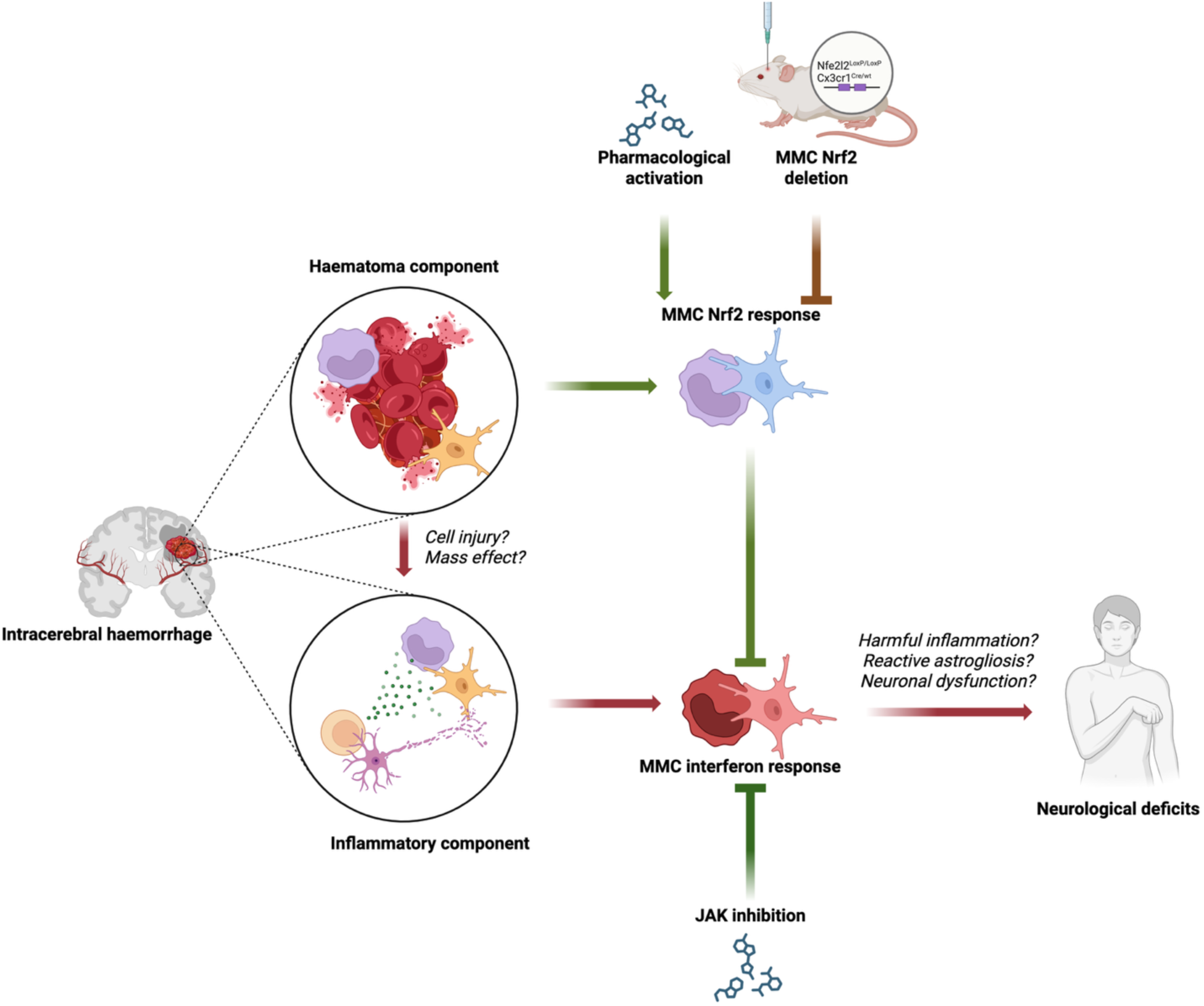
Proposed model for the influence of MMC NRF2 on neurological outcome after ICH. After ICH, factors released by the haematoma (e.g. iron, haem, reactive oxygen species) and/or environmental factors (e.g. ischaemia) cause activation of Nrf2 in MMCs. This can be augmented or suppressed by pharmacological activation with CDDO-TFEA, or cre-mediated deletion of functional Nrf2, respectively. Simultaneously, ICH evokes an inflammatory response which includes induction of an interferon-stimulated response, potentially secondary to cellular injury and/or mass effect. The MMC interferon-stimulated response contributes to neurological deficits, potentially through injurious downstream inflammatory MMC or reactive astrogliotic responses. MMC Nrf2 activation serves to limit the harmful MMC interferon response but does so incompletely in humans with ICH. Direct pharmacological JAK inhibition, upstream of the interferon response can rescue harms associated with excessive MMC interferon responses after ICH, allowing for improved recovery of neurological deficits.

## DISCUSSION

Following ICH, a complex series of events takes place in the brain that ultimately defines the clinical outcome, be it recovery, long-lasting deficits, or death. Myeloid immune cell responses concentrated in the perihematomal area are increasingly implicated in haematoma evolution and clinical outcome^9,11,50^. Understanding these events can point to potential therapeutic strategies to improve outcomes. Our study has shown that patients who have died after ICH exhibit gene expression profiles associated with both deleterious interferon pathway activation as well as adaptive-protective NRF2-driven responses converging on MMCs with both cell-autonomous and non-cell autonomous consequences. Moreover, we show that manipulating these responses after ICH, both through MMC-targeted endogenous NRF2 deletion and translationally relevant pharmacologic NRF2 activation, or suppression of interferon responses, has the potential to alter outcomes. We propose that NRF2 activation in response to ICH serves to limit the development of a harmful interferon-stimulated MMC phenotype (Figure 10).

NRF2 is a widely expressed transcription factor whose role is context- and cell type-dependent^5^. NRF2 regulates gene sets including antioxidant, conjugation detoxification, proteostasis and those mediating certain metabolic pathways^22,51–53^. NRF2 activation is also thought to be anti-inflammatory in nature, particularly in macrophages^27,54^. One key mechanism is to provide greater redox buffering capacity to limit reactive oxygen species-induced pro-inflammatory pathway activation^55^. The induction of the prototypical NRF2 target gene *HMOX1*, primarily localised to perihaematomal MMCs in human post-mortem tissue is significant as, along with the associated spatial and deep transcriptome profiling analyses, it points to an adaptive-protective response which evolves over the first eight days after ICH (the median of our subacute group), persists for at least 21 days (the median of our chronic group) and is localised to an area crucial to haematoma resolution^9^. After ICH, excess iron, immune cell activation, and/or reactive oxygen species production as a consequence of perturbed metabolism can activate NRF2^5^. We show that activation of NRF2 in MMCs occurs after ICH in mice as in humans and also following exposure of microglia to clot-conditioned medium *in vitro*. Significantly, mice which lack functional NRF2 in MMCs display worse outcomes after ICH, placing microglial/macrophage NRF2 as a functionally important pathway. However, despite the evidence that the MMC NRF2 pathway is activated and is both cytoprotective and anti-inflammatory in the preclinical ICH model, the endogenous response evident in humans after ICH may be too little, too late to prevent significant secondary injury which is associated with early ISG induction. This supports the rationale to augment NRF2 activity beyond endogenous ICH-induced levels through pharmacological means^44,56^. An analogous situation exists in the response of astrocytes in neurodegenerative disease models of ß-amyloidopathy and tauopathy, where endogenous NRF2 responses are also observed, but are insufficient to prevent neuronal and synapse loss^46^. However, artificial over-activation of NRF2 in astrocytes strongly attenuates proteinopathy and downstream patho-progression and cognitive deficits^46^. In the case of ICH, we propose that pharmacological activators of NRF2 after ICH would enhance MMC NRF2 activity more quickly and strongly than that which occurs endogenously, with a beneficial effect. If instituted within 24h of ICH onset, pharmacologically enhanced NRF2 activation might reprogramme the window of transcriptional responses to early brain injury that we observed in the first four days of ICH onset and aligns with others’ findings of a four-day window of transcriptional activation in haematoma-derived immune cells that is associated with clinical outcomes^8,9^.

Beyond direct haem detoxification, the mechanism(s) by which NRF2 activation in microglia, macrophages and other MMCs can alter the events after ICH are likely several-fold. One pathway influenced by NRF2 status is ISG expression. ISG responses are observed after ICH in patients and in mice. This may reflect a direct response to interferon secretion but could also be elicited after ICH by excessive exposure to necrotic cell-derived double-stranded DNA and consequent activation of cGAS-STING^57^. One would expect this stimulus to be maximal during the first 24-72h following ICH, during which period it has been previously noted that terminal deoxynucleotidyl transferase dUTP nick end labelling (TUNEL) is maximal.^8^ It is notable that *in vitro* microglial exposure to LPS but not CCM induced an ISG response, reinforcing that MMC NRF2 deficiency *in vivo* likely sensitises the brain to secondary inflammatory stimuli induced by ICH but not haematoma products directly. Furthermore, in vitro, we find that microglial NRF2 deficiency causes an exaggerated interferon-stimulated phenotype in both microglia and astrocytes after an inflammatory insult. Pharmacological NRF2 activation with CDDO-TFEA reduced ISG responses in cell culture as well as in our in vivo mouse ICH model. Moreover, the deficits observed in mice lacking NRF2 in MMCs are rescued by Baricitinib, an inhibitor of JAK which transduces interferon signals. Collectively this points to a scenario whereby NRF2 status in MMCs modulates cell-intrinsic and extrinsic ISG responses, key determinants of adverse outcomes.

Our study suggests that a potential therapeutic approach could involve modulation of the NRF2 and/or interferon signalling pathways post-ICH. Baricitinib has an established clinical profile for the treatment of rheumatoid arthritis and Covid-19^48,58,59^ and we found it improved recovery of neurological deficits in mice lacking NRF2 in MMCs. Moreover, our work suggests that boosting NRF2 activity may also be beneficial via the suppression of interferon signalling. This is particularly relevant for the majority of patients with ICH who are of older age and thus are predisposed to senescence-associated priming of MMCs, exaggerated ISG responses, and reduced NRF2 potency^5,60–62^. We showed that CDDO-TFEA reduced ISG responses *in vitro* and *in vivo*, raising the possibility that targeting NRF2 as well as interferon signalling may be an optimal strategy by targeting post-ICH events in complementary ways. CDDO-TFEA (RTA-404) is a close structural relative of RTA-408 (Omaveloxolone, Skyclarys), a drug recently approved for treatment of Friedreich’s ataxia, a condition also associated with intracellular iron accumulation^42,43^. Baricitinib and/or Skyclarys may help the process of inflammatory resolution after ICH and promote recovery via other ways such as enhanced redox buffering. The therapeutic efficacy of these drugs as monotherapy, and in combination, could be assessed further in a preclinical in vivo setting with post-ICH dosing. These experiments would compare dosing frequency, duration and route of administration and test consistency of results with an alternative mouse ICH model, such as one using autologous blood injection.^32^ Such a study could also provide the opportunity to validate findings of our mass spectrometry imaging analysis in a targeted fashion. We have demonstrated a reduction in motor deficit, however future studies could consider alternative outcomes including cognitive testing and co-ordination and investigate changes in neuronal function directly using neurophysiological approaches and blood brain barrier function or oedema using magnetic resonance imaging and analysis of transepithelial electrical resistance. Further in vitro testing could also be undertaken in a fully human system.

To conclude, it is generally acknowledged that there is a therapeutic window, perhaps days-weeks in duration, within which secondary injury after ICH might be influenced and that both brain-resident microglia and infiltrating MMCs can shape the ultimate outcome. Our study places NRF2 and interferon signalling, specifically within mononuclear myeloid cells, as mechanistically linked influencers of neurological consequences of ICH, both of which are targetable by compounds currently in clinical use.

## METHODS

### Regulatory approval

The NHS Lothian Research Ethics Committee (LREC 2003/8/27), East of Scotland Research Ethics Service (16/ES/0084), Scotland A Research Ethics Committee (10/MRE00/23) and the North East Newcastle & North Tyneside 1 Research Ethics Committee (09/H0906/52+5) granted approval for studies on human patient-derived tissue. Written consent was provided by participants, or their legal representative if participants lacked mental capacity, in accordance with the Human Tissue (Scotland) Act 2006 and the Adults with Incapacity (Scotland) Act 2000.

We conducted all animal experiments in accordance with the UK Animals (Scientific Procedures) Act 1986, under UK Home Office institutional and project licences with local ethical and veterinary approval (Biomedical Research Resources, University of Edinburgh). All investigators conducting licenced procedures held valid UK Home Office personal licences.

### Case-control neuropathological studies

#### Participants

We included patients with first ever spontaneous cerebral-small vessel disease associated ICH which occurred between June 2010 - 31 May 2016 (cohorts 1 and 3), or April 2018 – November 2021 (cohort 2) who were enrolled in the Lothian IntraCerebral Haemorrhage, Pathology, Imaging and Neurological Outcome (LINCHPIN) study, which is a prospective, community-based inception ICH cohort^63^. These were derived from distinct sub-studies and as such inclusion and exclusion criteria for each cohort are described in Supplementary Figures 1,2 and 4. Cohort 4 patients were using the same criteria as cohort 3 with the removal of time constraints and the additional criterion that haematoma margin must be visible in the unsectioned embedded block of brain tissue.

#### Neuropathological assessment

A senior consultant neuropathologist (CS) conducted post-mortem examinations and brain tissue sampling, and undertook histological assessment with small vessel disease grading, Braak stage, Thal phase and CAA burden assessment^64–66^. For cohort 1, two ipsilateral specimens were selected from archived tissue blocks per patient whilst for cohort 2, one ipsilateral perihaematomal specimen and one contralateral anatomically-matched sample were prospectively obtained. These included tissue which was immediately adjacent, within 1cm, to the edge of haematoma, but without macroscopic haematoma present. Data for all other clinical variables were extracted from patient medical records and radiological images^67,68^.

#### RNA extraction, purification, and sequencing

We performed RNA extraction and purification using the RNeasy Lipid Tissue Mini Kit (Qiagen, 74804). RNA integrity was assessed by RNA electrophoresis (RNA 6000 Nano Assay, Agilent 5067-1511; Bioanalyzer Agilent, 2100). Pooled barcoded TruSeq stranded mRNA-seq libraries (Illumina) were prepared according to manufacturer’s instructions and sequenced in a S1 flow cell to a depth of approximately 50 million 50 base pair (bp) paired-end reads per sample using a NovaSeq 6000 sequencer (Illumina). We aligned reads to the Human Genome using STAR (version 2.5.3a), determined per-gene read counts using featureCounts (version 1.5.2), normalised gene expression to gene length using fragments per kilobase of transcript per million mapped reads (FPKM) and determined differential gene expression using DESeq2 (version 1.18.1) with a Benjamini-Hochberg adjusted p-value (p_adj_) threshold of 0.05 and quality surrogate variable analysis (qSVA) to correct for sample quality bias^69–72^. For cohort 1, we compared mean read counts per gene for the two samples of each individual patient for against the read counts for each control sample.

#### Gene co-expression network analysis

We used Graphia (v2.1) to produce a gene coexpression network (GCN) of all genes with a non-zero FPKM value in any sample^73^. This graphically represents a gene-to-gene Pearson correlation matrix with an edge threshold of Pearson R>0.964. We used Pearson R ranked 𝜅-nearest neighbours edge reduction (𝜅=5) and Markov Clustering (MCL) with an inflation value of 1.25 to produce large clusters that adhered to GCN topology^73–76^. We next annotated genes specifically (>5 fold expressed by one cell type versus any other cell type) expressed by brain myeloid cells, neurons, astrocytes, endothelial cells, and oligodendrocytes using a previously published RNA sequencing dataset of immunopanned human brain cells^28^. We defined genes as NRF2-regulated if, in a previously published dataset, they were differentially expressed in either NRF2-/- or Keap1-/- mouse embryonic fibroblasts (MEFs), compared with wild-type MEFs, and identified as containing an NRF2 binding region by chromatin immunoprecipitation sequencing^22^. Finally, we tested for enrichment of all clusters for cell type or NRF2-regulation status annotations in Graphia using Fisher’s exact test with enrichment threshold of adjusted p-value (p_adj_)<0.05.

#### Dual in-situ RNA hybridisation-immunohistochemistry

We performed Dual in-situ RNA hybridisation-immunohistochemistry (ISRH-IHC) on 4µm thick brain tissue sections obtained at autopsy of patients included to cohort 3. We first stained sections from all patients for haem oxygenase 1 mRNA (*HMOX1*) by Fast Red ISRH and then counterstained for IBA1 using 3,3′-diaminobenzidine (DAB). Six perihaematomal sections with strong ISRH signal were then selected for staining with GFAP or Olig2 by IHC after ISRH. ISRH was performed using BaseScope™ Detection Reagent Kit v2 – Red (ACDbio, 323900) according to manufacturer’s instructions. For DAB staining, the Novolink Max Polymer Detection System (Leica, RE7280-K) was used with the following antibodies: 1:3000 IBA1 Abcam, ab178846; 1:100 OLIG2 (Abcam, ab109186); GFAP 1:800 (Dako, Z0334).

For each stained section and negative control, we acquired ten non-overlapping field of view (FOV) brightfield images using an Axio Imager D2 (Zeiss). Images were automatically processed using Fiji (version 2.1.0/1.53c)^77^ to determine the proportion of each cell that was stained by DAB or Fast Red (Supplementary Methods 1). Cells were labelled as DAB positive if the proportion of their area detected as DAB+ was greater than the 95^th^ percentile of the equivalent measurement made in DAB negative control sections. We then calculated total Fast Red (*HMOX1*) and DAB (IBA1, OLIG2 or GFAP) stained area as well as total *HMOX1* area in DAB positive cells and DAB negative regions in ten FOVs.

#### CosMx Spatial transcriptomics

From potentially eligible patients, we selected four ICH patients who died <3d (acute), or >12 days (chronic) and 4 controls who died suddenly of non-neurological causes. 7mm formalin fixed paraffin-embedded brain tissue cores at the haematoma margin were sectioned to 5µm thickness and mounted on two slides with two patients from each group per slide, and processed according to the CosMx™ Spatial Molecular Imaging (SMI) manufacturer’s instructions for FFPE RNA assays (Bruker/NanoString). For FFPE human brain tissue, manufacturer-recommended modifications were applied. These included an additional 70% ethanol wash for 2 hours to dehydrate the tissue after target retrieval, tissue permeabilisation by proteinase K digestion at 3 µg/mL in 1X PBS containing 0.5% Tween-20 for 30 min at 40 C, and preparation of fiducials at 0.0005%.

Spatial transcriptomic profiling was performed using the CosMx Human 6K Discovery Panel, and the tissue was stained with the CosMx Human Neuroscience Cell Segmentation Kit (DAPI, rRNA, Neuro Histone and GFAP) and CD68. Additionally, the TrueVIEW™ autofluorescence quencher (Vector Laboratories) was applied to the stained slides according to the manufacturer’s instructions to reduce background fluorescence.

Images of 627 fields of view (0.51 × 0.51 mm per field), across the two slides were acquired on the CosMx™ SMI instrument using the manufacturer’s recommended settings for Human Brain tissue. Raw image data were processed and decoded using AtoMx™ Spatial Informatics Platform (v2.2.1) with the default Brain Tissue (RNA) setting. Cells were segmented based on nuclear and membrane markers provided in the segmentation kit, and probe counts were assigned to segmented cells using the CosMx analysis pipeline. Haematoma regions were segmented using QuPath (v.0.2.3).

### *In vivo* mouse experiments

#### Mouse lines

We used five genotypes of mice, all of which were crossed to a C57Bl/6J background. Wild-type (WT; C57Bl/6J-*Cx3cr1*^+/+^:*Nfe2l2*^wt/wt^; Charles River ^78^® Strain), Nfe2l2 floxed mice with loxP sites flanking Nfe2l2-exon 5 which encodes the DNA binding domain of NRF2 (NRF2^LoxP^; C57Bl/6J-*Cx3cr1*^+/+^:*Nfe2l2^f^*^l/fl^; JAX 025433) and Cx3cr1-cre (Cre; C57Bl/6J-*Cx3cr1*^Cre/+^:*Nfe2l2^wt^*^/wt^; JAX 025524) were purchased. Knock out (NRF2^ΔMMC^) mice lacking functional NRF2 in MMCs (C57Bl/6J-*Cx3cr1*^Cre/+^:*Nfe2l2^f^*^l/fl^) were generated by crossing NRF2^LoxP^ and Cre mice.

#### Stereotactic striatal injection of bacterial collagenase

We induced intracerebral haemorrhage using stereotactic injection of bacterial collagenase in mature male adult mice (conditional knock-out studies median 33 weeks age [range 24-44 weeks]; pharmacological studies median 19 weeks age [range 12-30 weeks]). All mice were anaesthetised using inhaled isofluroane and nitrous oxide in oxygen and received adjuvant subcutaneous buprenorphine (50μg/kg; Vetergesic, CEVA). Whilst under anaesthesia, we used a core body temperature-regulated homeothermic pad to ensure stable normal temperature during surgery. Under strict asepsis, a single burrhole was placed at 2mm lateral, 2.5mm deep to bregma. 0.5µL 0.09 units/μLtype VII bacterial collagenase (Sigma, C2399) or 0.5µL vehicle sterile 0.9% sodium chloride solution was injected through a glass micropipette at 1µL/min the micropipette left *in situ* for 10 minutes to minimise reflux. Solely for baseline experiments to establish background and cre-dependent gene expression in NRF2^LoxP^, NRF2^ΔMMC^ and Cx3cr1^Cre^ lines in the sham state, we used mice from an independent experiment that received bilateral striatal injections of 0.9% sodium chloride solution using a Hamilton syringe (VWR). Mice used for comparison of ICH with sham states or to determine NRF2-dependent gene expression after ICH underwent unilateral injection with a micropipette. The incision was closed and 5% lignocaine (EMLA, AstraZeneca) topical anaesthetic applied. During recovery from anaesthesia, animals were placed in a cage with both warm and cool zones to allow for self-regulation of body temperature. Once fully recovered they were returned to individually ventilated cages, provided with wet and dry feed (Special Diets Services, 801010) as well as water *ad libitum* for 24 or 72h until culled by transcardial perfusion (Histology - 0.9% sodium chloride solution followed by 4% paraformaldehyde; RT-qPCR – Diethyl pyrocarbonate treated 0.9% sodium chloride solution) or overdose of anaesethetic (RNA-sequencing; Sodium pentobarbital; Pentoject). For pharmacological studies we administered 10mg/kg 2-cyano-3,12-dioxooleana-1,9-dien28-oic acid trifluoroethylamide (CDDO-TFEA) immediately prior to induction of anaesthesia or 10mg/kg Baricitinib immediately prior to induction of anesthesia as well as at 24h and 48h post-surgery^44,79^. At all time points a vehicle only (Corn oil with 10% dimethylsulfoxide) control group of mice was used. For all experiments, the order of surgery, assignment to surgical or sham groups, and pharmacological treatment were randomised (https://www.random.org/lists/). The research team and technical staff were masked to assignment and mouse genotype until study closure.

#### Sample size

For experiments where the primary outcome was forelimb grip strength we used pilot data to determine that for 80% power to detect a 30% difference in percentage change in grip strength from −33% with alpha of 0.05 and standard deviation of 8.1%, we require 11 mice per condition. Where gene expression was the primary outcome, we predicted a fold-change in CD36 of 2.69, a coefficient of variation of 55% and counts per million reads mapped of 38.^80,81^ For a false discovery rate <10% and alpha 0.05 we would require 6 mice per condition. For each of these analyses we included an additional 1 mouse per every 5 mice used to allow an attrition rate of 20%. Numbers used per analysis, exclusions and reasons for exclusion are described in Supplementary table 4.

#### Animal monitoring and behavioural outcomes

We monitored post-operative welfare by twice daily sickness scoring according to the IMPROVE guidance and daily weighing of mice^82^. All assessors were masked to individual mouse genotype, haemorrhage status and treatment allocation. Forelimb grip strength was our prespecified primary outcome. It measured (Bioseb BIO-GS3) pre-operatively, at 24, 48 and 72h post-operatively. Percentage change in grip strength from baseline and recovery from 24h to 72h were calculated. The Clark neurological deficit score was calculated daily post-operatively^83^.

#### Histology

Following transcardial perfusion-fixation, we dissected mouse brains from the cranial vault and post-fixed in 4% paraformaldehyde and then 20% sucrose for a further 24h each. We snap froze brains in isopentane and stored them at −20°C before taking 20µm coronal sections using a cryostat (Leica). We performed H&E staining using Harris haematoxylin (Epredia Shandon, 6765004) and aqueous eosin (VWR, 341973R). For haematoma volume quantification we applied nickel-enhanced 3,3′-diaminobenzidine (DAB) solution (Vector, SK-4100) to sequential brain sections at 400µm increments and counterstained with haematoxylin. We imaged sections using an Axioscan (Zeiss) slide scanner, measured the total DAB stained area (µm^2^) for all sections using QuPath (v.0.2.3) and multiplied this by 4x10^-7^ to derive the haematoma volume in µL^84^.

We performed tyramide signal amplification immunofluorescence staining after antigen retrieval in a 97.5°C solution of 10mM tris, 1mM EDTA, pH 8.6 for 20 mintues. We incubated sections in a blocking solution with 5% normal goat serum (NGS; Vector, S-1000) at room temperature, then overnight with anti-Iba1 antibody (1:500; Wako, 019-19741), 0.3% triton X-100 (Sigma, X100) and 2.5% NGS. We quenched peroxidases with 3% hydrogen peroxide and then applied biotinylated goat anti-rabbit IgG (1:200, Vector, BA-1000) in 0.1% tween 20 tris-buffered saline (TBST) for 1.5h. To complete the IBA1 stain, we used the Alexa Fluor 647 Tyramide SuperBoost^TM^ (Invitrogen, B40936) kit according to manufacturer’s instructions. Next, keeping slides protected from light we repeated these processes including antigen retrieval to strip the primary and secondary antibodies whilst retaining the tyramide signal. For the second stain, we used a further streptavidin and biotin block (Invitrogen, R37628) prior to applying the second primary biotinylated antibody. For the second secondary antibody we used either rabbit anti-P2Y12 (AnaSpec, ANA55043A) or rabbit anti-HMOX1 (Enzo, ADI-SPA-895) at 1:500 and stained with Alexa Fluor 555-conjugated tyramide (B40933). We dried our slides and mounted with fluorescence mounting medium (Dako, S302380-2). For all stains, we included fluorescence minus-one (FMO) negative controls with one primary antibody omitted. All staining runs were completed in a single batch.

For fluorescence stain quantification we used an Axioscan (Zeiss) slide scanner to acquire an image of one section per mouse at the level of the haematoma. Images were obtained for all stained channels and the fluorescein isothiocyanate (FITC) channel to measure tissue autofluorescence. Using QuPath (v.0.2.3), we defined the haematoma as a highly autofluorescent region in the FITC channel and automatically defined perihaematomal regions of interest (Figure 4B). Next, we quantified HMOX1 and P2Y12 staining in IBA1 stained areas using QuPath positive cell detection. First we defined Iba1 stained regions as those with a mean staining threshold above the noise level detected in Iba1 unstained FMO controls. The threshold for P2Y12 or HMOX1 staining positivity was defined as the 95th percentile for stain intensity in Iba1 stained detections of P2Y12 or Hmox1 negative FMO controls. For comparison of groups, the proportion of cells that were at or above the 75th percentile in sham mice of NRF2^LoxP^ and Cre genotypes was defined as “Hi” staining. For illustrative purposes, we acquired high resolution images of representative sections using a Nikon A1R confocal resonant scanning microscope and produced maximum intensity projection images using Fiji (v.2.1.0/1.53c)^77^.

For DAB staining of NeuN, we applied rabbit anti-NeuN 1:4000 (Abcam, AB177487) in 2.5% anti-goat blocking solution to 20µm thick coronal brain sections overnight at 4°C at the level of haematoma following blocking and quenching as above. Secondary biotinylated goat anti-rabbit IgG (Vector, BA-1000) and then avidin-biotinylated peroxidase complex (Vector PK-6100) were applied for 1h at room temperature before incubation with DAB solution (Vector, SK-4100), H&E staining and coverslipping as above. We used brightfield microscopy to obtain high magnification images in cortical, perihaematomal and haematoma regions of interest and manually quantified normal, atypical and pyknotic NeuN positive cell body morphologies, blind to region of interest and genotype.

For mass spectrometry imaging, we lifted archived 20µm thick coronal brain sections at the level of haematoma from SuperFrost Plus (Epredia, 10149870) slides by incubating in a bath of pH 9 Tris-EDTA at 70°C for 1h to remove OCT and allow facilitate careful lifting of the tissue using a scalpel in a room temperature waterbath. Tissue sections were floated and then mounted on Matrix-Assisted Laser Deorption/Ionisation (MALDI) IntelliSlides (Bruker, 1868957), dried at room temperature and then frozen at −20C before processing. Tissue from five randomly allocated mice were mounted per slide. These were processed using a HTX M5+ sprayer (HTX Technologies, LLC). Firstly tissue sections were digested through the spraying of Trypsin using the default trypsin settings within the sprayer. We allowed tissue digestion to occur by placing slides in a sealed dish containing 50 mM TEAB (Triethylammonium bicarbonate) at 37 degrees overnight. We then applied a CHCA (α-Cyano-4-hydroxycinnamic acid) matrix in a 50% Acetonitrile, 0.1% Trifluoroacetic acid buffer using the appropriate preset in the sprayer software. We next collected data about peptide abundance on haematoma and perihaematomal tissue regions within these slides in a Bruker Rapiflex Tissue Typer mass spectrometer at 50um spatial resolution within the mass/charge (m/z) range of 500-1800.

#### Fluorescence activated cell sorting (FACS)

We dissected brain tissue of mice killed by overdose of anaesthetic and placed brains in ice cold Dulbecco’s phosphate buffered saline (DPBS; Invitrogen, 14287080). We removed the cerebellum and olfactory bulbs before performing mechanical and enzymatic dissociation using the gentleMACS Octo Dissociator, according to manufacturer’s instructions (Miltenyi Biotech, 130-096-427). We removed debris and myelin using a cell strainer and gradient centrifugation (Miltenyi, 130-110-916). We applied red blood cell lysis solution (Miltenyi, 130-094-183). We estimated live cells isolated per brain using trypan blue exclusion (Sigma, T8154) and a haemocytometer. Next, we applied 1:100 rat anti-mouse Cd16/32 (BioLegend, 101302), Cd11b-FITC (1:200; BioLegend, 101206), Cd45-Pacific blue (1:50; BioLegend, 103126), ASCA-2-APC (1:200; Miltenyi, 130-116-245), Ly6g-PE (1:1000, BioLegend, 127607), O4-PE (1:200; Miltenyi, 130-117-357) and Draq7 (1:200; Invitrogen, D15105). FMO samples were also prepared. We isolated Microglia, MdCs and astrocytes using a FACSAria Il flow cytometer (Beckton Dickinson Immunocytometry Systems [BD]) on Purity precision, running BD FACSDiva (v6.13; Supplementary Figure 5). We determined the fraction of each cell type per sample using FCS Express 7 Research (v7.04.0014) and calculated the absolute number of each cell type per brain by taking the product of this and the total number of cells estimated earlier using a haemocytometer.

#### RNA isolation and sequencing

We isolated RNA from sorted cells immediately following sorting using RNeasy Plus Micro kits (Qiagen, 74034). We produced cDNA for sequencing using the SMART-Seq V4 PLUS ultra-low RNA input kit (Clontech) and sequencing libraries using Nextera XT DNA library Preparation kit (Illumina). These were sequenced using the NextSeq system (Illumina) in three 75 cycle high output runs to give 20-25 million single-end reads per sample. Read alignment, counts, generation of FPKM and differential expression analysis were then undertaken as described above, but without qSVA.

#### Reverse transcription quantitative polymerase chain reaction (RT-qPCR)

For RT-qPCR, we synthesised cDNA from RNA isolated as above using the Transcriptor First Strand cDNA synthesis Kit (Roche, 04897030001), according to manufacturer’s instructions. We performed qPCR using Power SYBR Green (Applied Biosystems, 4368708) on the Mx3000p qPCR system (Agillent) using custom primer DNA oligonucleotide (Sigma, VC00021N) sequences described in Table 1. We used mean Ct value for two technical repeats to calculate fold change in gene expression using the 2^-ΔΔCt^ method, normalised to *Rpl13a* expression and a control reference sample^85^.

**Table 1.**
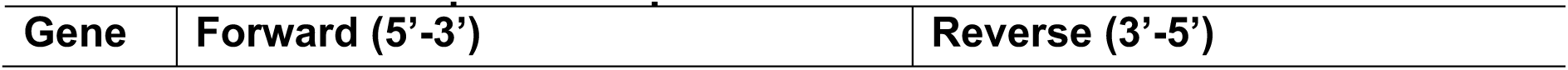
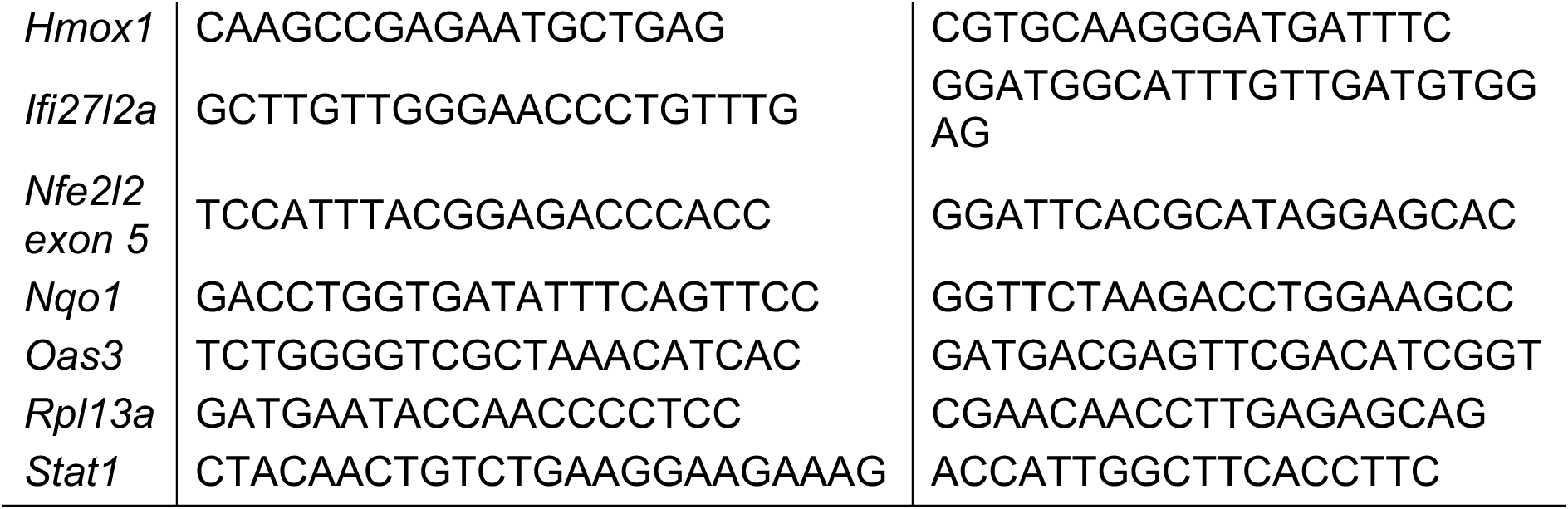
In vivo mouse primer sequences.

### In vitro analyses

We established all cell cultures in sterile conditions under a laminar flow cell culture hood and all reagents were passed through a 0.22μm filter prior to use. The preparation of all reagents has been previously described in detail^13^. Mixed sex cells were used for all in vitro analyses.

#### Mouse cortical mixed primary glial culture

We used Postnatal (P0-P4) mouse pups of C57BL/6J background that were either NRF2 wild type (Nfe2l2^+/+^; WT) or with global NRF2 deletion (Nfe2l2^-/-^; KO; JAX 017009, supplied by Professor Masayuki Yamamoto). These were decapitated and brain tissue decapitated in ice cold dissection media (DM+K; sodium sulfate, potassium sulfate, magnesium chloride, calcium chloride, HEPES, phenol red, D-(+)-glucose, kynurenic acid). We used 20 mins papain digestion (papain 36,000 USP units/mL [Merck Millipore,5125]) at 37°C before washing and triturating in DMEM solution (Dulbecco’s modified Eagle’s medium [Gibco, 31966-021], 10% foetal bovine serum [Invitrogen, 10108-165], 1x antibiotic-antimycotic [streptomycin, penicillin, amphotericin B; Thermo Fisher, 15240062]). We plated this cell suspension onto poly-D-lysine (BD Bioscience, 354210) coated T175 flasks and placed these in a humidified incubator (37°C, 5% CO2, 20% O2) for 2.5h before replacing the media with fresh DMEM solution and allowing growth to confluency for 12-14 days. We cultured tissue from WT and KO mice separately.

#### Human primary astrocyte culture

We thawed frozen primary human astrocytes (-190oC; Caltag Medsystems, SC-1800) and plated them in in poly-D-lysine-coated T175 flasks. They were incubated with human astrocyte medium (Caltag Medsystems, SC-1801) for a maximum of 7 passages using Trypsin EDTA (Sigma, T3924). For experiments, these were seeded in 24-well tissue culture plates (Greiner, 662160) at 3-5x105 cells per well and left for 3 days until 80-90% confluency was achieved.

#### Rat primary neuronal culture

We dissected the cortices of Sprague Dawley rat embryos of embryonic day 20.5 on ice cold DM+K media and removed the hippocampi. We performed enzymatic digestion at 37°C using papain for 40 minutes, with papain solution changed midway. We then washed the cortices twice with 37°C DM+K and then twice more with 37°C 1% neurobasal-A (neurobasal-A-medium [NBA; Gibco, 10888-022]; 1% v/v rat serum [Harlan SeraLab, R-0100B], 1x antibiotic-antimycotic, 2% v/v B-27 supplement [Invitrogen, 17504-044], 1mM L-glutamine) before triturating in 1% NBA. We diluted this cell suspension in 37oC OptiMEM I+ (OptiMEM I [Gibco, 31985-047], 0.36% w/v D-[+]-glucose, 1x antibiotic-antimycotic) to 14mL per cortical hemisphere used and plated 0.5ml on top of each well containing human primary astrocytes, having first aspirated the human astrocyte medium. After 2.5h, we aspirated the OptiMEM I+ and debris and replaced with 1mL fresh 37°C 1% NBA with 4.8μM Cytosine β-D-arabinofuranoside hydrochloride (AraC; Sigma, C6645). These co-cultures of human astrocytes and rat neurons were incubated until DIV2 whereupon we further 1% NBA with AraC to each well. At DIV 4, we exchanged 1mL media for 0% NBA (NBA, 1x antibiotic-antimycotic, 2% v/v B-27 supplement, 1mM L-glutamine) with 0.18% w/v D-[+]-glucose. We produced single species neuron-astrocyte co-cultures were produced by plating 0.5mL rat suspension onto poly-D-lysine and laminin (Sigma, 114956-81-9) coated 24 well tissue culture plates. For these, after 2 hours and 30 minutes, OptiMEM I+ was aspirated and replaced with 1% NBA without AraC. On DIV0 1mL 1% NBA with AraC was added.

#### Extraction of primary mouse microglia from mixed glial cultures

After 12-14 days of growth, we placed mixed glial cultures on an orbital shaker at 150rpm for one hour at 37°C to detach microglia. We collected the media and centrifuged at 150xG for five minutes. We resuspended pellets in 0% NBA to achieve a concentration of 2x10^5^ live cells per mL. 1mL of this was plated on top of DIV7 neuron-astrocyte cultures, having first removing the media from each well. These were incubated for two days to produce co-cultures of WT or KO primary mouse microglia, WT human astrocytes and WT rat neurons for experimentation.

#### Mouse blood clot conditioned media

We extracted blood from >12-week-old C57BL/6J WT mice by decapitation following overdose with a lethal dose of sodium pentobarbital (Pentoject). We mixed 4.5mL whole blood without anticoagulants with 90mL transfection medium with insulin (TMI; 89% salt-glucose-glycine solution [sodium chloride, sodium bicarbonate, potassium chloride, magnesium chloride, calcium chloride, HEPES, glycine, glucose, sodium pyruvate], 10% minimum essential medium [Gibco, 51200038], 1x antibiotic-antimycotic, 7.5μg/mL bovine insulin [Sigma, I0516]). We disaggregated blood clot in this solution with a 10ml pipette and placed half of the total volume in an immersion sonicator (Langford Ultrasonics, 575H) for 15 minutes before recombining the sonicated and non-sonicated components and plating the mixture onto T175 flasks. These were incubated for 48h to allow leukocytes and lymphocytes of the nonsonicated blood to respond to cell lysis as well as for processes of thrombosis and fibrinolysis, and evaporation of pentobarbital to occur. We then aspirated the blood clot conditioned media (CCM) and sonicated for 15 minutes before centrifuging at 2000xG for 15 minutes. The pellet was discarded, and the supernatant passed through a 0.22µM filter. We used phase-contrast microscopy was to ensure no debris or erythrocytes were present in the filtered CCM and the limulus amebocyte lysate test (Pierce, 88282) to ensure that there was no detectible endotoxin. Neat CCM was frozen and stored at −80°C for up to six months. On day of use, it was warmed to 37°C and diluted to 25% using warmed TMI and was not refrozen. All experiments used the same batch of CCM.

#### Stimulation of cultures

On the day of simulation, we exchanged the culture medium of our mixed species cultures with TMI. Two hours later, CDDO-TFEA was added to wells as appropriate to achieve a final concentration of 100nM. 24h later media was aspirated and replaced with 1ml CCM or TMI. Wells which had been pre-treated with CDDO-TFEA received a further 100nM dose. We added LPS (150ng/mL) or hydrogen peroxide (200µM or 400µM) to appropriate wells. We then returned all cells to the incubator for 24h before terminating the experiment.

#### Cell death

We quantified cell death using nuclear pyknosis. To measure this, we fixed cells using a solution of 3% paraformaldehyde (PFA; Sigma, P6148) and 4% sucrose in phosphate buffered saline (PBS) for 20 minutes, washed and permeabilised cells using 0.5% NP-40 (Sigma, 492016) in PBS. We then coverslipped using mounting medium with 4′,6-diamidino-2-phenylindole (DAPI; Vector, H-1200-10). We imaged three fields of view per well using a Leica AF6000 microscope with a DFC30X digital camera and counted the percentage of pyknotic nuclei manually, masked to culture setting and after online randomisation of the order of images (https://www.random.org/lists/).

#### Immunocytochemistry

We used fluorescence immunocytochemistry to visualise each cell type in mixed species culture. Cultures were fixed and permeabilised as above and a cocktail of primary antibodies applied (microglia [Rabbit anti-IBA1 1:500; Wako, 019-19741], astrocytes [rat anti-GFAP 1:500; Invitrogen 13-0300] and neurons [mouse anti-pan-neuronal Alexa Fluor-488, directly conjugated Neuro-Chrom 1:500; Milli-Mark MAB2300X]) for 12h, protected from light on an orbital shaker at 15rpm. We removed primary antibodies and applied secondaries (anti-rabbit IgG [Alexa Fluor-594, 1:200; Abcam, ab150080] and anti-rat IgG [Alexa Fluor 350, 1:200; Invitrogen, A21093]) for 1h at room temperature on an orbital shaker. These were washed off and we applied glass coverslips using fluorescence mounting medium (Vector, H-1000-10) before imaging as above.

#### RNA isolation and sequencing

We isolated RNA immediately on termination of each experiment using High Pure RNA Tissue kits (Roche 12033674001) according to manufacturer’s instructions. We prepared TruSeq stranded mRNA-seq libraries (Illumina) and sequenced these in an S2 flow cell using a NovaSeq 6000 sequencer (Illumina) to a depth of approximately 100 million 100bp paired-end reads per sample. We used SARGASSO, (Sargasso Assigns Reads to Genomes According to Species-Specific Origin) as described previously, to assign reads to cell type according to species, and performed read alignment and differential expression analysis as described earlier^13^.

### Statistical analysis

For RNA sequencing data analysis we used RStudio running R Core^86,87^. We used DEseq2 to determine log_2_ fold changes (L2FC) in individual gene expression and Benjamini-Hochberg adjusted p-values (p_adj_) to determine differential gene expression^71^. For *in vitro* comparisons, pairwise analyses were used with matching of samples arising from the same animals and on the same cell culture plate. To identify enrichment of the Molecular Signatures Database Hallmark gene sets^88,89^ and the ChIP-X ENCODE-ChEA consensus Enrichr gene sets^26^ in a hypothesis-free fashion, we used two sequential enrichment tests, ordered by L2FC (fgsea v1.10.1; gage v2.34.0) and called enriched sets if they were significantly enriched (p_adj_<0.05) in both^90,91^. We present the normalised enrichment scores (NES) and p_adj_ derived from the fgsea test. As the ChIP-X ENCODE-ChEA gene sets vary widely in set size, which can bias normalised enrichment score towards greater enrichment of smaller sets, we present the normalised enrichment scores from the larger and smaller significantly enriched gene sets separately. For hypothesis-driven enrichment tests we applied the fgsea test alone. For gene coexpression network (GCN) analyses we used Graphia (v2.1.)^73^. For GCN analyses we excluded all genes with an FPKM value of 0 in any sample so as to prevent spurious correlation of undetected transcripts. For all other RNA sequencing analyses, we only included genes with a mean FPKM greater than 1. For comparisons of NRF2^ΔMMC^ versus NRF2^LoxP^ mouse RNA sequencing data, we excluded genes which were differentially expressed (padj<0.05 or >1.5 DESeq2 fold change) with expression of the Cx3cr1-cre construct in sham operated mice. For sorted astrocytes we detected transcriptomic evidence of minor contamination of our sorted fraction with MMCs. To address this we excluded genes with a minimum FPKM <5 in any sample or that was defined as a myeloid-specific gene in our GCN analyses.^28^ We excluded genes with predicted misassignment to cell type >10% in any sample from analyses of mixed species cell culture.

For probe-based spatial transcriptomic analysis, fields of view failing AtoMx quality control (qcFlagsFOV) and cells failing cell-level AtomMx quality control (qcCellsPassed) were excluded. Gene expression counts were log-normalised using Seurat (v4.3.0) and called for downstream analyses.

Cells were assigned a distance to the nearest haematoma margin, with values of 0 indicating cells within the haematoma and distance bins defined in 1mm increments extending outward from the margin. Cell types were annotated by mapping cells to the Allen Brain Atlas Human MTG SEA-AD taxonomy using the deep generative mapping algorithm implemented in MapMyCells (v1.6.1).^30^

Differential expression was performed on pseudobulk-aggregated counts using DESeq2 (v1.36.0), modelling the effects of time from ICH onset (acute, chronic, control) and distance from the haematoma.^71^

For mass spectrometry imaging we collected data from regions of haematoma and then three 500um thick circumferential regions moving outwards from the haematoma margin. Data files were analysed with the SCiLs lab software package (v2025b Core, Bruker) and mean abundance levels for rounded m/z ratios for each slide region were extracted for processing within R (version 4.1.3). Abundance levels were then re-calibrated using spline-based correction using abundant matrix markers within the data. M/z ratios were mapped to protein abundance. Signal from multiple peptides attributed to the same protein were summed. The 25% most lowly expressed proteins across all samples were dropped from analysis. No imputation of missing data was used. To reduce skew, we normalised data using a log_2_(intensity+1) transform and then analysed using a mixed effects type III ANOVA with fixed effects of genotype, or time from ICH onset (depending on experiment), distance from haematoma, and the interaction between these effects. Random effects of slide and subject were included. Proteins with the 20% highest residual variation were dropped and fixed effects were adjusted for multiple testing using the Benjamini-Hochberg method. As no statistically significant effects were found following this, unadjusted analyses are presented but with strong cautioning required for interpretation and a requirement for future external validation/refutation of findings.

We throughout this paper we used two-tailed t-tests, one-way and two-way ANOVA for continuous normally distributed data with Welsh and Geisser-Greenhouse corrections applied as appropriate^86,87,92^. For *post-hoc* multiple comparisons after one-way and two-way ANOVA we elected to use Dunnett’s T3 and the Holm-Šidák test, respectively, *a priori*. For non-normally distributed data we used Mann-Whitney U and Kruskal Wallis tests. For all analyses we considered a p-value or p_adj_-value <0.05 to indicate statistical significance.

### Data Availability

The following RNAseq data have been deposited at the EMBL EBI:

ArrayExpress (https://www.ebi.ac.uk/biostudies/arrayexpress). E-MTAB-14323: “RNA-seq of perihaematomal and anatomically matched contralateral brain tissue after intracerebral hemorrhage”; E-MTAB-14324: “RNA-seq of microglia and monocyte-derived cells in NRF2 knockout and wild-type mice after induced intracerebral hemorrhage or sham treatment”; E-MTAB-14327: “RNA-seq of triple cultures of neurons, astrocytes and microglia treated with blood clot conditioned media, lipopolysaccharide, and NRF2-activating drug CDDO-TFEA.”

## ACKNOWLEDGEMENTS

Tissue samples were supplied by The Edinburgh Brain and Tissue bank, which is supported by the Medical Research Council, and the Manchester Brain Bank, which is part of the Brains for Dementia Research programme, jointly funded by Alzheimer’s Research UK and Alzheimer’s Society. We gratefully acknowledge the technical assistance of Ying Zhou in preparing and maintaining cell cultures. RNA sequencing was performed by Edinburgh Genomics, which is funded by the UK NERC, BBSRC and MRC and by Cambridge Genomic Services. Cell sorting was performed by Dr Martin Waterfall of the Ashworth Laboratories flow cytometry facility at the University of Edinburgh. We acknowledge the assistance of Dr Pete Bankhead in writing scripts for use in QuPath. Slide scanning was performed at the Histology Department of the Queen’s Medical Research Institute at the University of Edinburgh and confocal microscopy at the IMPACT facility in Hugh Robson Building at the Centre for Discovery Brain Sciences of the University of Edinburgh. Tissue sectioning and mounting for CosMx analysis was undertaken by Bev Notman, research assistant at the Centre for Clinical Brain Sciences, University of Edinburgh.

## AUTHOR CONTRIBUTIONS

Conceptualisation: JJML, JQ, CS, NS, RASS, BWM, GEH. Data curation: JJML, XH, ORD. Formal analysis: JJML, XH, FS, YC, ORD. Funding Acquisition: JJML, NS, RASS, BWM, GEH. Investigation: JJML, JQ, PB, SM, CL, KM, FS, YC, BG, NS. Methodology: JJML, JQ, JB, PB, SM, CK, KM, JM, BG, ORD, CS, NS, RASS, BWM, GEH. Resources: CS, RASS, BWM, GEH. Software: JJML, YC, XH, BG, ORD. Supervision: CS, NS, RASS, BWM, GEH. Writing – original draft: JJML. Writing – review and editing: All.

## SUPPLEMENTARY DATA

Supplementary tables and figures

