## Supplementary for "Mononuclear myeloid cells mount an adaptive protective response to intracerebral haemorrhage by activation of NRF2"

**Index**

**Supplementary Table 1: Characteristics of included patients in RNA sequencing studies**
Related to figures 1,2, and 6

| Age | Sex | Death-autopsy interval (hours) | RIN Perihaematomal | RIN Contralateral | Pathological findings | Previous TIA | Previous Ischaemic stroke | Cause of death | ICH location | ICH volume (ml, abc/2) | ICH score | ICH Onset to death (days) |
|---|---|---|---|---|---|---|---|---|---|---|---|---|
| **Cohort 1: ICH cases** | | | | | | | | | | | | |
| 85 | Female | 85 | 6.3 | NA | ICH, Severe SVD, BB II | No | No | ICH, pneumonia | Lobar | 89 | 4 | 7 |
| 83 | Female | 35 | 6.2 | NA | ICH, BB III, CAA | No | No | ICH | Lobar | 8 | 1 | 7 |
| 70 | Male | 82 | 5.1 | NA | ICH, BB I, Severe SVD | No | Yes | Aspiration pneumonia, ICH | Lobar | 67 | 2 | 8 |
| 57 | Male | 83 | 5.7 | NA | ICH, Severe SVD | No | No | ICH, hepatic failure | Lobar | 17 | 1 | 27 |
| **Cohort 1: sudden non-neurological death controls** | | | | | | | | | | | | |
| 87 | Female | 24 | 6.1 | NA | No abnormality | No | No | CCF | NA | NA | NA | NA |
| 76 | Female | 129.5 | 7.3 | NA | No abnormality | No | No | IHD | NA | NA | NA | NA |
| 71 | Female | 96 | 6.6 | NA | No abnormality | No | No | IHD | NA | NA | NA | NA |
| 57 | Male | 64 | 7.3 | NA | Mild SVD | No | No | IHD | NA | NA | NA | NA |
| 71 | Female | 95 | 5.8 | NA | Mild SVD | No | No | Suffocation | NA | NA | NA | NA |
| 72 | Male | 60 | 6.3 | NA | Mild SVD | No | No | IHD | NA | NA | NA | NA |
| 73 | Male | 66 | 5.9 | NA | Moderate SVD, mild CAA, BB I | No | No | IHD | NA | NA | NA | NA |
| **Cohort 2: ICH cases with bilateral sampling** | | | | | | | | | | | | |
| 59 | Male | 39 | 6.2 | NA | ICH, Severe SVD, CAA | No | Yes | ICH | Basal ganglia | 22 | 3 | 1 |
| 74 | Female | 31 | NA | 6 | ICH, CAA, BB VI, severe SVD | Yes | Yes | ICH, HTN, vascular dementia | Probable lobar | 146 | 3 | 1 |
| 91 | Male | 132 | 6.0 | 5 | ICH, CAA, BB II, severe SVD | Yes | No | ICH, AF, vascular dementia, TIAs | Lobar | 47 | 4 | 1 |
| 78 | Male | 27 | 4.2 | 6 | ICH, severe non amyloid SVD | No | No | ICH, HTN | Probable deep | 107 | 3 | 1 |
| 67 | Male | 29 | 6.9 | NA | ICH CAA, BB I, severe SVD | No | Yes | ICH, ALD | Basal Ganglia | 50 | 1 | 8 |
| 68 | Male | 30.5 | 7.1 | 5.6 | ICH, BB II, brain stem Lewy Body Disease, moderate SVD, primary age related tauopathy | No | No | ICH | Lobar | 88 | 2 | 11 |
| 77 | Male | 23 | 5.9 | 6.2 | ICH BB II, severe SVD, primary age related tauopathy | No | No | ICH | Probable deep | 76 | 3 | 4 |
| 85 | Female | 24 | 6.2 | 7.3 | ICH, CAA, BB V, moderate SVD, | No | No | ICH, aspiration pneumonia, AD | Probable lobar | 116 | 3 | 8 |

RIN: RNA integrity number; TIA: Transient ischaemic attack; ICH: intracerebral haematoma; SVD: non-amyloid small vessel disease; BB: Braak and Braak stage; CAA: cerebral amyloid angiopathy; NA: Not applicable/sample not obtained/not sequenced; CCF: congestive cardiac failure; IHD: ischaemic heart disease; HTN: essential hypertension; AF: atrial fibrillation, ALD: alcohol associated liver disease (cirrhosis); AD: Alzheimer’s disease.

### Supplementary Table 2: Cluster 16 genes

Related to figure 1. Data and scripts to derive all gene clusters are available in supplementary data.

| Gene name | Ensembl ID | Entrez ID | Description |
|---|---|---|---|
| CLU | ENSG00000120885 | 1191 | clusterin [Source:HGNC Symbol;Acc:HGNC:2095] |
| ITM2C | ENSG00000135916 | 81618 | integral membrane protein 2C [Source:HGNC Symbol;Acc:HGNC:6175] |
| AQP4 | ENSG00000171885 | 361 | aquaporin 4 [Source:HGNC Symbol;Acc:HGNC:637] |
| CD63 | ENSG00000135404 | 967 | CD63 molecule [Source:HGNC Symbol;Acc:HGNC:1692] |
| CNN3 | ENSG00000117519 | 1266 | calponin 3 [Source:HGNC Symbol;Acc:HGNC:2157] |
| DTNA | ENSG00000134769 | 1837 | dystrobrevin alpha [Source:HGNC Symbol;Acc:HGNC:3057] |
| OST4 | ENSG00000228474 | 100128731 | oligosaccharyltransferase complex subunit 4, non-catalytic [Source:HGNC Symbol;Acc:HGNC:32483] |
| TXNIP | ENSG00000265972 | 10628 | thioredoxin interacting protein [Source:HGNC Symbol;Acc:HGNC:16952] |
| DAD1 | ENSG00000129562 | 1603 | defender against cell death 1 [Source:HGNC Symbol;Acc:HGNC:2664] |
| HSPB8 | ENSG00000152137 | 26353 | heat shock protein family B (small) member 8 [Source:HGNC Symbol;Acc:HGNC:30171] |
| ANXA5 | ENSG00000164111 | 308 | annexin A5 [Source:HGNC Symbol;Acc:HGNC:543] |
| STOM | ENSG00000148175 | 2040 | stomatin [Source:HGNC Symbol;Acc:HGNC:3383] |
| LAPTM4A | ENSG00000068697 | 9741 | lysosomal protein transmembrane 4 alpha [Source:HGNC Symbol;Acc:HGNC:6924] |
| TMBIM1 | ENSG00000135926 | 64114 | transmembrane BAX inhibitor motif containing 1 [Source:HGNC Symbol;Acc:HGNC:23410] |
| ZFP36L1 | ENSG00000185650 | 677 | ZFP36 ring finger protein like 1 [Source:HGNC Symbol;Acc:HGNC:1107] |
| MGST1 | ENSG00000008394 | 4257 | microsomal glutathione S-transferase 1 [Source:HGNC Symbol;Acc:HGNC:7061] |
| NPC2 | ENSG00000119655 | 10577 | NPC intracellular cholesterol transporter 2 [Source:HGNC Symbol;Acc:HGNC:14537] |
| MT1X | ENSG00000187193 | 4501 | metallothionein 1X [Source:HGNC Symbol;Acc:HGNC:7405] |
| MCL1 | ENSG00000143384 | 4170 | MCL1, BCL2 family apoptosis regulator [Source:HGNC Symbol;Acc:HGNC:6943] |
| CD59 | ENSG00000085063 | 966 | CD59 molecule (CD59 blood group) [Source:HGNC Symbol;Acc:HGNC:1689] |
| ID4 | ENSG00000172201 | 3400 | inhibitor of DNA binding 4, HLH protein [Source:HGNC Symbol;Acc:HGNC:5363] |
| UBE2L6 | ENSG00000156587 | 9246 | ubiquitin conjugating enzyme E2 L6 [Source:HGNC Symbol;Acc:HGNC:12490] |
| ALDH1L1 | ENSG00000144908 | 10840 | aldehyde dehydrogenase 1 family member L1 [Source:HGNC Symbol;Acc:HGNC:3978] |
| PLPP1 | ENSG00000067113 | 8611 | phospholipid phosphatase 1 [Source:HGNC Symbol;Acc:HGNC:9228] |
| MSN | ENSG00000147065 | 4478 | moesin [Source:HGNC Symbol;Acc:HGNC:7373] |
| CCNG1 | ENSG00000113328 | 900 | cyclin G1 [Source:HGNC Symbol;Acc:HGNC:1592] |
| SCRG1 | ENSG00000164106 | 11341 | stimulator of chondrogenesis 1 [Source:HGNC Symbol;Acc:HGNC:17036] |
| CDKN1A | ENSG00000124762 | 1026 | cyclin dependent kinase inhibitor 1A [Source:HGNC Symbol;Acc:HGNC:1784] |
| SMIM30 | ENSG00000214194 | 401397 | small integral membrane protein 30 [Source:HGNC Symbol;Acc:HGNC:48953] |
| RAB13 | ENSG00000143545 | 5872 | RAB13, member RAS oncogene family [Source:HGNC Symbol;Acc:HGNC:9762] |
| TMEM176B | ENSG00000106565 | 28959 | transmembrane protein 176B [Source:HGNC Symbol;Acc:HGNC:29596] |
| CEBPD | ENSG00000221869 | 1052 | CCAAT/enhancer binding protein delta [Source:HGNC Symbol;Acc:HGNC:1835] |
| CHPT1 | ENSG00000111666 | 56994 | choline phosphotransferase 1 [Source:HGNC Symbol;Acc:HGNC:17852] |
| MSI2 | ENSG00000153944 | 124540 | musashi RNA binding protein 2 [Source:HGNC Symbol;Acc:HGNC:18585] |
| PALLD | ENSG00000129116 | 23022 | palladin, cytoskeletal associated protein [Source:HGNC Symbol;Acc:HGNC:17068] |
| SYF2 | ENSG00000117614 | 25949 | SYF2 pre-mRNA splicing factor [Source:HGNC Symbol;Acc:HGNC:19824] |
| HHLA3 | ENSG00000197568 | 11147 | HERV-H LTR-associating 3 [Source:HGNC Symbol;Acc:HGNC:4906] |
| ARRDC4 | ENSG00000140450 | 91947 | arrestin domain containing 4 [Source:HGNC Symbol;Acc:HGNC:28087] |
| FCGRT | ENSG00000104870 | 2217 | Fc fragment of IgG receptor and transporter [Source:HGNC Symbol;Acc:HGNC:3621] |
| BCCIP | ENSG00000107949 | 56647 | BRCA2 and CDKN1A interacting protein [Source:HGNC Symbol;Acc:HGNC:978] |
| TNFRSF1A | ENSG00000067182 | 7132 | TNF receptor superfamily member 1A [Source:HGNC Symbol;Acc:HGNC:11916] |
| TMEM176A | ENSG00000002933 | 55365 | transmembrane protein 176A [Source:HGNC Symbol;Acc:HGNC:24930] |
| PIR | ENSG00000087842 | 8544 | pirin [Source:HGNC Symbol;Acc:HGNC:30048] |
| ARRDC3 | ENSG00000113369 | 57561 | arrestin domain containing 3 [Source:HGNC Symbol;Acc:HGNC:29263] |
| P4HA1 | ENSG00000122884 | 5033 | prolyl 4-hydroxylase subunit alpha 1 [Source:HGNC Symbol;Acc:HGNC:8546] |
| LRRC3B | ENSG00000179796 | 116135 | leucine rich repeat containing 3B [Source:HGNC Symbol;Acc:HGNC:28105] |
| CBFB | ENSG00000067955 | 865 | core-binding factor beta subunit [Source:HGNC Symbol;Acc:HGNC:1539] |
| ARHGEF6 | ENSG00000129675 | 9459 | Rac/Cdc42 guanine nucleotide exchange factor 6 [Source:HGNC Symbol;Acc:HGNC:685] |
| SAMHD1 | ENSG00000101347 | 25939 | SAM and HD domain containing deoxynucleoside triphosphate triphosphohydrolase 1 [Source:HGNC Symbol;Acc:HGNC:15925] |
| PDCD4 | ENSG00000150593 | 27250 | programmed cell death 4 [Source:HGNC Symbol;Acc:HGNC:8763] |
| BTG1 | ENSG00000133639 | 694 | BTG anti-proliferation factor 1 [Source:HGNC Symbol;Acc:HGNC:1130] |
| AKR1C3 | ENSG00000196139 | 8644 | aldo-keto reductase family 1 member C3 [Source:HGNC Symbol;Acc:HGNC:386] |
| SEC11A | ENSG00000140612 | 23478 | SEC11 homolog A, signal peptidase complex subunit [Source:HGNC Symbol;Acc:HGNC:17718] |
| SHC1 | ENSG00000160691 | 6464 | SHC adaptor protein 1 [Source:HGNC Symbol;Acc:HGNC:10840] |
| DRAM2 | ENSG00000156171 | 128338 | DNA damage regulated autophagy modulator 2 [Source:HGNC Symbol;Acc:HGNC:28769] |
| DERA | ENSG00000023697 | 51071 | deoxyribose-phosphate aldolase [Source:HGNC Symbol;Acc:HGNC:24269] |
| DYNLT1 | ENSG00000146425 | 6993 | dynein light chain Tctex-type 1 [Source:HGNC Symbol;Acc:HGNC:11697] |
| HVCN1 | ENSG00000122986 | 84329 | hydrogen voltage gated channel 1 [Source:HGNC Symbol;Acc:HGNC:28240] |
| MAGT1 | ENSG00000102158 | 84061 | magnesium transporter 1 [Source:HGNC Symbol;Acc:HGNC:28880] |
| PLCD1 | ENSG00000187091 | 5333 | phospholipase C delta 1 [Source:HGNC Symbol;Acc:HGNC:9060] |
| HSCB | ENSG00000100209 | 150274 | HscB mitochondrial iron-sulfur cluster cochaperone [Source:HGNC Symbol;Acc:HGNC:28913] |
| ETFDH | ENSG00000171503 | 2110 | electron transfer flavoprotein dehydrogenase [Source:HGNC Symbol;Acc:HGNC:3483] |
| FAM189A2 | ENSG00000135063 | 9413 | family with sequence similarity 189 member A2 [Source:HGNC Symbol;Acc:HGNC:24820] |

| Gene name | Ensembl ID | Entrez ID | Description |
|---|---|---|---|
| FSTL1 | ENSG00000163430 | 11167 | follistatin like 1 [Source:HGNC Symbol;Acc:HGNC:3972] |
| MOB1A | ENSG00000114978 | 55233 | MOB kinase activator 1A [Source:HGNC Symbol;Acc:HGNC:16015] |
| FNTA | ENSG00000168522 | 2339 | farnesyltransferase, CAAX box, alpha [Source:HGNC Symbol;Acc:HGNC:3782] |
| TRIM22 | ENSG00000132274 | 10346 | tripartite motif containing 22 [Source:HGNC Symbol;Acc:HGNC:16379] |
| IL17RC | ENSG00000163702 | 84818 | interleukin 17 receptor C [Source:HGNC Symbol;Acc:HGNC:18358] |
| PYGL | ENSG00000100504 | 5836 | glycogen phosphorylase L [Source:HGNC Symbol;Acc:HGNC:9725] |
| CWF19L2 | ENSG00000152404 | 143884 | CWF19 like 2, cell cycle control (S. pombe) [Source:HGNC Symbol;Acc:HGNC:26508] |
| CD302 | ENSG00000241399 | 9936 | CD302 molecule [Source:HGNC Symbol;Acc:HGNC:30843] |
| CMTM6 | ENSG00000091317 | 54918 | CKLF like MARVEL transmembrane domain containing 6 [Source:HGNC Symbol;Acc:HGNC:19177] |
| ATL3 | ENSG00000184743 | 25923 | atlastin GTPase 3 [Source:HGNC Symbol;Acc:HGNC:24526] |
| CD4 | ENSG00000010610 | 920 | CD4 molecule [Source:HGNC Symbol;Acc:HGNC:1678] |
| SLC7A2 | ENSG00000003989 | 6542 | solute carrier family 7 member 2 [Source:HGNC Symbol;Acc:HGNC:11060] |
| LIMS1 | ENSG00000169756 | 3987 | LIM zinc finger domain containing 1 [Source:HGNC Symbol;Acc:HGNC:6616] |
| ANO6 | ENSG00000177119 | 196527 | anoctamin 6 [Source:HGNC Symbol;Acc:HGNC:25240] |
| PIGF | ENSG00000151665 | 5281 | phosphatidylinositol glycan anchor biosynthesis class F [Source:HGNC Symbol;Acc:HGNC:8962] |
| TGIF1 | ENSG00000177426 | 7050 | TGFB induced factor homeobox 1 [Source:HGNC Symbol;Acc:HGNC:11776] |
| ITPKC | ENSG00000086544 | 80271 | inositol-trisphosphate 3-kinase C [Source:HGNC Symbol;Acc:HGNC:14897] |
| ABHD5 | ENSG00000011198 | 51099 | abhydrolase domain containing 5 [Source:HGNC Symbol;Acc:HGNC:21396] |
| SLC7A6OS | ENSG00000103061 | 84138 | solute carrier family 7 member 6 opposite strand [Source:HGNC Symbol;Acc:HGNC:25807] |
| STK17A | ENSG00000164543 | 9263 | serine/threonine kinase 17a [Source:HGNC Symbol;Acc:HGNC:11395] |
| PQLC3 | ENSG00000162976 | 130814 | PQ loop repeat containing 3 [Source:HGNC Symbol;Acc:HGNC:28503] |
| AFF1 | ENSG00000172493 | 4299 | AF4/FMR2 family member 1 [Source:HGNC Symbol;Acc:HGNC:7135] |
| USP53 | ENSG00000145390 | 54532 | ubiquitin specific peptidase 53 [Source:HGNC Symbol;Acc:HGNC:29255] |
| PARP4 | ENSG00000102699 | 143 | poly(ADP-ribose) polymerase family member 4 [Source:HGNC Symbol;Acc:HGNC:271] |
| PIEZO1 | ENSG00000103335 | 9780 | piezo type mechanosensitive ion channel component 1 [Source:HGNC Symbol;Acc:HGNC:28993] |
| SMAD9 | ENSG00000120693 | 4093 | SMAD family member 9 [Source:HGNC Symbol;Acc:HGNC:6774] |
| LINC01094 | ENSG00000251442 | NA | long intergenic non-protein coding RNA 1094 [Source:HGNC Symbol;Acc:HGNC:49219] |
| TGFBR1 | ENSG00000106799 | 7046 | transforming growth factor beta receptor 1 [Source:HGNC Symbol;Acc:HGNC:11772] |
| FAS | ENSG00000026103 | 355 | Fas cell surface death receptor [Source:HGNC Symbol;Acc:HGNC:11920] |
| ADA2 | ENSG00000093072 | 51816 | adenosine deaminase 2 [Source:HGNC Symbol;Acc:HGNC:1839] |
| FCF1 | ENSG00000119616 | 51077 | FCF1, rRNA-processing protein [Source:HGNC Symbol;Acc:HGNC:20220] |
| TMOD3 | ENSG00000138594 | 29766 | tropomodulin 3 [Source:HGNC Symbol;Acc:HGNC:11873] |
| MAP3K20 | ENSG00000091436 | 51776 | mitogen-activated protein kinase kinase kinase 20 [Source:HGNC Symbol;Acc:HGNC:17797] |
| TGDS | ENSG00000088451 | 23483 | TDP-glucose 4,6-dehydratase [Source:HGNC Symbol;Acc:HGNC:20324] |
| TMBIM4 | ENSG00000155957 | 51643 | transmembrane BAX inhibitor motif containing 4 [Source:HGNC Symbol;Acc:HGNC:24257] |
| SP110 | ENSG00000135899 | 3431 | SP110 nuclear body protein [Source:HGNC Symbol;Acc:HGNC:5401] |
| TRAF3IP2 | ENSG00000056972 | 10758 | TRAF3 interacting protein 2 [Source:HGNC Symbol;Acc:HGNC:1343] |
| BHMT2 | ENSG00000132840 | 23743 | betaine--homocysteine S-methyltransferase 2 [Source:HGNC Symbol;Acc:HGNC:1048] |
| PARP14 | ENSG00000173193 | 54625 | poly(ADP-ribose) polymerase family member 14 [Source:HGNC Symbol;Acc:HGNC:29232] |
| SQOR | ENSG00000137767 | 58472 | sulfide quinone oxidoreductase [Source:HGNC Symbol;Acc:HGNC:20390] |
| SLC25A51 | ENSG00000122696 | 92014 | solute carrier family 25 member 51 [Source:HGNC Symbol;Acc:HGNC:23323] |
| SYNC | ENSG00000162520 | NA | syncoilin, intermediate filament protein [Source:HGNC Symbol;Acc:HGNC:28897] |
| TNFRSF11 B | ENSG00000164761 | 4982 | TNF receptor superfamily member 11b [Source:HGNC Symbol;Acc:HGNC:11909] |
| KDELC2 | ENSG00000178202 | 143888 | KDEL motif containing 2 [Source:HGNC Symbol;Acc:HGNC:28496] |
| AC083799. 1 | ENSG00000203644 | NA | NA |
| HEBP2 | ENSG00000051620 | 23593 | heme binding protein 2 [Source:HGNC Symbol;Acc:HGNC:15716] |
| IRAK4 | ENSG00000198001 | 51135 | interleukin 1 receptor associated kinase 4 [Source:HGNC Symbol;Acc:HGNC:17967] |
| RGS9 | ENSG00000108370 | 8787 | regulator of G protein signaling 9 [Source:HGNC Symbol;Acc:HGNC:10004] |
| LMOD1 | ENSG00000163431 | 25802 | leiomodin 1 [Source:HGNC Symbol;Acc:HGNC:6647] |
| MYD88 | ENSG00000172936 | 4615 | myeloid differentiation primary response 88 [Source:HGNC Symbol;Acc:HGNC:7562] |
| LYN | ENSG00000254087 | 4067 | LYN proto-oncogene, Src family tyrosine kinase [Source:HGNC Symbol;Acc:HGNC:6735] |
| SLC25A43 | ENSG00000077713 | 203427 | solute carrier family 25 member 43 [Source:HGNC Symbol;Acc:HGNC:30557] |
| ZC3HAV1 | ENSG00000105939 | 56829 | zinc finger CCCH-type containing, antiviral 1 [Source:HGNC Symbol;Acc:HGNC:23721] |
| NID1 | ENSG00000116962 | 4811 | nidogen 1 [Source:HGNC Symbol;Acc:HGNC:7821] |
| NEDD1 | ENSG00000139350 | 121441 | neural precursor cell expressed, developmentally down-regulated 1 [Source:HGNC Symbol;Acc:HGNC:7723] |
| EHD4 | ENSG00000103966 | 30844 | EH domain containing 4 [Source:HGNC Symbol;Acc:HGNC:3245] |
| SLC35D1 | ENSG00000116704 | 23169 | solute carrier family 35 member D1 [Source:HGNC Symbol;Acc:HGNC:20800] |
| LPP | ENSG00000145012 | 4026 | LIM domain containing preferred translocation partner in lipoma [Source:HGNC Symbol;Acc:HGNC:6679] |
| AL392172.1 | ENSG00000228106 | NA | NA |
| OTULIN | ENSG00000154124 | 90268 | OTU deubiquitinase with linear linkage specificity [Source:HGNC Symbol;Acc:HGNC:25118] |

### Supplementary Table 3: Characteristics of patients included in cohort 3

Related to figure 2.

| | Age | Sex | ICH onset to death (d) | Death-autopsy interval (h) | Non-ICH pathological findings | Primary causes of death | ICH location | ICH volume (ml) |
|---|---|---|---|---|---|---|---|---|
| Case | 80 | Male | 3 | 83 | Lacunar infarcts | ICH | Thalamus | 18 |
| Case | 85 | Female | 3 | 36 | Lacunar infarcts, Braak tangle stage III | ICH | Frontal | 121 |
| Case | 78 | Female | 3 | 40 | Lacunar infarcts | ICH | Frontal | 98 |
| Case | 67 | Male | 3 | 103 | Sporadic CAA, Braak tangle stage IV | ICH | Frontal | 76 |
| Case | 88 | Female | 3 | 65 | Moderate non-amyloid SVD, severe arteriolar Aß-CAA, Braak tangle stage III | ICH | Temporal | 90 |
| Case | 63 | Male | 3 | 28 | Glioblastoma | Aspiration pneumonia secondary to ICH | Basal ganglia | 51 |
| Case | 83 | Male | 4 | 88 | Sporadic CAA, Braak tangle stage V | ICH | Frontal | 94 |
| Case | 79 | Male | 4 | 84 | Sporadic CAA Braak tangle stage I | ICH | Thalamus | 24 |
| Case | 81 | Female | 5 | 46 | Sporadic CAA | ICH | Temporal | 62 |
| Case | 85 | Female | 5 | 27 | Lacunar infarcts, Braak tangle stage II | ICH | Thalamus | 7 |
| Case | 77 | Female | 5 | 79 | Lacunar infarcts, brain stem Lewy body disease | ICH | Thalamus | 6 |
| Case | 64 | Female | 6 | 57 | Lacunar infarcts | Aspiration pneumonia secondary to ICH | Thalamus | 8 |
| Case | 81 | Male | 6 | 77 | Moderate non-amyloid SVD, mild arteriolar Aß-CAA | Aspiration pneumonia secondary to ICH | Temporal | 64 |
| Case | 75 | Male | 7 | 35 | Infarct/s >10 mm in diameter | Bronchopneumonia secondary to ICH | Thalamus | 29 |
| Case | 85 | Female | 7 | 85 | Lacunar infarcts, Braak tangle stage II | ICH | Parietal | 72 |
| Case | 67 | Male | 7 | 77 | Lacunar infarcts, Braak tangle stage II | Aspiration pneumonia secondary to ICH | Thalamus | 12 |
| Case | 80 | Female | 7 | 41 | Sporadic CAA, Braak tangle stage VI | Bronchopneumonia secondary to ICH | Parietal | 115 |
| Case | 87 | Male | 8 | 48 | Lacunar infarcts | ICH | Temporal | 46 |
| Case | 85 | Female | 8 | 87 | Lacunar infarcts, sporadic CAA, Braak tangle stage III | ICH | Occipital | 97 |
| Case | 91 | Female | 8 | 41 | Lacunar infarcts | ICH | Basal ganglia | 10 |
| Case | 83 | Male | 8 | 81 | Lacunar infarcts, Braak tangle stage I | Aspiration pneumonia secondary to ICH | Occipital | 21 |
| Case | 77 | Male | 9 | 122 | Lacunar infarcts | Aspiration pneumonia secondary to ICH | Frontal | 61 |
| Case | 75 | Female | 9 | 43 | Moderate non-amyloid SVD, mild arteriolar Aß-CAA, Braak tangle stage II | ICH | Basal ganglia | 8 |
| Case | 75 | Male | 8 | 50 | Infarct/s >10 mm in diameter | Bronchopneumonia secondary to ICH | Thalamus | 8 |
| Control | 78 | Male | NA | 95 | No abnormality detected | Bronchopneumonia and congestive cardiac failure secondary to Hypertensive and ischaemic heart disease | NA | NA |
| Control | 83 | Female | NA | 50 | Non-amyloid SVD | Myocardial infarction secondary to coronary atherosclerosis | NA | NA |
| Control | 79 | Female | NA | 45 | Small vessel disease | Ischaemic heart disease secondary to coronary atherosclerosis | NA | NA |
| Control | 75 | Male | NA | 47 | Small vessel disease | Myocardial infarction secondary to coronary atherosclerosis | NA | NA |
| Control | 79 | Female | NA | 46 | Lacunar infarcts | Haemopericardium secondary to ruptured atherosclerotic aneurysm of the aortic root | NA | NA |
| Control | 76 | Male | NA | 90 | Lacunar infarcts | Myocardial infarction secondary to coronary atherosclerosis | NA | NA |

**Supplementary Table 4: Characteristics of patients included in cohort 4**
Related to figure 3.

| Group | Age | Sex | ICH onset to death (d) | Death-autopsy interval (h) | Non-ICH pathological findings | Primary causes of death | ICH location | ICH volume (ml) |
|---|---|---|---|---|---|---|---|---|
| Acute | 76 | Male | 2 | 111 | Moderate non-amyloid SVD, mild CAA | Intracerebral haemorrhage | Basal ganglia | 24 |
| Acute | 91 | Male | 2 | 83 | Severe non-amyloid SVD, | Intracerebral haemorrhage | Basal ganglia | 144.76 |
| Acute | 88 | Male | 1 | 89 | Moderate non-amyloid SVD, mild CAA | Intracerebral haemorrhage | Parietal | 57.575 |
| Acute | 84 | Male | 1 | 23 | Severe non-amyloid SVD, | Intracerebral haemorrhage | Basal ganglia | 67.704 |
| Chronic | 92 | Female | 21 | 86 | Moderate non-amyloid SVD, | Intracerebral haemorrhage | Basal ganglia | 19.9 |
| Chronic | 78 | Male | 21 | 36 | Severe non-amyloid SVD, moderate CAA | Bronchopneumonia, unspecified | Basal ganglia | 108.3 |
| Chronic | 88 | Female | 41 | 59 | Severe non-amyloid SVD, mild CAA | Intracerebral haemorrhage | Thalamic | 13.662 |
| Chronic | 90 | Female | 13 | 91 | Severe non-amyloid SVD, | Intracerebral haemorrhage | Basal ganglia | 55.7 |
| Control | 82 | Male | NA | 95 | Moderate non-amyloid SVD, severe CAA | Squamous cell carcinoma of the lung | NA | NA |
| Control | 79 | Female | NA | 80 | Mild non-amyloid SVD, Mild CAA | Squamous cell carcinoma of the lung with metastases to liver and peritoneum | NA | NA |
| Control | 81 | Male | NA | 38 | Mild non-amyloid SVD, Mild CAA | Lung cancer non-specified | NA | NA |
| Control | 83 | Female | NA | 78 | Mild non-amyloid SVD, Mild CAA | Acute kidney injury secondary to decompensated cardiac failure | NA | NA |

### Supplementary Table 5: Numbers and exclusions in preclinical studies

Related to figures 4,5,6,7,8, and 9

| Substrate | Analysis and figure numbers | Experimental group | Number of animals used | Exclusions | Reasons for exclusion | Final numbers analysed |
|---|---|---|---|---|---|---|
| Mouse RNA | FACS then RNA sequencing. Figure 4 and 6, supplementary figures 8, 9, 10, 12, 18, and 19. | Sham $Nrf2^{LoxP}$ | n=3 | Nil | NA | n=3 |
| | | ICH $Nrf2^{LoxP}$ | n=6 | 1 | One sample of microglia was found have high expression of Aqp4 and Gfap suggestive of astrocyte contamination and was therefore excluded. | n=5 (microglia), n=6 (MdCs, astrocytes) |
| | | ICH $Nrf2^{\Delta MMC}$ | n=6 | Nil | | n=6 |
| | RT-qPCR. Figure 9, Supplementary figure 23 | ICH WT Vehicle | n=6 | Nil | | n=6 |
| | | ICH WT CDDO-TFEA | n=6 | Nil | | n=6 |
| Mouse brain sections | Haematoma volume analysis. Figure 5, supplementary figure 17 | ICH $Nrf2^{LoxP}$ | n=13 | Nil | | n=13 |
| | | ICH $Nrf2^{\Delta MMC}$ | n=13 | Nil | | n=13 |
| | | ICH $Cx3cr1^{CRE}$ | n=6 | 1 | One mouse reached humane endpoint at day 1. Culled and excluded. | n=5 |
| | | ICH WT | n=6 | Nil | | n=6 |
| | Immunohistochemistry. Figure 5, supplementary figures 13 and 15. | Sham $Nrf2^{LoxP}$ | n=7 | Nil | | n=7 |
| | | Sham $Nrf2^{\Delta MMC}$ | n=7 | Nil | | n=7 |
| | | Sham $Cx3cr1^{CRE}$ | n=7 | Nil | | n=7 |
| | | ICH $Nrf2^{LoxP}$ | n=13 | 1 | 1 mouse brain poor fixation and stain failed blinded quality check. Excluded. | n=12 |
| | | ICH $Nrf2^{\Delta MMC}$ | n=13 | 1 | 1 mouse brain poor fixation and stain failed blinded quality check. Excluded. | n=12 |
| | | ICH $Cx3cr1^{CRE}$ | n=7 | Nil | | n=7 |
| | Immunohistochemistry supplementary figure 17 | ICH $Cx3cr1^{CRE}$ | n=6 | 1 | One mouse reached humane endpoint at day 1. Culled and excluded. | n=5 |
| | | ICH WT | n=6 | Nil | | n=6 |
| | Mass spectrometry imaging Supplementary figures 11 and 16 | Sham WT day 3 | n=2 | Nil | | n=2 |
| | | ICH WT day 1 | n=3 | Nil | | n=3 |
| | | ICH WT day 3 | n=7 | Nil | | n=7 |
| | | ICH WT day 7 | n=4 | Nil | | n=4 |
| | | ICH WT day 28 | n=3 | Nil | | n=3 |
| | | ICH $Nrf2^{LoxP}$ | n=13 | Nil | | n=13 |
| | | ICH $Nrf2^{\Delta MMC}$ | n=13 | Nil | | n=13 |
| Mouse behaviour | Grip strength and NDS LoxP vs. ΔMMC. Figure 5, supplementary figure 14 | Sham $Nrf2^{LoxP}$ | n=7 | Nil | | n=7 |
| | | Sham $Nrf2^{\Delta MMC}$ | n=7 | Nil | | n=7 |
| | | ICH $Nrf2^{LoxP}$ | n=13 | Nil | | 13 |
| | | ICH $Nrf2^{\Delta MMC}$ | n=13 | Nil | | 13 |
| | Weight and sickness (pools IHC/grip strength with FACS sorting analysis) LoxP vs. ΔMMC. Supplementary figure 14 | Sham $Nrf2^{LoxP}$ | n=10 | Nil | | n=10 |
| | | Sham $Nrf2^{\Delta MMC}$ | n=7 | Nil | | n=7 |
| | | ICH $Nrf2^{LoxP}$ | n=19 | Nil | | n=19 |
| | | ICH $Nrf2^{\Delta MMC}$ | n=19 | Nil | | n=19 |
| | Grip strength and NDS Cre vs. WT. Supplementary figure 17 | ICH $Cx3cr1^{CRE}$ | n=6 | 1 | One mouse reached humane endpoint at day 1. Culled and excluded. | n=5 |
| | | ICH WT | n=6 | Nil | | n=6 |
| | Weight and sickness. Cre vs. WT. Supplementary figure 17 | ICH $Cx3cr1^{CRE}$ | n=12 | 1 | One mouse reached humane endpoint at day 1. Culled and excluded. | n=11 |
| | | ICH WT | n=10 | Nil | | n=10 |
| | Weight and sickness. Supplementary figure 23 | ICH WT Vehicle | n=6 | Nil | | n=6 |
| | | ICH WT CDDO-TFEA | n=6 | Nil | | n=6 |

| | | | | | | |
|---|---|---|---|---|---|---|
| | Grip strength NDS, weight and sickness. Figure 9 and supplementary figure 23 | ICH Nrf2$^{\Delta MMC}$ Vehicle | n=14 | 4 | Four sustained complications of gavage. Culled and excluded. | n=10 |
| | | ICH Nrf2$^{\Delta MMC}$ Baricitinib | n=14 | 5 | Five sustained complications of gavage. Culled and excluded. | n=9 |
| Microglia-astrocyte-neuron culture | RNA sequencing with SARGASSO cell-type deconvolution. Figures 7 and 8. Supplementary figures 20 and 21. | Nrf2-KO microglia, WT astrocytes, WT neurons | Microglia n=3<br>Astrocytes n=3 | 1 | In one replicate mouse reads indicated contamination with mouse astrocytes (high Aqp4 and Gfap expression). Therefore mouse microglial reads from this replicate were excluded from analysis. | Microglia n=2<br>Astrocytes n=3 |
| | | WT microglia, WT astrocytes, WT neurons | Microglia n=3<br>Astrocytes n=3 | Nil | | Microglia n=3<br>Astrocytes n=3 |

**Supplementary Figure 1: Flow diagram of patient inclusion to cohort 1**
Related to figures 1 and 6

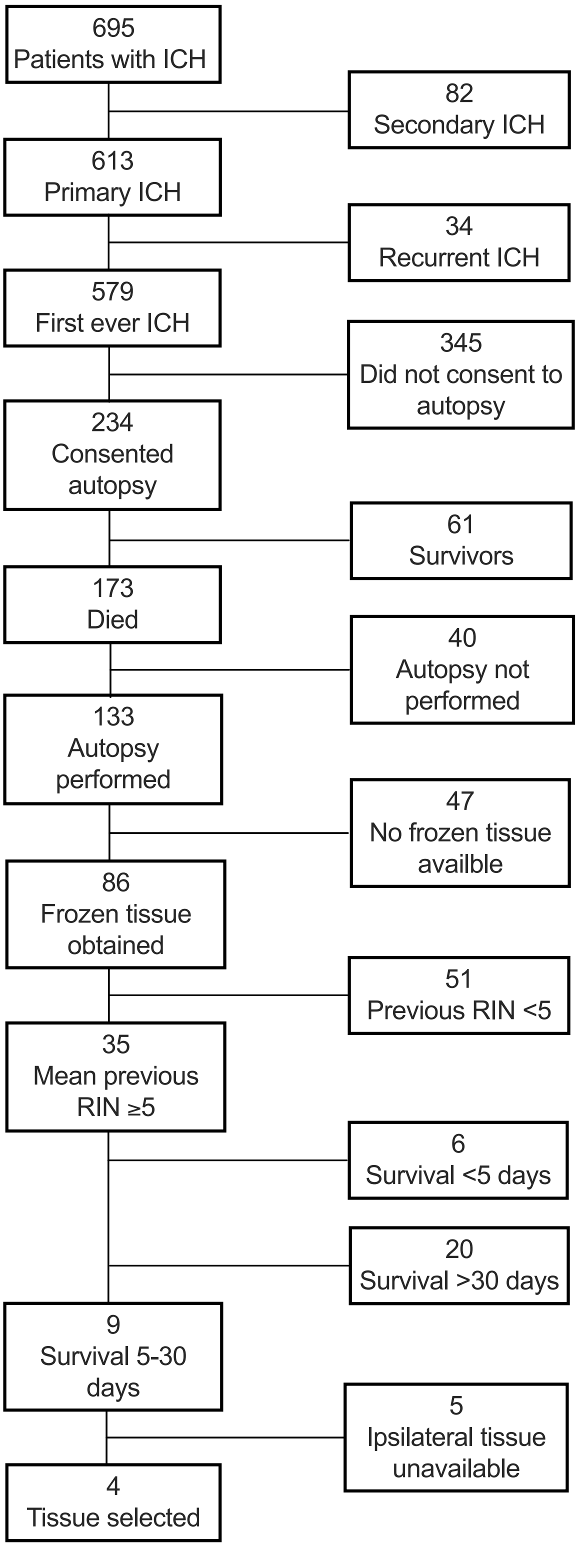

**Supplementary Figure 2: Flow diagram of patient inclusion to cohort 2**
Related to figure 1

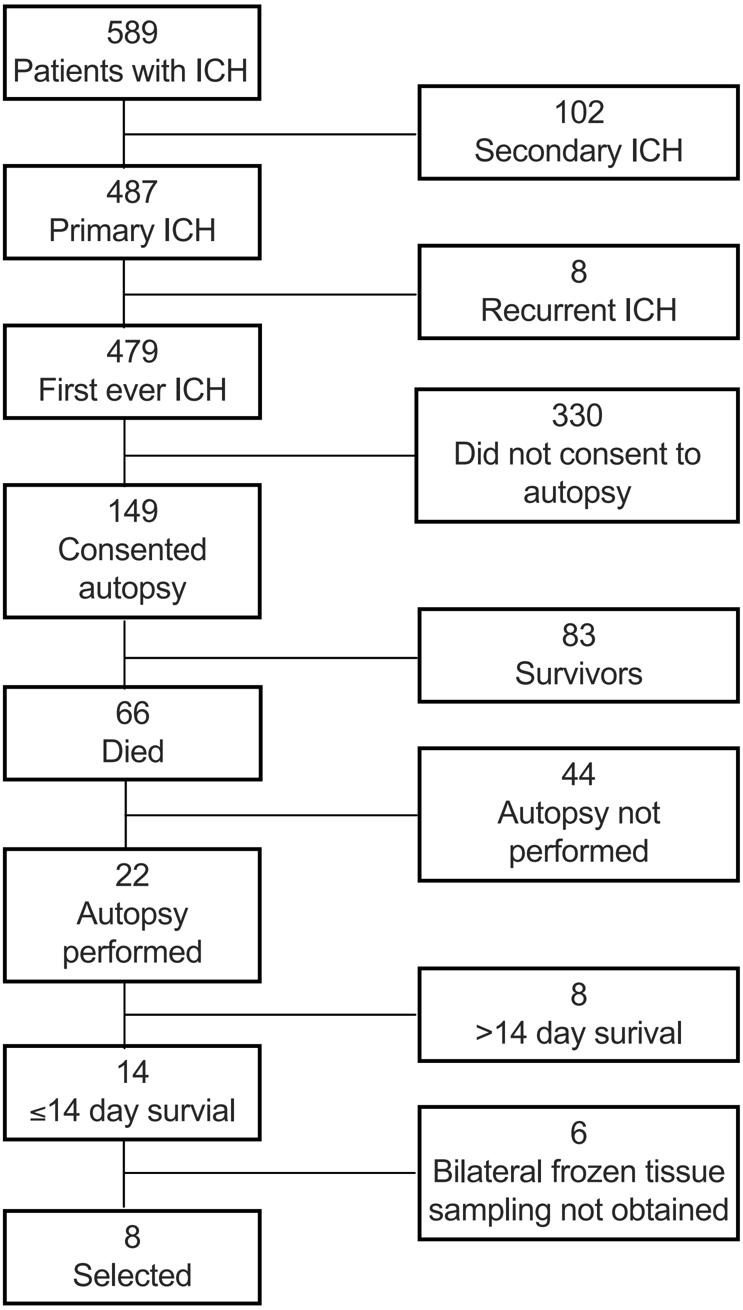

A

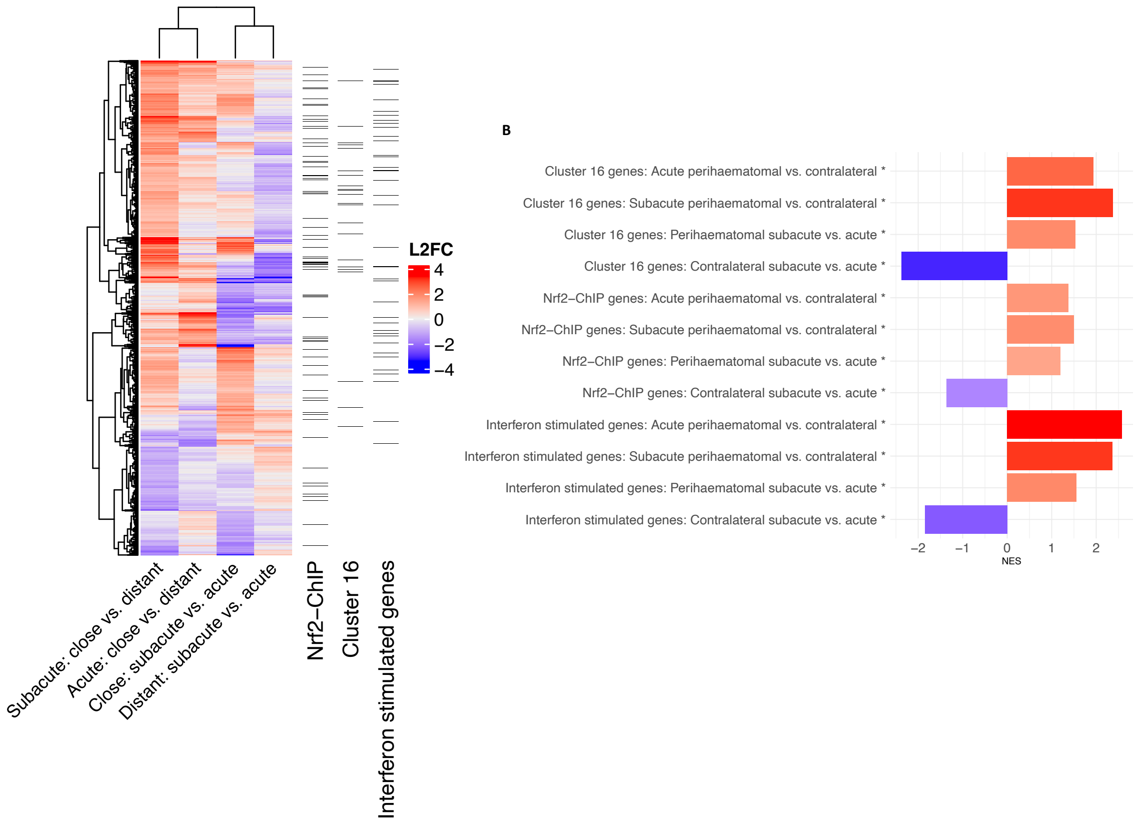

**Supplementary Figure 3: Spatiotemporal consequences of ICH in cohort 2 patients.**

Related to figure 1. (**A**) Heat map of $\log_2$ fold changes (L2FC) in expression of significantly differentially expressed genes (DEseq2 B-H $p_{adj}<0.05$) in brain tissue samples of patients (cohort 2) who died after ICH from perihaematomal (n=7) vs. contralateral (n=6) samples and patients who died within 48h after ICH symptom onset (acute; n=4) vs those who died 4-12 days after ICH symptom onset (subacute; n=4). Checks in right three columns highlight Nrf2 target genes (Nrf2 ChIP), those from cluster 16 (Figure 1D) and Hallmark interferon stimulated genes. (**B)** Bar blot of GSEA derived NES values for gene sets and comparisons shown in A. *All Benjamini-Hochberg padj<0.05.

**Supplementary Figure 4: Flow diagram of patient inclusion to cohort 3**
Related to figure 2.

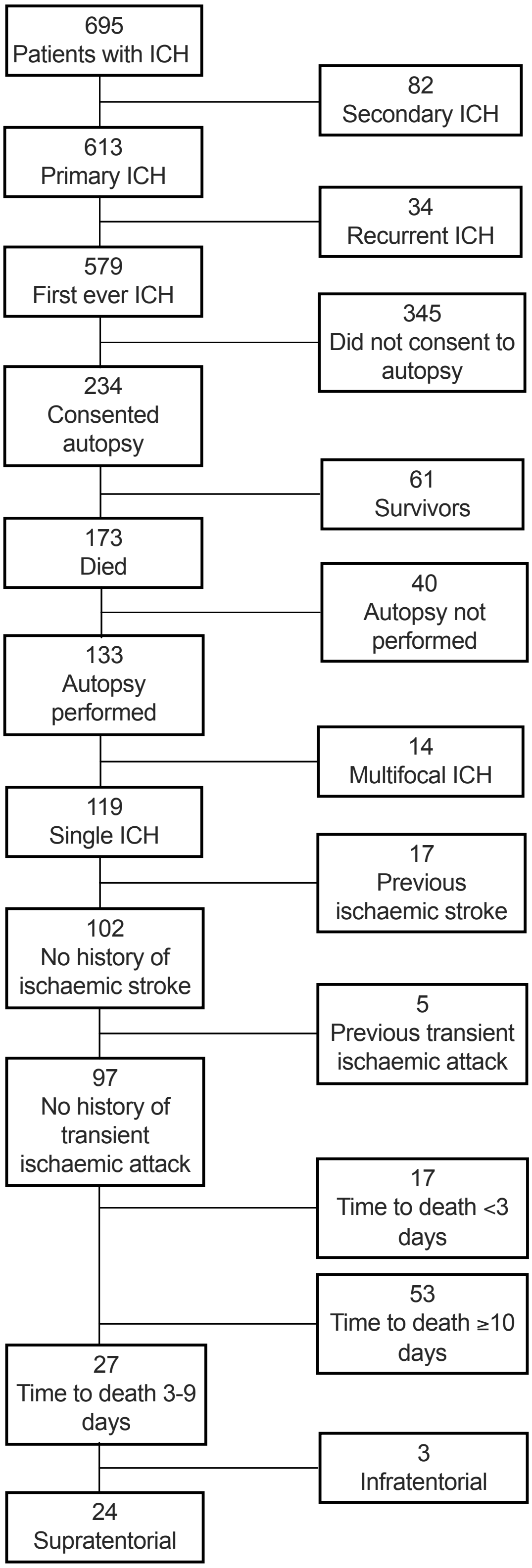

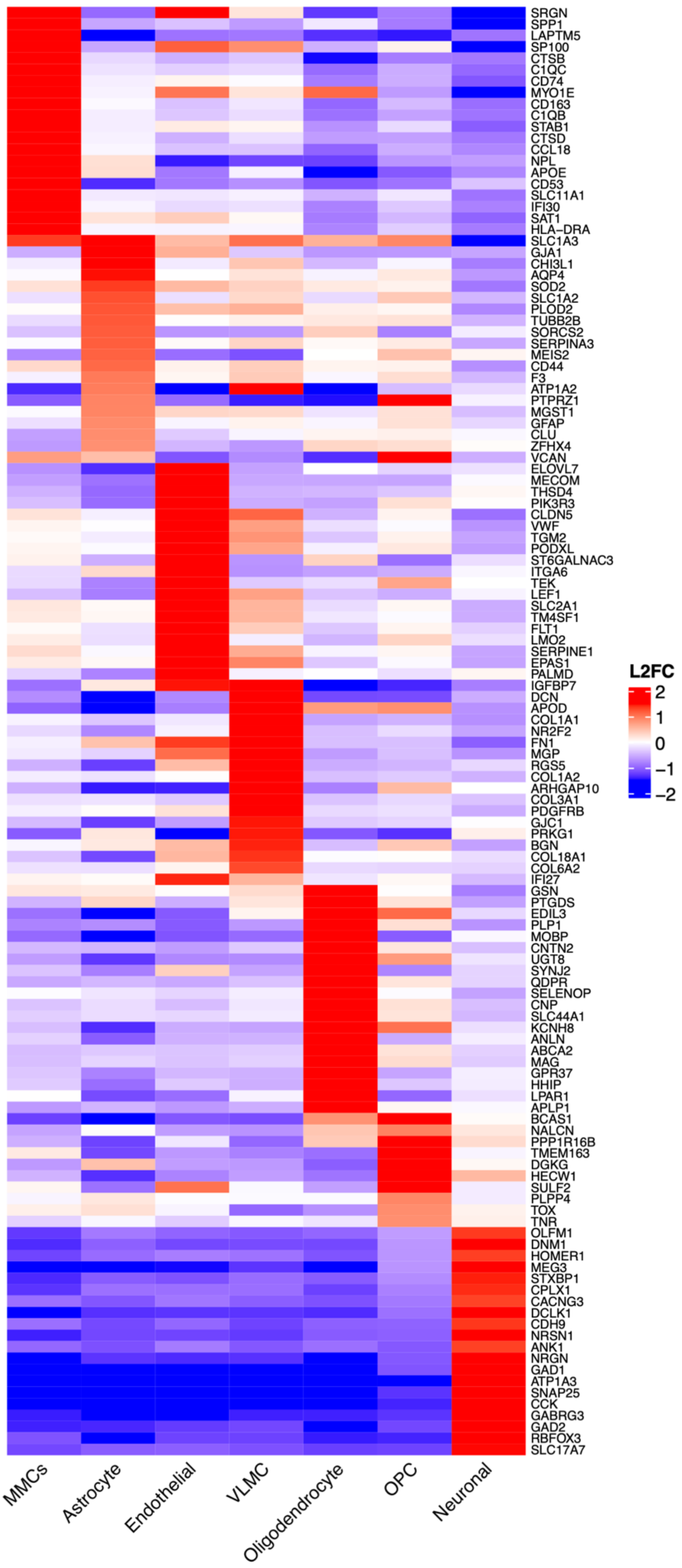

**Supplementary Figure 5: Cell-type marker genes in post-mortem brain tissue of patients after ICH or sudden non-neurological death.**

Related to figure 3. Heatmap of $\log_2$ fold changes (L2FC) of genes differentially expressed ($p_{adj}<0.05$) by one cell type compared with all other cell types. The 20 most highly specifically expressed genes per cell type shown where the gene expressed at least 0.5 L2FC greater than the next highest cell type and 0.8 L2FC versus all other cells combined. Possible cell type annotations derived from 10x Human MTG SEA-AD taxonomy (CCN20230505) include microglia or periventricular macrophages (Microglia_PVM, in our dataset this is synonymous with MMCs), astrocytes, endothelial cells, vascular leptomeningeal cells (VLMC), oligodendrocytes, oligodendrocyte progenitor cells (OPC) and neurons. N=4 patients/group: acute, chronic and controls

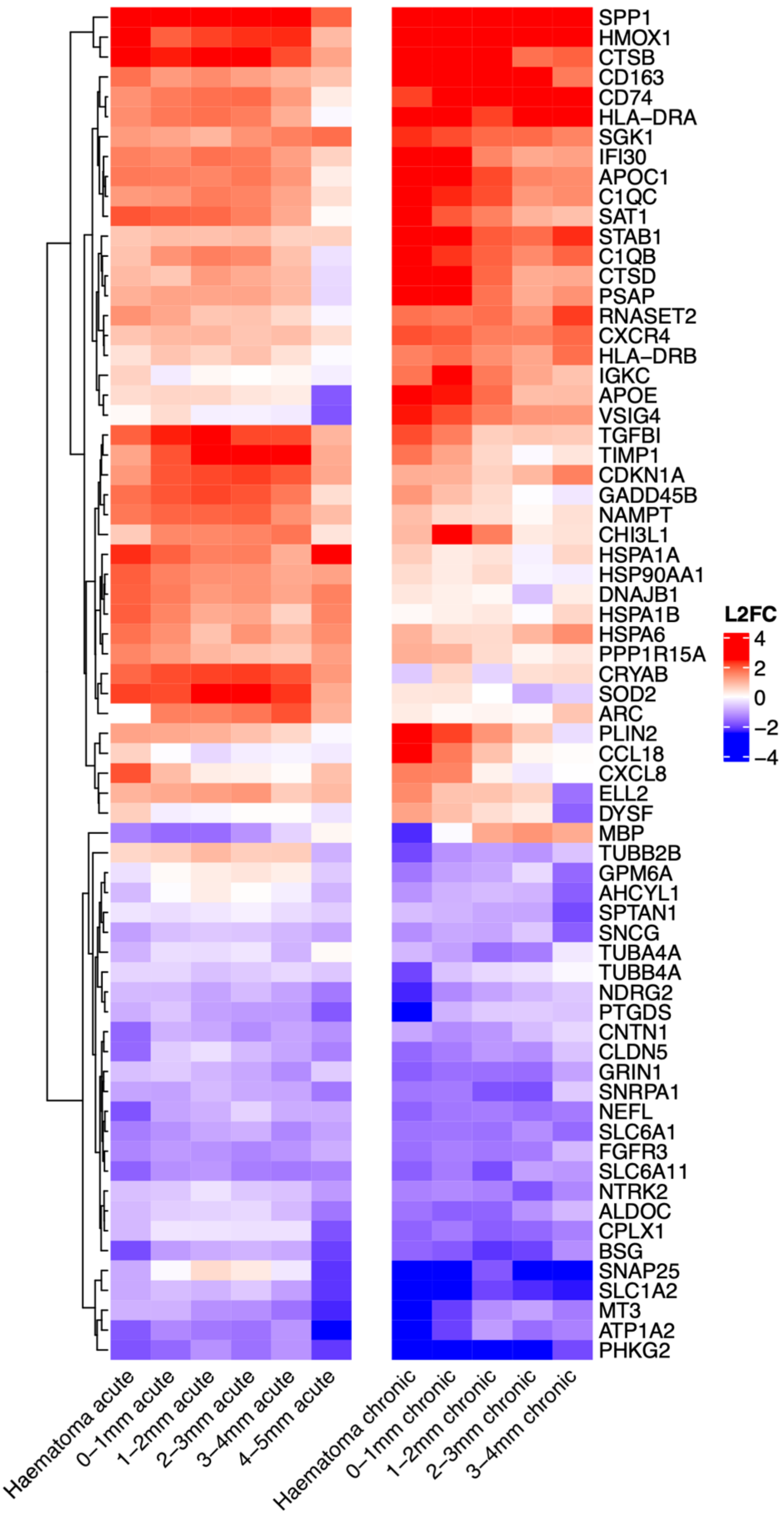

**Supplementary Figure 6: Differentially expressed genes by brain MMCs after ICH compared with sudden non-neurological death.**

Related to figure 3. Heatmap of $\log_2$ fold changes (L2FC) of genes differentially expressed ($p_{adj}<0.05$, L2FC>±1.5) by post-mortem brain MMCs in at least one distance bin from haematoma surface after either acute or chronic ICH compared with sudden non-neurological death, according to distance from the haematoma margin. N=4 patients/group: acute, chronic and controls. Rows clustered according to L2FC.

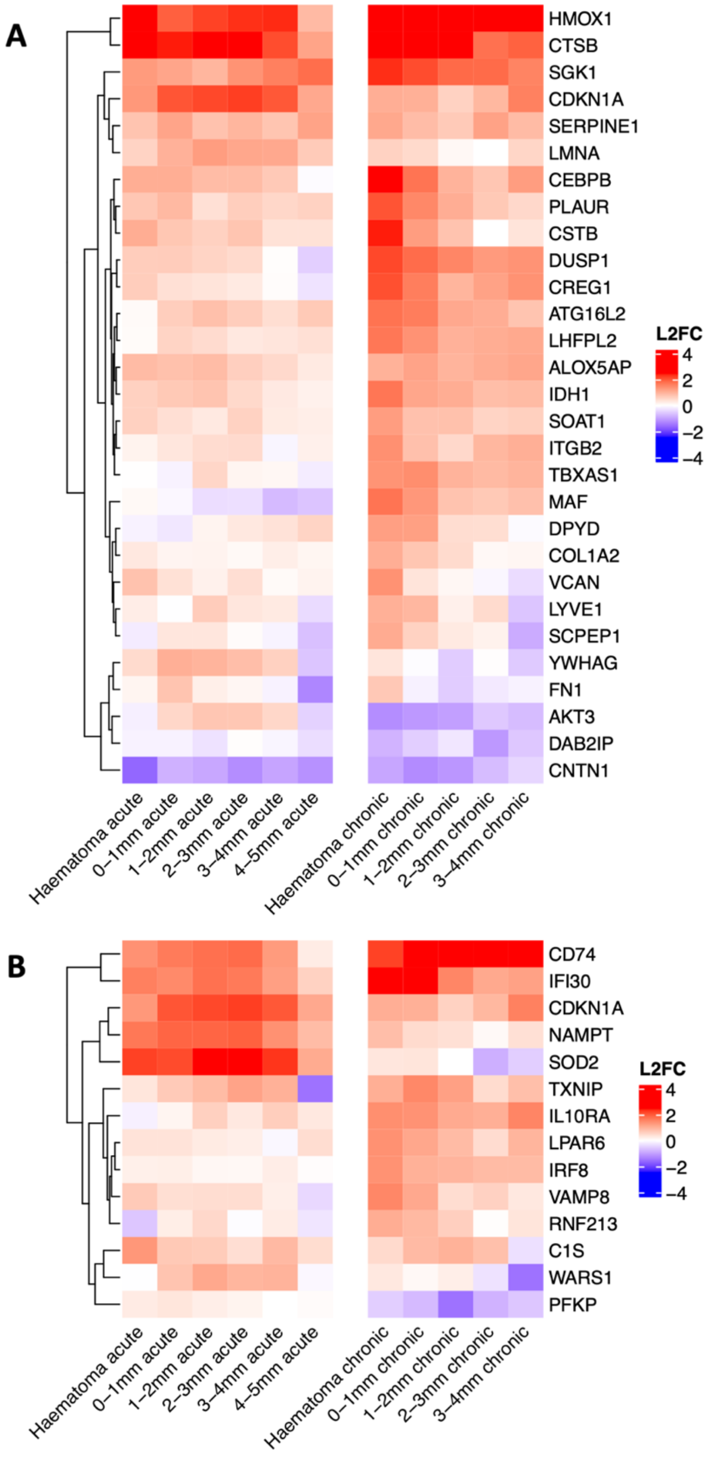

**Supplementary Figure 7: Differentially expressed Nrf2 target genes and ISGs by brain MMCs after ICH compared with sudden non-neurological death.**
Related to figure 3. Heatmap of $\log_2$ fold changes (L2FC) of Nrf2 target genes (**A**) or ISGs (**B**) differentially expressed ($p_{adj}<0.05$, L2FC>±1.0) by post-mortem brain MMCs in at least one distance bin from haematoma surface after either acute or chronic ICH compared with sudden non-neurological death, according to distance from the haematoma margin. N=4 patients/group: acute, chronic and controls. Rows clustered according to L2FC. Gene sets as previously annotated by an independent ChIP-seq experiment or the Hallmark interferon stimulated gene sets.

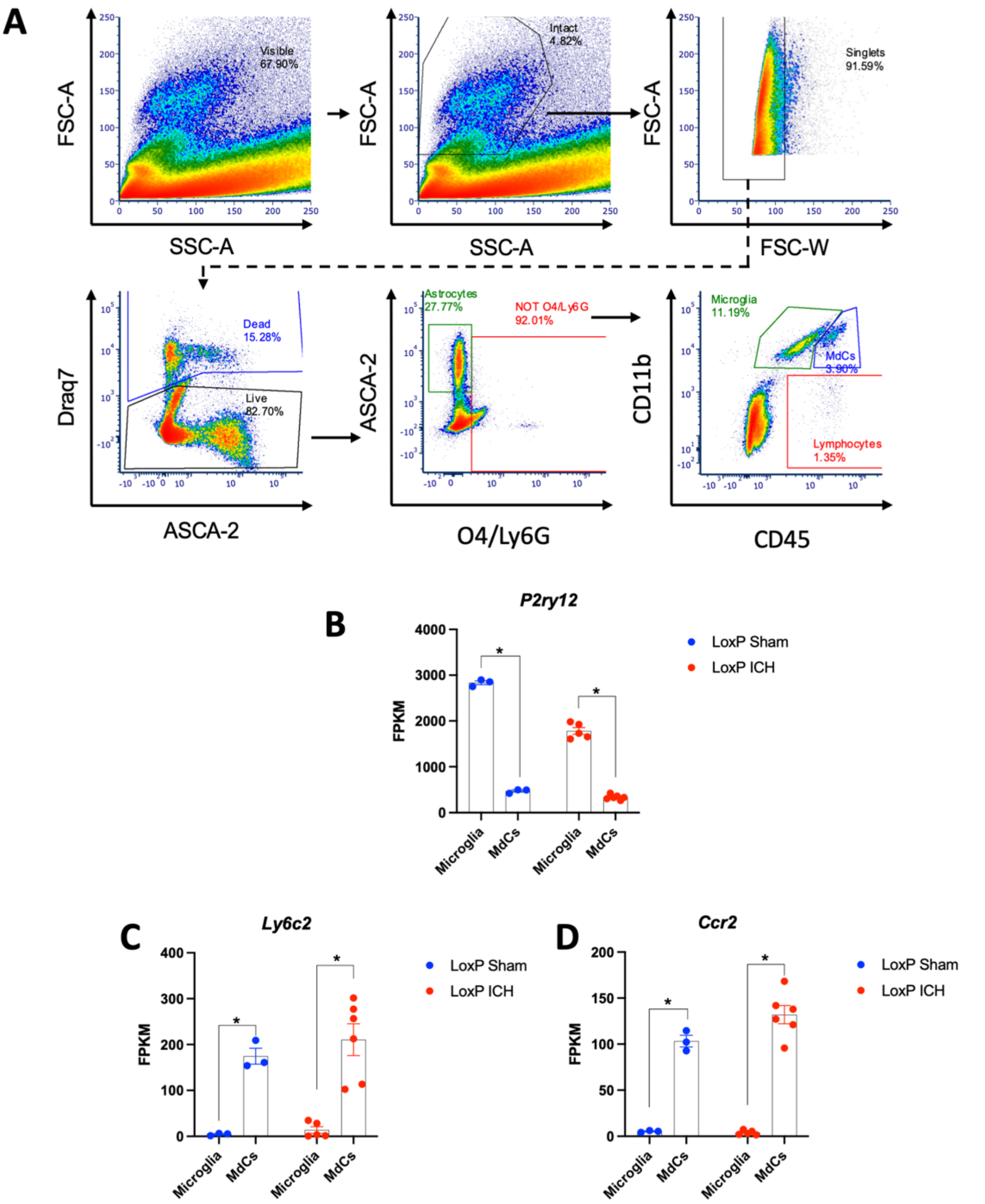

**Supplementary Figure 8: Fluorescence activated cell sorting.**

Related to figure 4. **A** FACS sorting strategy Forward scatter amplitude (FSC-A), width (FSC-W), side scatter amplitude (SSC-A) and Draq7 staining were used to identify intact, live, single cells. Microglia were defined as ASCA-2-, O4-, Ly6G-, CD11b+,$CD45^{int}$ and MdCs were ASCA-2-, O4-, Ly6G-, CD11b+,$CD45^{hi}$. **B-D** Microglial specific expression of *P2ry12* and MdC specific expression of *Ly6c2* and *Ccr2* confirm purity of sorting of sorted microglia and MdCs from mice after ICH (n=5 microglia, n=6 MdCs) or sham (n=3) surgery. **B** *P2ry12* expression in fragments per kilobase of transcript per million mapped reads (FPKM). DESeq2 Benjamini-Hochberg-adjusted p value ($p_{adj}$) for microglia versus MdCs in sham ($p_{adj}<0.0001$) or ICH ($p_{adj}<0.0001$). **C** *Ly6c* expression in FPKM for microglia versus MdCs in sham ($p_{adj}<0.0001$) or ICH ($p_{adj}<0.0001$). **D** *Ccr2* expression in FPKM for microglia versus MdCs in sham ($p_{adj}<0.0001$) or ICH ($p_{adj}<0.0001$).

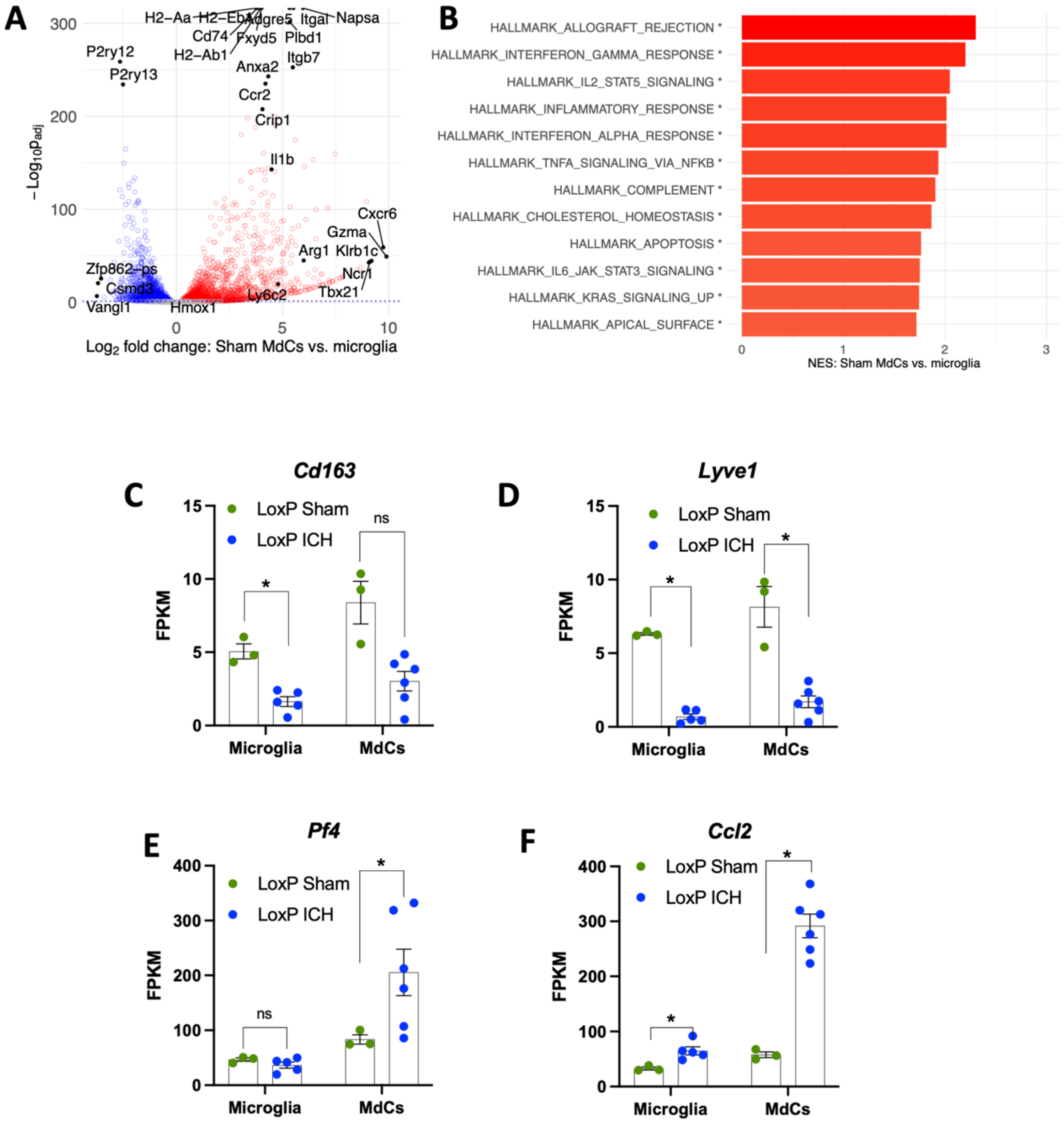

**Supplementary Figure 9: Sham MdCs exhibit distinct transcriptional phenotypes to sham microglia and MdCs after ICH.**

Related to figure 4. **A** Volcano plot of DESeq2 log2 fold differences in gene expression in microglia compared with MdCs after sham surgery. Red indicates higher expression in microglia, blue higher expression in MdCs, grey no detected difference in expression (DESeq2 Benjamini-Hochberg adjusted p ($p_{adj}$) ≥0.05) between both cell types. Clipped values at top of chart indicate genes where $p_{adj}$=0. **B** Bar plot of Gene Set Enrichment Analysis (GSEA) Normalised Enrichment Scores (NES) in MdCs compared with microglia after ICH for most highly enriched Hallmark Gene sets (*Benjamini-Hochberg adjusted p<0.05) **C-F** Expression of selected genes expressed by border associated macrophages, *Cd163* (**C**; microglia $p_{adj}$<0.001; MdCs $p_{adj}$=0.11) and *Lyve1* (**D**; microglia $p_{adj}$<0.001; MdCs $p_{adj}$=0.001) as well as those expressed by monocyte derived macrophages, *Pf4* (**E**; microglia $p_{adj}$=0.49; MdCs $p_{adj}$=0.011) and *Ccl2* (**F**; microglia $p_{adj}$<0.001; MdCs $p_{adj}$<0.001). Error bars ± SEM. Fragments Per Kilobase of transcript per Million mapped reads (FPKM). N=3 sham microglia/MdCs, 6 ICH microglia/MdCs.

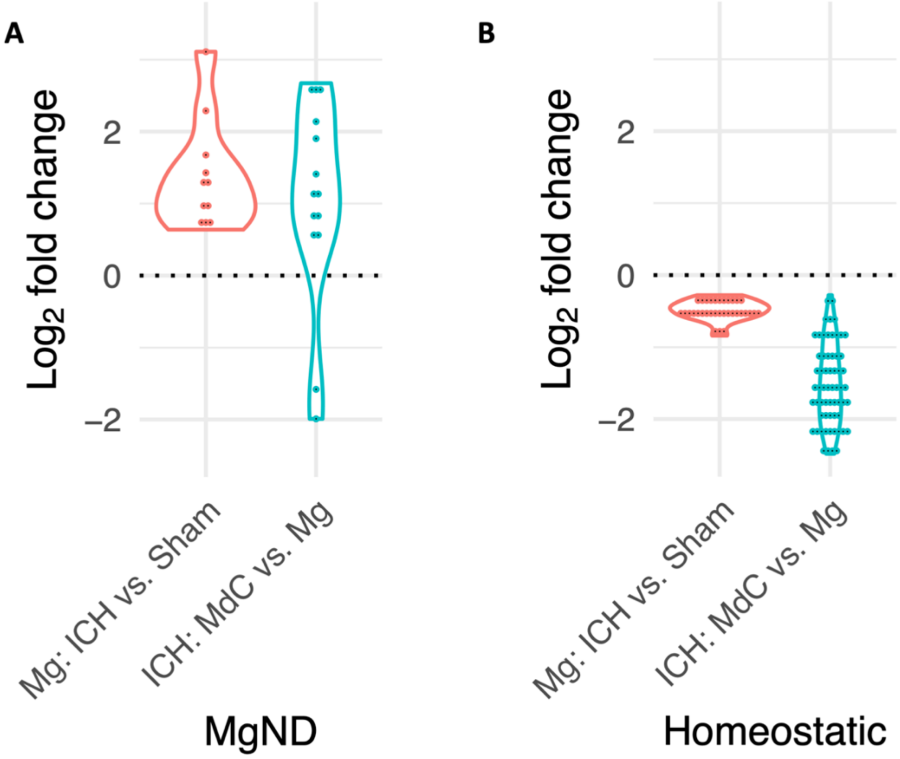

**Supplementary Figure 10: After ICH Microglia and MdCs adopt transcriptional signatures associated with neurodegenerative disease and repress homeostatic gene expression.**
Related to figure 4. Genes previously shown to be induced in microglia in neurodegenerative disease states (**A**; MgND) are increased in LoxP microglia after ICH vs. sham and in LoxP MdCs vs. microglia after ICH. Type 2 repeated-measures ANOVA ICH Microglia effect of ICH F(1-66)=55; p<0.0001. MdCs vs. Mg F(1-185)=90; p<0.0001. Conversely genes induced by homeostatic microglia (**B**) were repressed. Type 2 repeated-measures ANOVA ICH Microglia effect of ICH F(1-198)=96; p<0.0001. MdCs vs. Mg F(1-488)=911; p<0.0001. DEseq2 Log2 fold changes plotted. Only genes with DESeq2 Benjamini-Hochberg adjusted p<0.05 plotted. N=3 sham microglia/MdCs, 6 ICH microglia/MdCs. Gene sets defined according to Krasemann, et al. *Immunity* 2017; 47(3):566-581 (see main text for full citation).

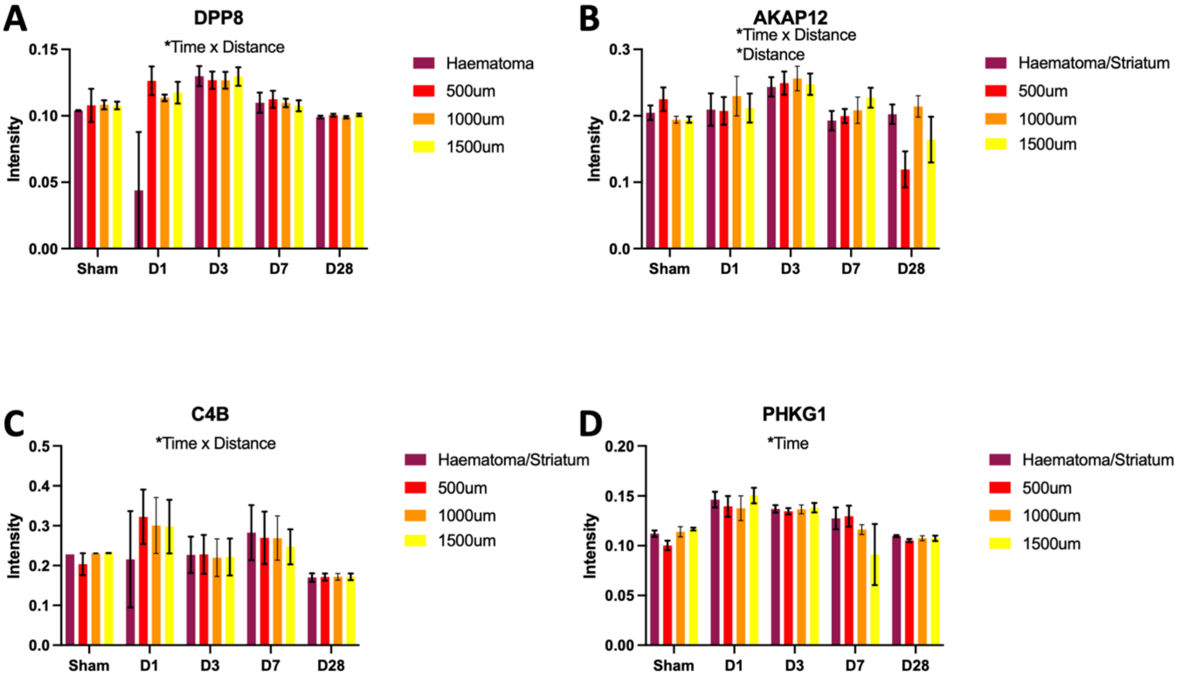

**Supplementary Figure 11: Temporal changes in protein expression after ICH in wild-type mice measured by mass spectrometry imaging.**

Related to figure 4. Selected potentially differentially expressed proteins ($p<0.05$) according to distance from haematoma in whole brain tissue of mice with myelomononuclear Nrf2 deficiency (KO) compared with LoxP control littermates. Selected proteins were induced after ICH and were highly expressed by glia: astrocytes (**A,D**) Endothelial cells (**B**), astrocytes and MMCs (**C**). Statistically significant findings in this analysis were not robust to correction for multiple testing. Sham data from day 3 shown for example was not included in statistical testing of effect of distance or time. **A** DPP8 time F(3-9)=2.0, p=0.18; distance F(3-39)=2.8, p=0.052; time*distance F(9-39)=4.9, p=0.00021. **B** AKAP12 time F(3-9)=0.30, p=0.82; distance F(3-39)=3.4, p=0.028; time*distance F(9-39)=3.0, p=0.0086. **C** C4B time F(3-9)=1.9, p=0.20; distance F(3-39)=1.1, p=0.38; time*distance F(3-39)=2.8, p=0.012. **D** PHKG1 time F(3-9)=10.0, p=0.0031; distance F(3-39)=0.50, p=0.68; time*distance F(3-39)=1.1, p=0.40. Mixed effects analysis with fixed effects of time, distance from haematoma and the interaction between time and distance. Random effects of subject and the slide each section was mounted on are also accounted for. D1 ICH n=3, d3 ICH n=7, d3 sham n=2, d7 ICH n=4, d28 ICH n=3.

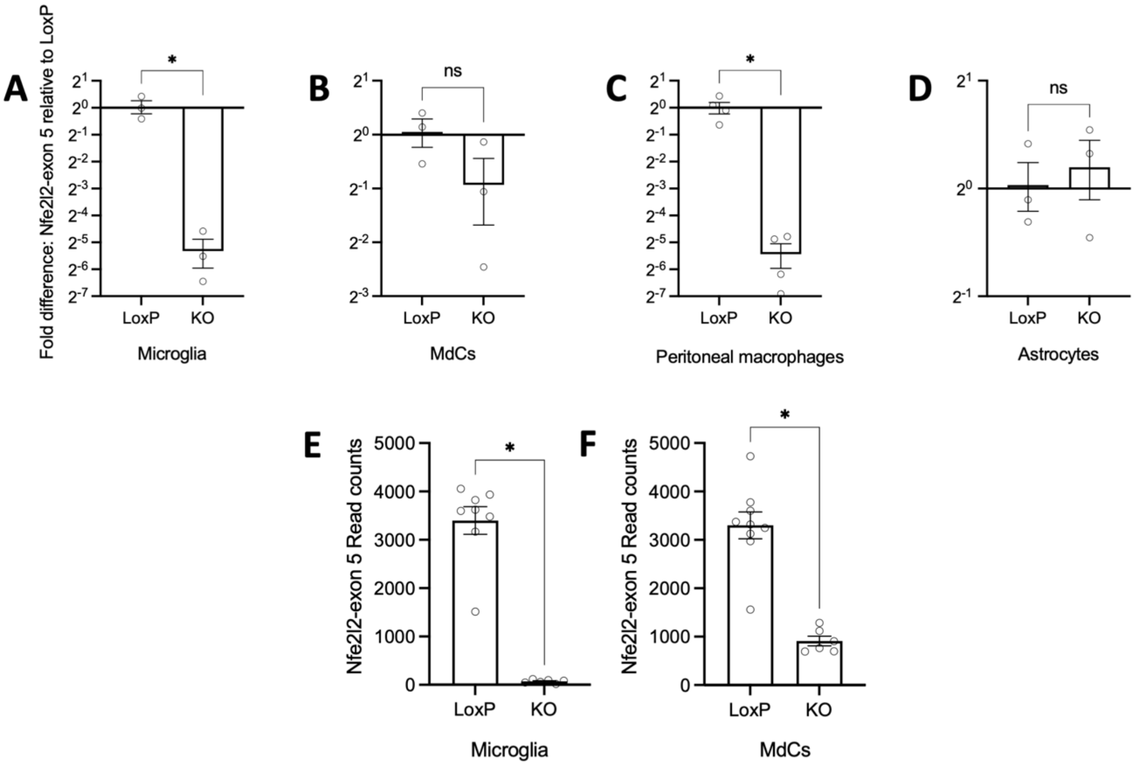

**Supplementary Figure 12: MMC restricted deletion of Nfe2l2 exon 5.**
Related to figure 5. Expression of Nfe2l2 exon 5 by microglia (**A)**, brain MdCs (**B**), peritoneal macrophages (**C**), and astrocytes (**D**) of LoxP and KO mice measured by RT-qPCR. Fold difference is relative to mean LoxP expression value. Microglia, MdCs and astrocytes n=3, peritoneal macrophages n=4. Two-tailed t-tests microglia t=5.9, df=4, p=0.0041; MdCs t=1.8, df=4, p=0.14; astrocytes t=0.46, df=4, p=0.67; peritoneal macrophages t=6.6, df=6, p=0.0006. Expression of Nfe2l2 exon 5 by microglia (**E**) and brain MdCs (**F**) measured as read counts from RNA sequencing data. Microglia LoxP n=8, KO n=6. Two-tailed t-test t=10.0, df=12, p<0.0001. MdCs LoxP n=9, KO n=6. Two-tailed t-test t=6.8, df=13, p<0.0001.

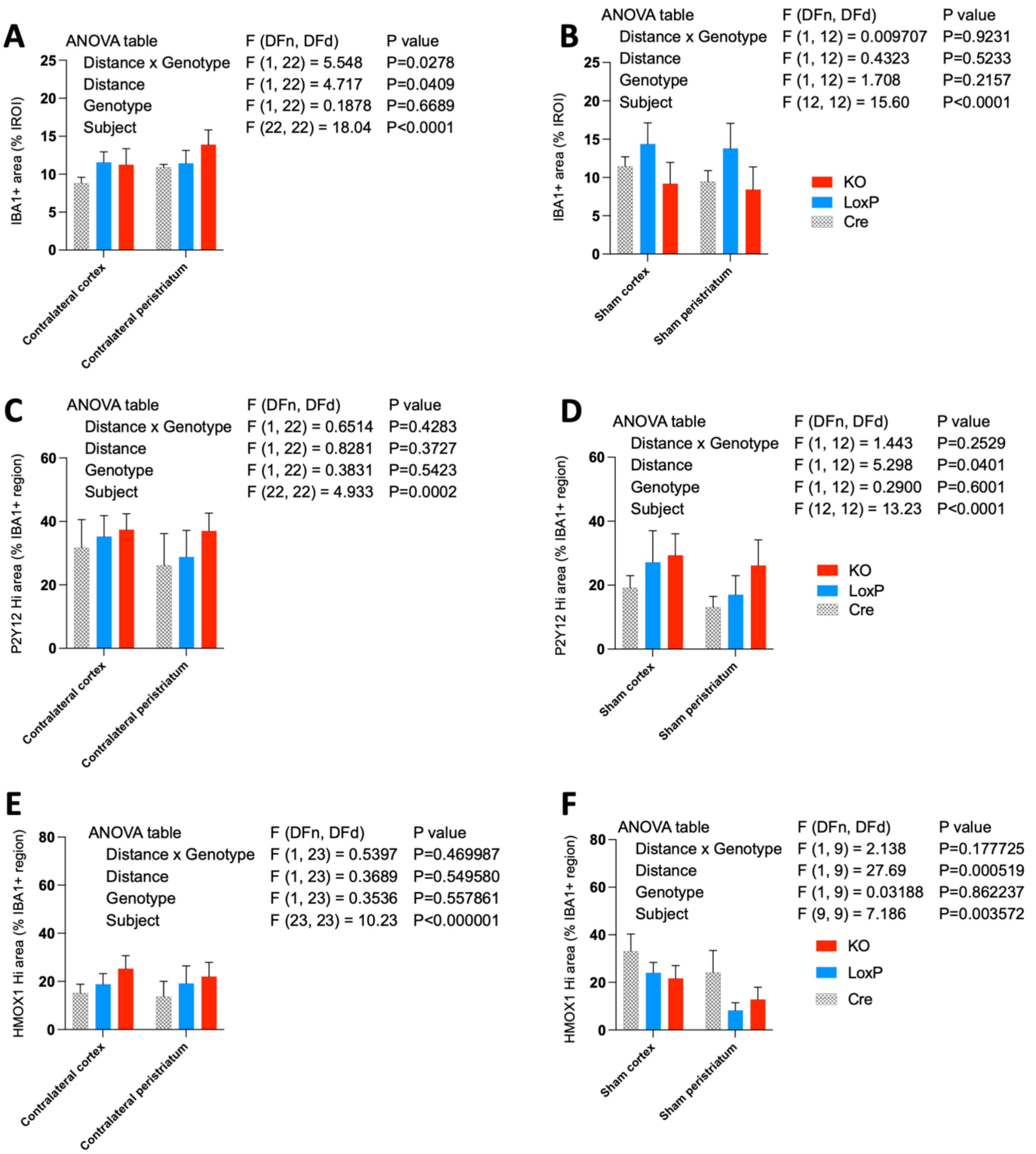

**Supplementary Figure 13: P2Y12 and HMOX1 expression are similar outside the perihaematomal region of LoxP and KO mice.**

Related to figure 5. Immunofluorescence staining from regions contralateral to ICH (**A, C, E**) and of sham operated mice (**B,D,F**). Cortical and peristriatal regions were manually defined for each mouse at the haematoma or sham injection level. Comparisons made between KO and loxP mice with comparable values for Cre mice also provided. Two-way ANOVA tables presented on graphs. Post-hoc multiple comparison p-values adjusted using the Holm-Šídák method. **A-B** IBA1 staining expressed as a percentage of each ROI (KO versus LoxP **A**: cortex $p_{adj}$=0.91; peristriatum $p_{adj}$=0.57. **B**: cortex $p_{adj}$= 0.37; peristriatum $p_{adj}$=0.37). **C-D** P2Y12 stained area as a percentage of the total IBA1 stained area (KO versus LoxP **C**: cortex $p_{adj}$=0.82; peristriatum $p_{adj}$=0.61. **D**: cortex $p_{adj}$= 0.84; peristriatum $p_{adj}$=0.65). **E-F** HMOX1 stained area as a percentage of the total IBA1 stained area (KO versus LoxP **E**: cortex $p_{adj}$=0.68; peristriatum $p_{adj}$=0.73. **F**: cortex $p_{adj}$= 0.76; peristriatum $p_{adj}$=0.76). All error bars +/- SEM. ICH loxP and KO n=12/group, cre and sham n=7/group.

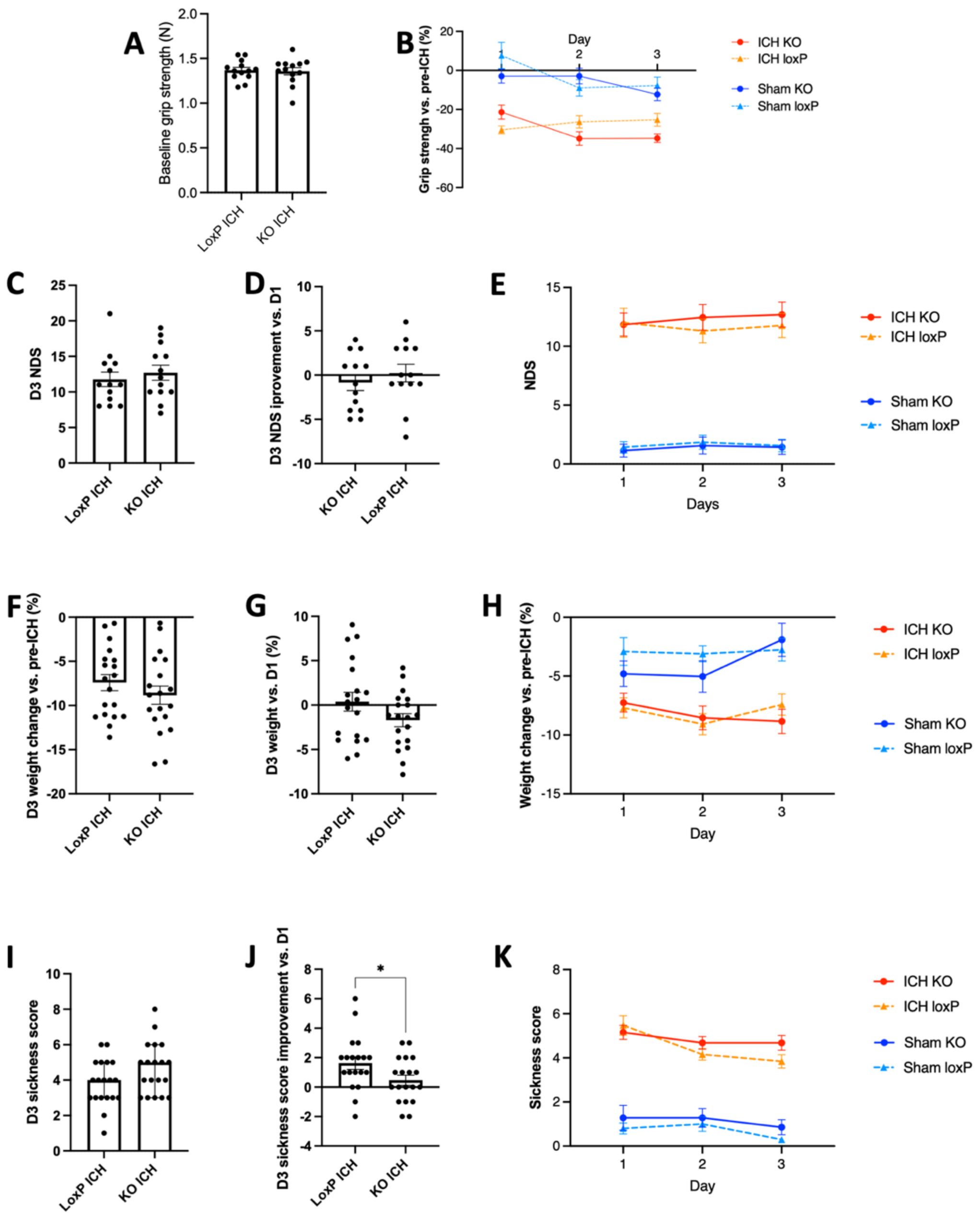

**Supplementary Figure 14: Additional behavioural consequences of Nrf2 deletion in mononuclear myeloid cells.**

Related to figure 5. Secondary behavioural outcome measures. Continuous data were analysed by Welch's corrected two-tailed t-test and discrete data by two-tailed Mann-Whitney U. **A** Baseline pre-ICH grip strength in Newtons (N; t=0.24, df=22.3; p=0.81). **B** Plot of mean percentage of daily grip strength versus pre-ICH/sham surgery. **C-E** Clark's Neurological Deficit Score (NDS) at day three (**C**; U=72; p=0.53), day three versus day one (**D**; U=69.5; p=0.45), and mean daily NDS (**E**). **F-H** Percentage weight change at day three versus pre-ICH (**F**; t=1.0; df=35.4; p=0.30), day three versus day one (**G**; t=1.6; df=32.1; p=0.11), and mean daily weight change versus pre-ICH (**H**). **i-K** General sickness score at day three (**I**; U=127; p=0.12), improvement in sickness score at day three versus day one (**J**; U=114.5; p=0.0498) and mean daily sickness score (**K**). Error bars ± SEM, Grip and NDS n=13 loxP and KO ICH; all other groups n=7. Weight and sickness score n=19 loxP ICH and KO ICH; n=10 loxP sham; n=7 KO sham.

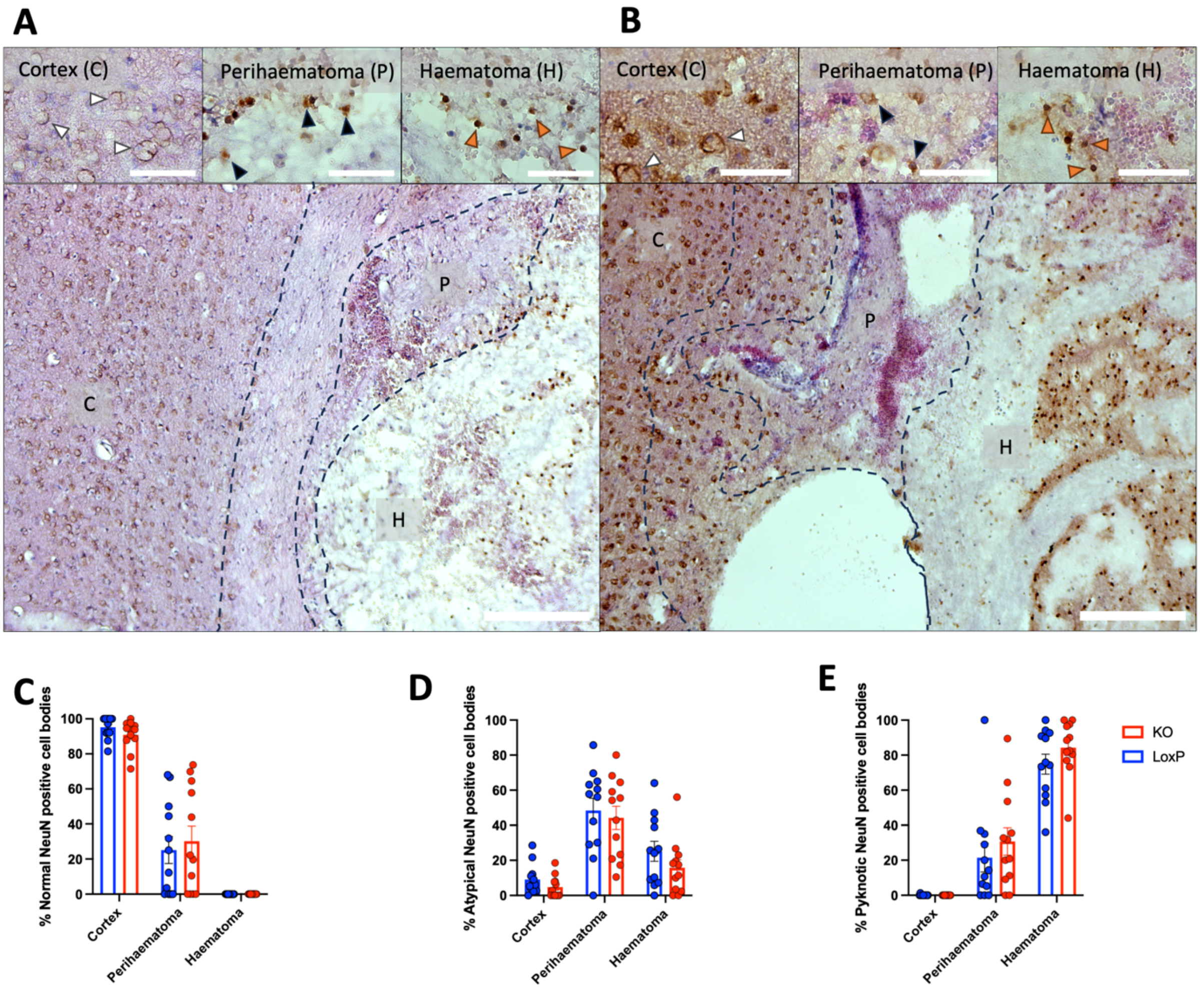

| ANOVA table | F (DFn, DFd) | P value |
|---|---|---|
| Region x Genotype | F (2, 44) = 0.4445 | P=0.6440 |
| Region | F (2, 44) = 183.9 | P<0.0001 |
| Genotype | F (1, 22) = 0.003695 | P=0.9521 |
| Subject | F (22, 44) = 0.8615 | P=0.6390 |

| ANOVA table | F (DFn, DFd) | P value |
|---|---|---|
| Region x Genotype | F (2, 44) = 0.1904 | P=0.8273 |
| Region | F (2, 44) = 34.78 | P<0.0001 |
| Genotype | F (1, 22) = 1.768 | P=0.1973 |
| Subject | F (22, 44) = 1.353 | P=0.1935 |

| ANOVA table | F (DFn, DFd) | P value |
|---|---|---|
| Region x Genotype | F (2, 44) = 0.4610 | P=0.6336 |
| Region | F (2, 44) = 101.3 | P<0.0001 |
| Genotype | F (1, 22) = 2.307 | P=0.1430 |
| Subject | F (22, 44) = 0.7724 | P=0.7396 |

**Supplementary Figure 15: Neuronal pyknosis after ICH in NeuN-H&E stained mouse brain tissue.**

Related to figure 5. LoxP (**A**) and KO (**B**) tissue stained with DAB NeuN and H&E. Representative images show cortical, haematomal and perihaematomal regions of interest. Arrows indicate normal (white), atypical (black) and pyknotic NeuN positive nuclear morphologies. Scale bars 200µm, insert 50µm. Quantification of percentage normal (**C**), atypical (**D**) and pyknotic (**E**) NeuN positive neuronal cell bodies in LoxP (blue) and KO (red) mice. Two-way ANOVA tables provided. N=12/genotype. All post-hoc comparisons p>0.05

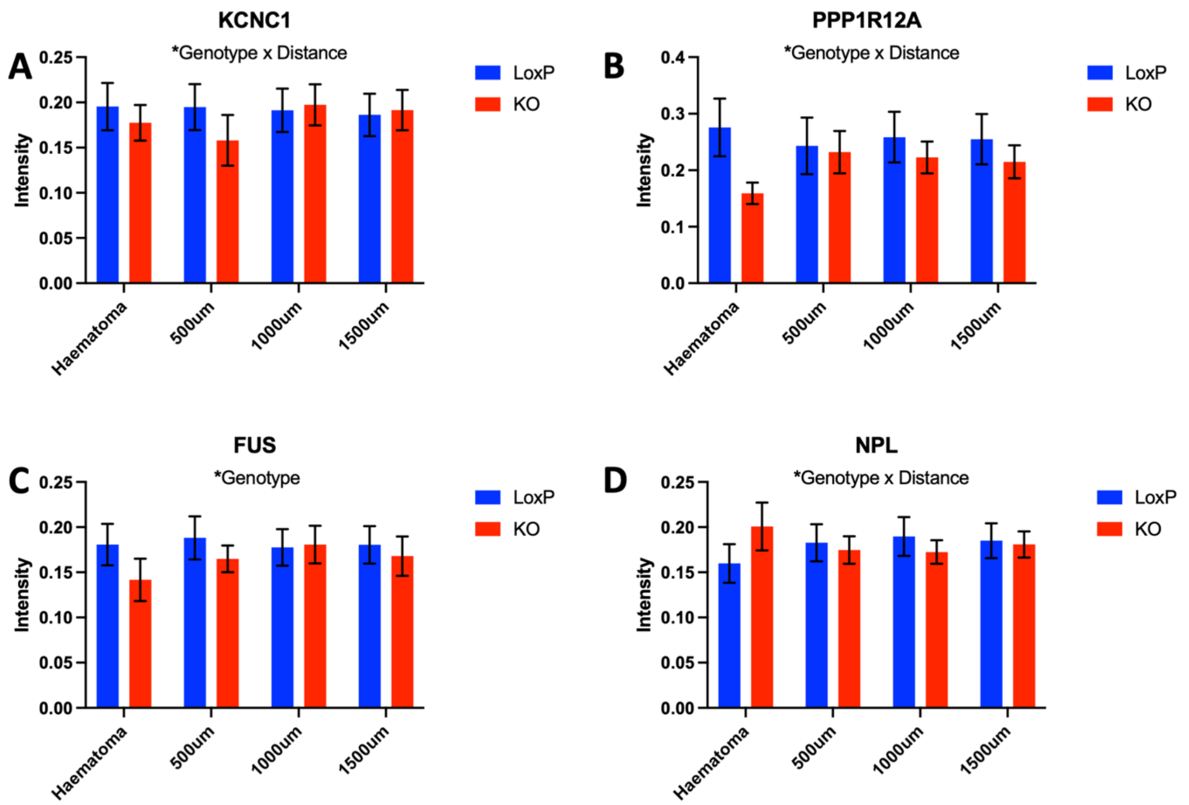

**Supplementary Figure 16: Protein expression in mice with myelomononuclear Nrf2 deficiency measured by mass spectrometry imaging.**

Related to figure 6. Selected potentially differentially expressed proteins ($p<0.05$) according to distance from haematoma in whole brain tissue of mice with myelomononuclear Nrf2 deficiency (KO) compared with LoxP control littermates. Selected proteins are highly expressed by neurons and astrocytes (**A-C**) or MMCs (**D**). Statistically significant findings in this analysis were not robust to correction for multiple testing. **A** KCNC1 genotype $F(1\text{-}17)=2.22$, $p=0.15$; distance $F(3\text{-}72)=1.9$, $p=0.13$; genotype*distance $F(3\text{-}72)=3.7$, $p=0.016$**.** **B** PPP1R12A genotype $F(1\text{-}17)=0.86$, $p=0.37$; distance $F(3\text{-}72)=0.74$, $p=0.53$; genotype*distance $F(3\text{-}72)=3.6$; $p=0.018$. **C** FUS genotype $F(1\text{-}17)=5.6$, $p=0.029$; distance $F(3\text{-}72)=1.1$, $p=0.36$; genotype*distance $F(3\text{-}72)=1.3$, $p=0.27$. **D** NPL genotype $F(1\text{-}17)=1.1$, $p=0.32$; distance $F(3\text{-}72)=0.05$, $p=0.98$; genotype*distance $F(3\text{-}72)=2.9$, $p=0.039$. Mixed effects analysis with fixed effects of genotype, distance from haematoma and the interaction between genotype and distance. Random effects of subject and the slide each section was mounted on are also accounted for. N=13/genotype.

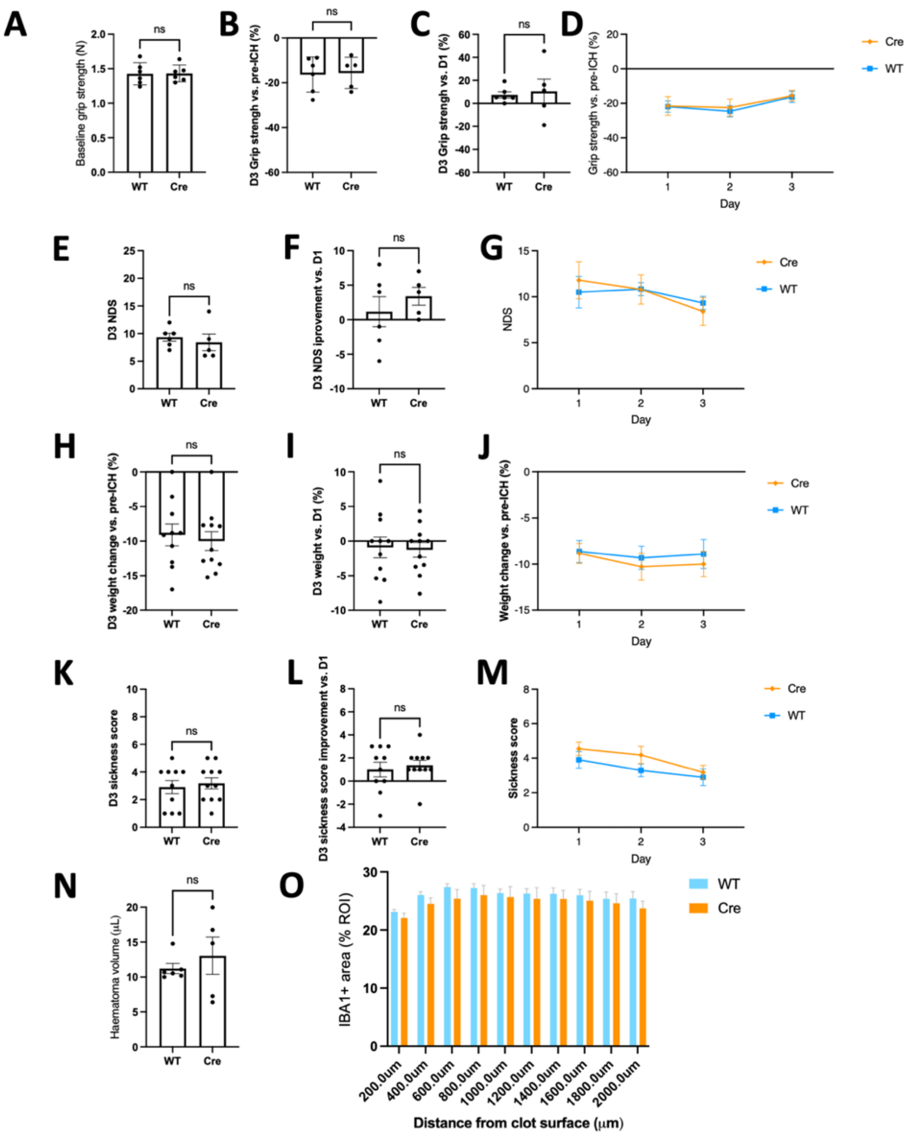

| ANOVA table | F (DFn, DFd) | P value |
|---|---|---|
| Distance x Genotype | F (9, 81) = 0.3540 | P=0.9532 |
| Distance | F (1.144, 10.30) = 9.417 | P=0.0098 |
| Genotype | F (1, 9) = 0.6067 | P=0.4560 |
| Subject | F (9, 81) = 39.69 | P<0.0001 |

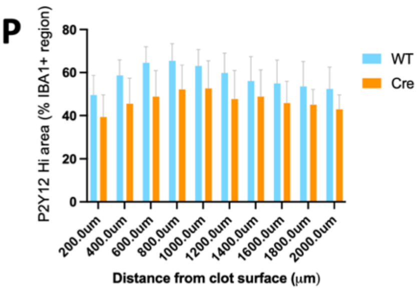

| ANOVA table | F (DFn, DFd) | P value |
|---|---|---|
| Distance x Genotype | F (9, 81) = 0.1922 | P=0.9944 |
| Distance | F (1.203, 10.83) = 2.419 | P=0.1466 |
| Genotype | F (1, 9) = 0.6838 | P=0.4297 |
| Subject | F (9, 81) = 49.84 | P<0.0001 |

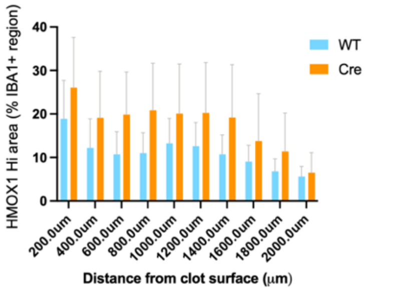

| ANOVA table | F (DFn, DFd) | P value |
|---|---|---|
| Distance x Genotype | F (9, 81) = 0.3678 | P=0.9472 |
| Distance | F (1.188, 10.69) = 4.389 | P=0.0559 |
| Genotype | F (1, 9) = 0.4127 | P=0.5366 |
| Subject | F (9, 81) = 56.33 | P<0.0001 |

**Supplementary Figure 17: Cx3cr1 haploinsufficiency and heterozygous Cre recombinase expression do not influence behavioural or histological outcome after ICH.**

Related to figure 6. Comparison of outcomes in C57Bl6J-Nfe2l2$^{wt/wt}$-Cx3cr1$^{cre/wt}$ (Cre) and C57Bl6J-Nfe2l2$^{wt/wt}$-Cx3cr1$^{wt/wt}$ (Wild type; WT) mice after ICH. **A** Baseline pre-

ICH grip strength in Newtons (N; Welch's t-test; t=0.07; df=9.3; p=0.95). **B-D** Day three grip strength percentage of pre-ICH grip strength (**B**; Welch's t-test; t=0.16; df=8.9; p=0.88), day three grip strength percentage of day one grip strength (**C**; Welch's t-test; t=0.27; df=4.5; p=0.80), mean percentage of daily grip strength versus pre-ICH (**D**). **E-G** Clark's Neurological Deficit Score (NDS) at day three (**E**; Mann Whitney U; U=9; p=0.31), day three versus day one (**F**; Mann Whitney U; U=11; p=0.50), and mean daily NDS (**G**). **H-J** Percentage weight change at day three versus pre-ICH (**H**; Welch's t-test; t=0.43; df=18.2; p=0.67), day three versus day one (**I**; Welch's t-test; t=0.19; df=18.0; p=0.85), and mean daily weight change versus pre-ICH (**J**). **K-M** General sickness score at day three (**K**; Mann Whitney U; U=49; p=0.70), improvement in sickness score at day three versus day one (**L**; Mann Whitney U; U=52.5; p=0.87) and mean daily sickness score (**M**). **N** Haematoma volume at day three post-ICH (Welch's t-test t=0.66; df=4.6; p=0.54). **O** IBA1 positively stained area in regions of interest (ROI) of varying distance from haematoma (clot) surface. Percentage of each region of interest stained for IBA1 shown. **P** P2Y12 highly stained area within IBA1 positive regions at different distances from haematoma surface. Percentage of the total IBA1 stained area co-staining for P2Y12 for each region of interest shown. **Q** HMOX1 highly stained area within IBA1 positive regions at different distances from haematoma surface. Percentage of the total IBA1 stained area co-staining for HMOX1 for each region of interest shown. **O-Q** Two-way ANOVA tables shown. Error bars ± SEM. Weight and sickness score n=10 WT, n=11 cre, baseline grip strength n= 6 WT and cre, all others n = 6 WT, n=5 cre.

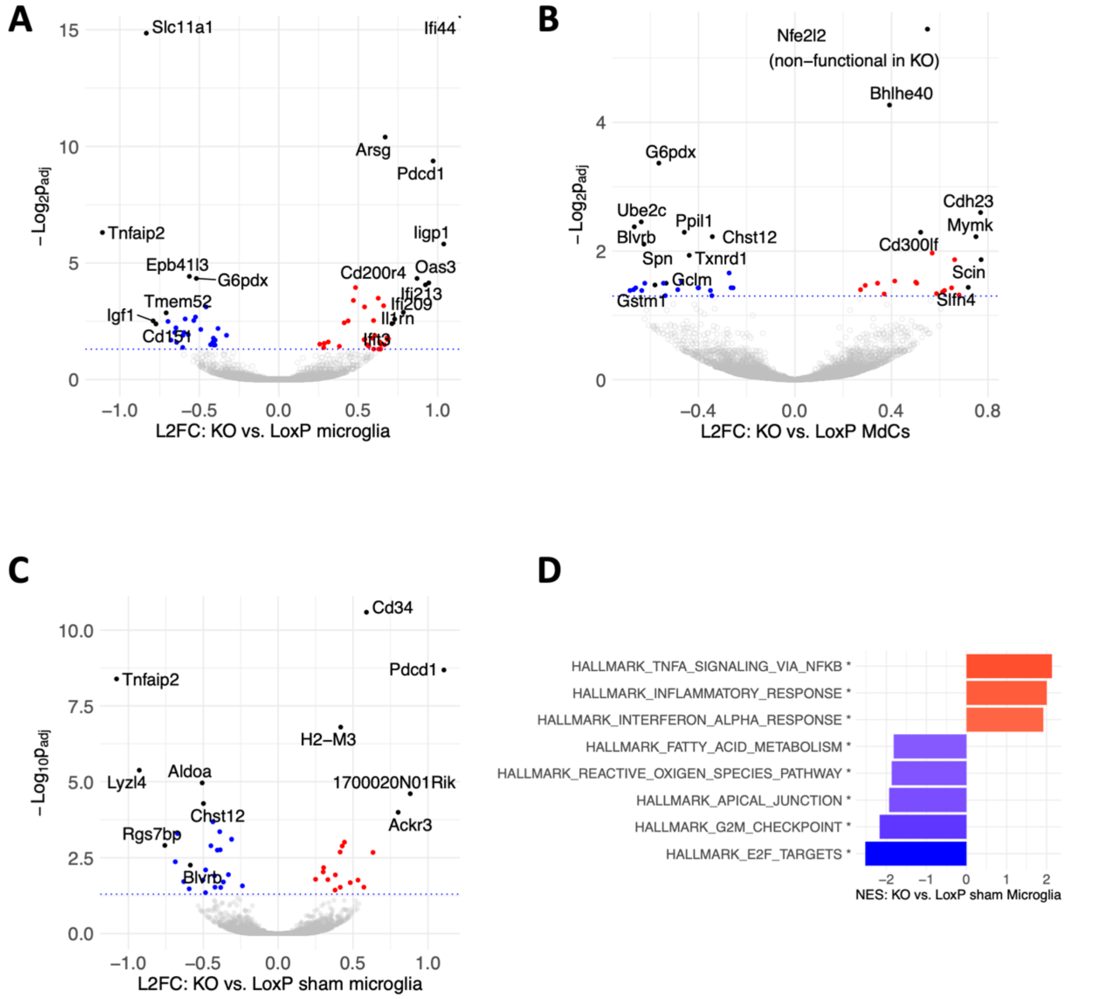

**Supplementary Figure 18: Effects of Nrf2 deletion on the microglial and MdC transcriptome.**

Related to figure 6. Volcano plots of DESeq2 log2 fold changes in gene expression and Benjamini-Hochberg adjusted p-values ($p_{adj}$) in the transcriptomes of FACS sorted microglia (**A,C**) and MdCs (**B**). The effect mononuclear myeloid cell Nrf2 deletion on cells isolated from LoxP or KO mice with ICH (**A,B**) or following sham surgery (**C**) are shown. Of note, the expression of Nfe2l2 mRNA (which encodes NRF2), was increased in KO compared with LoxP mice, which may reflect altered turnover of non-functional Nfe2l2 mRNA lacking exon 5. Red indicates higher expression in KO mice, blue higher expression in LoxP mice and grey no detected difference in expression ($p_{adj}$≥0.05) between both cell types. **D** Bar plots of Gene Set Enrichment Analysis (GSEA) Normalised Enrichment Scores (NES) for the five most highly positively and negatively enriched ($p_{adj}$<0.05) Hallmark gene sets in KO vs. LoxP microglia from mice following sham surgery. N= 6 ICH loxP microglia/MdCs, 5 ICH KO microglia, and 3 sham microglia/MdCs. Data for panels **C-D** from independent experiment with n=6 KO/loxP microglia.

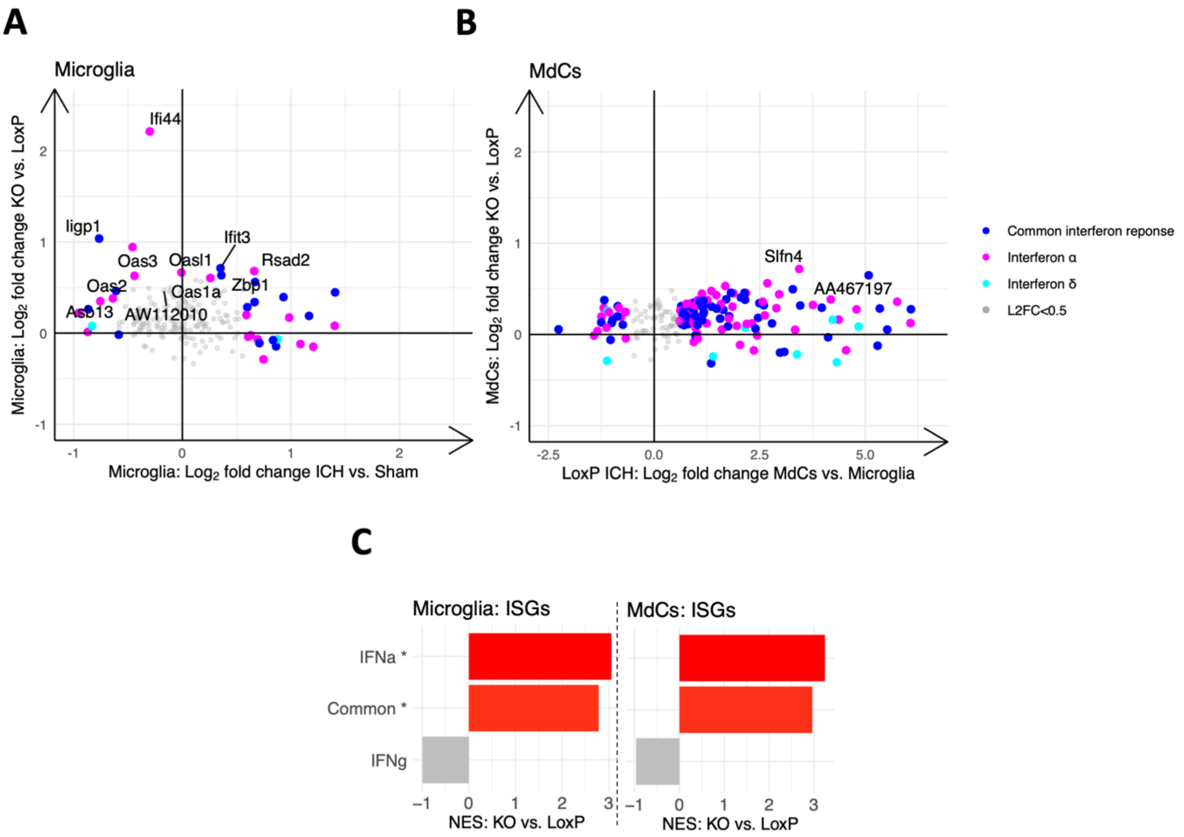

**Supplementary Figure 19: Effects of Nrf2 deletion on type-specific interferon-stimulated gene expression.**

Related to figure 6. **A-B** Scatter plot of L2FC in expression of interferon-α (IFNA; magenta), interferon-$\gamma$ (IFNG; cyan), and common (both α and $\gamma$; blue) stimulated genes of KO vs. LoxP after ICH (y-axis) and ICH vs. Sham LoxP (x-axis) microglia (**A**) and MdCs (**B**).[11] Genes with L2FC>1.5 in any direction or axis are coloured, those that are differentially expressed ($p_{adj}<0.05$) in KO vs. LoxP are labelled. **C** Enrichment plot of the above IFNA, IFNG and common ISG gene sets in KO vs. LoxP microglia and MdCs. Statistically significantly (*$p_{adj}<0.05$) positively enriched gene sets indicated in red, those that are not statistically significantly positively or negatively enriched are grey. N=6 ICH loxP microglia/MdCs, and 5 ICH KO microglia, 6 ICH KO MdCs, 3 sham KO/loxP microglia/MdCs.

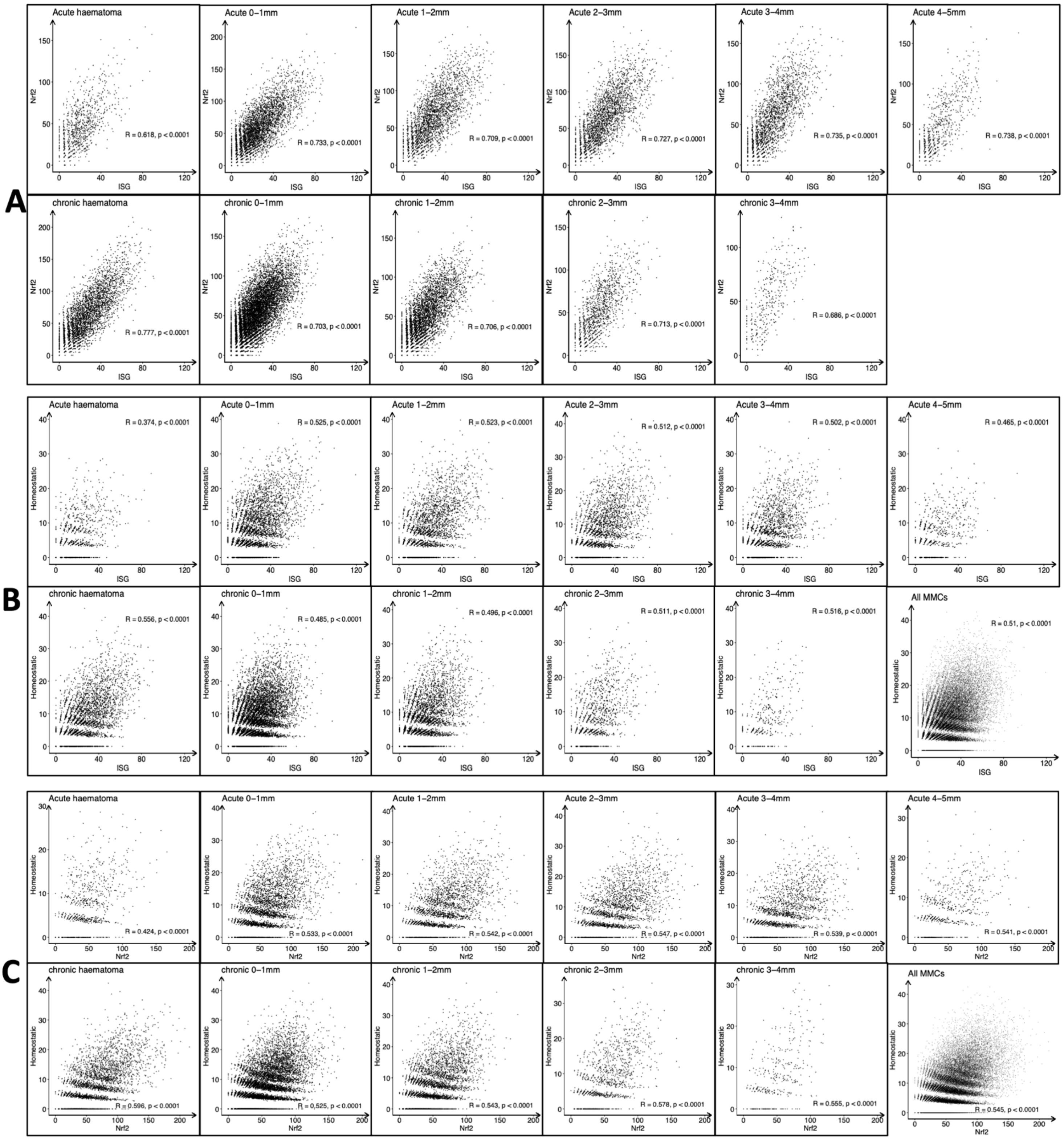

**Supplementary Figure 20: Association between Nrf2 target gene, ISG, and microglial homeostatic gene expression by human brain MMCs after ICH.** Related to figure 6. Scatter plots of summed expression of Nrf2 target genes, ISGs, and microglial homeostatic genes per MMC where each point represents an individual MMC from post-mortem human brain tissue after ICH. Pearson $R^2$ correlation coefficient for each of two plotted gene sets per cell. Each graph represents a time and distance from the haematoma. **A** Nrf2 vs. ISG expression (see main text for all cells at all distances pooled); **B** ISG vs. microglial homeostatic gene expression and **C** Nrf2 vs. microglial homeostatic gene expression. For each time/distance bin, the $R^2$ value of Nrf2 vs. ISG expression is approximately double that of either of these vs. a homeostatic gene set indicating that this correlation represents specific induction of these gene sets per cell, rather a general increase in gene expression/detection. N=4 acute, N=4 chronic.

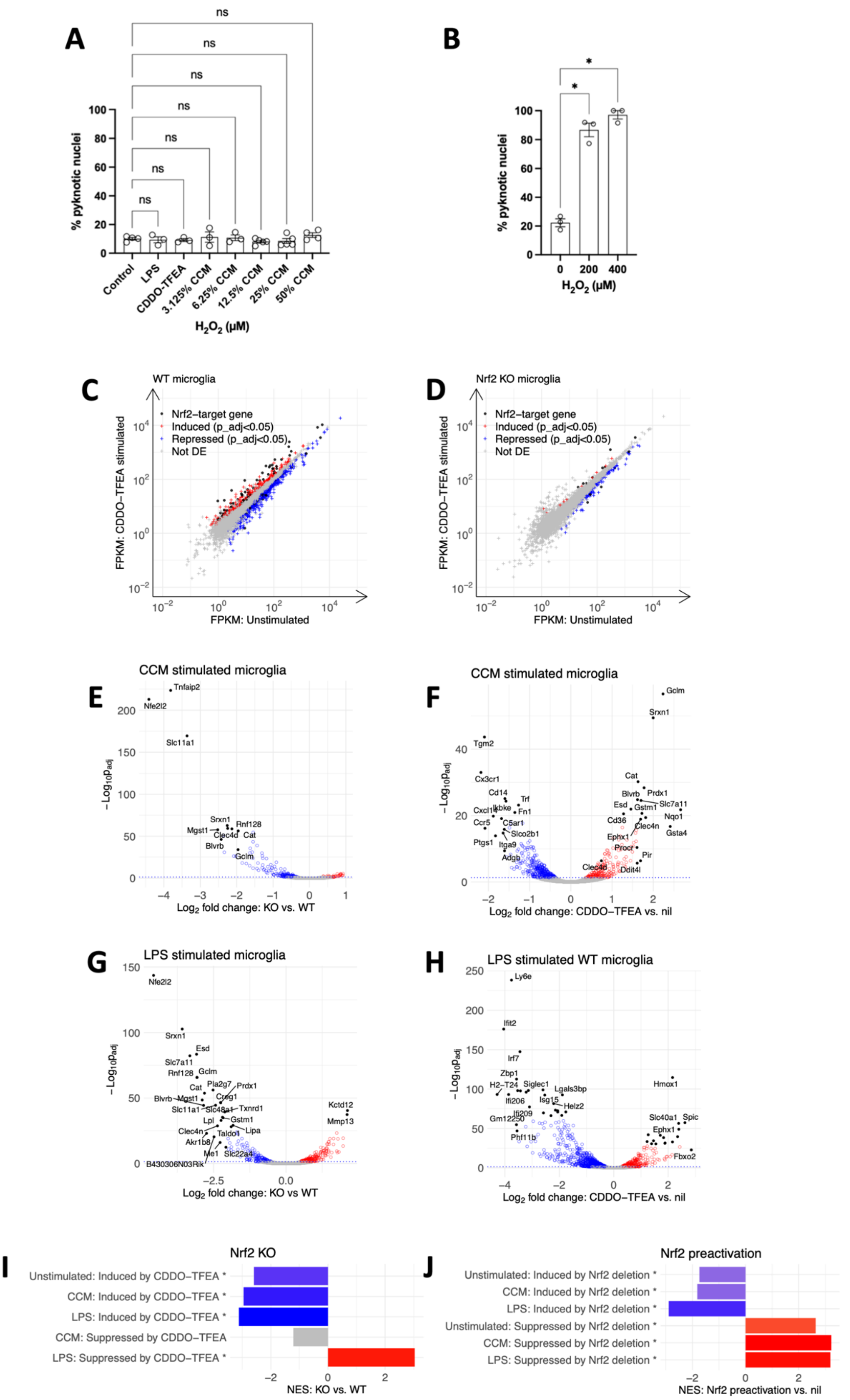

**Supplementary Figure 21: Effects of microglial Nrf2 deletion and Nrf2 activation after exposure to blood clot conditioned media or lipopolysaccharide**
Related to figure 7. **A-B** Bar charts showing percentage pyknotic nuclei for cultures of primary rat neurons and astrocytes stimulated by blood clot conditioned media (CCM),

LPS or CDDO-TFEA (**A**) or $H_2O_2$ (**B**) in separate experiments. Mean ± SEM, n=3-5. Scale bars 100μm. CCM One way ANOVA (**B**) F(7-22)=0.86, p=0.55. $H_2O_2$ One way ANOVA (**C**) F(2-6)=132, p<0.0001. **C-D** Scatter plot of average expression (FPKM) for CDDO-TFEA stimulated or unstimulated WT (**C**) or KO (**D**) microglia. Genes induced or suppressed ($p_{adj}$<0.05) with CDDO-TFEA are indicated in red or blue, respectively. Differentially expressed Nrf2 target genes identified in a previously published ChIP seq experiment (Malhortra, et al. 2010) are highlighted in black. **E-H** Volcano plots of DESeq2 log2 fold changes in KO versus WT microglial gene expression (**E**, **G**) or in WT microglia which have undergone Nrf2 preactivation by exposure to CDDO-TFEA (**F**,**H**). Microglia have been stimulated by blood clot conditioned media (CCM; **E, F**) or lipopolysaccharide (LPS; **G,H**). Red points indicate genes with increased expression in the experimental group, blue show genes which are reduced, and grey genes which are not differentially expressed ($p_{adj}$>0.05). n=2-3. **I-J** Bar plot of Gene Set Enrichment Analysis (GSEA) Normalised Enrichment Scores (NES) in KO compared with WT microglia (**I**) and microglia treated with CDDO-TFEA to preactivate Nrf2 vs. no pretreatment with CDDO-TFEA (**J**), for sets of genes that are significantly differentially expressed (*Benjamini-Hochberg $p_{adj}$<0.05) with CDDO-TFEA pretreatment (**I**) or Nrf2 deletion (**J**). This demonstrates the opposing effects of Nrf2 deletion and Nrf2 preactivation on microglia in unstimulated, CCM-stimulated and LPS-stimulated states. Red indicates positive enrichment; blue indicates negative enrichment; grey indicates no statistically significant enrichment following adjustment for multiple testing

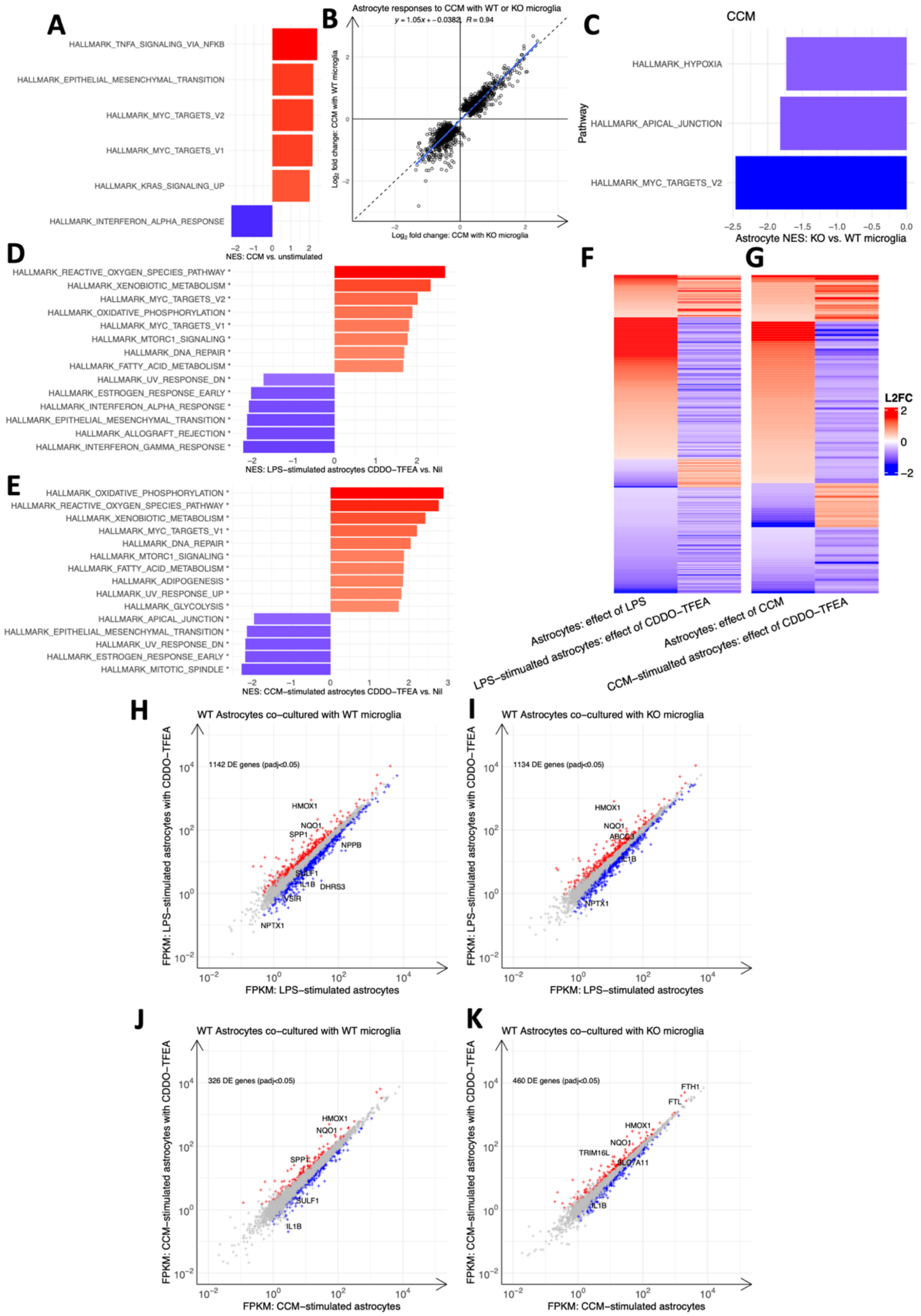

**Supplementary Figure 22: Astrocyte gene expression after exposure to blood clot conditioned media when co-cultured with WT or Nrf2-KO microglia.**

Related to figure 8. **A-C** Analyses of astrocyte gene expression when stimulated with blood clot conditioned media (CCM) and co-cultured with neurons and WT or KO microglia. **A** Bar chart of GSEA normalised enrichment score (NES) for the five most highly enriched or depleted Hallmark gene sets after CCM exposure (* $p_{adj}<0.05$). **B** Scatter plot of $log_2$ fold change in the expression of astrocyte genes which are differentially expressed following exposure to CCM when co-cultured with either WT or KO microglia. Solid blue line indicates fitted linear regression and 95% confidence interval. The formulaic expression of this line and the correlation coefficient are provided. Dashed black line of the form y=x provided for comparison. **C** Bar chart of GSEA NES for significantly enriched or depleted (* $p_{adj}<0.05$) Hallmark gene sets in WT astrocytes co-cultured with KO vs. WT microglia after CCM exposure. **D-E** Bar chart of GSEA NES for significantly enriched or depleted (* $p_{adj}<0.05$) Hallmark gene sets in astrocytes co-cultured with WT microglia and CDDO-TFEA or no CDDO-TFEA (nil) after LPS (**D**) or CCM (**E**) exposure. **F-G** Heatmap of astrocyte genes which are significantly ($p_{adj}<0.05$) differentially expressed after exposure to LPS (**F**) or CCM (**G**) and CDDO-TFEA. This indicates that CDDO-TFEA both augments and suppresses gene expression in a stimulus-dependent fashion. **H-K** Scatter plot of gene expression (FPKM) in astrocytes stimulated with LPS (**H-I**) or CCM (**J-K**) and co-cultured with either Nrf2-WT (**H, J**) or Nrf2-KO (**I,K**) microglia. Each graph is annotated with the number of significantly differentially expressed (DE) genes per comparison (DEseq2 $p_{adj}<0.05$). These show similar modulation of astrocyte gene expression by CDDO-TFEA when cultured with either microglial genotype. Red points indicate genes induced by CDDO-TFEA and blue genes repressed by CDDO-TFEA. n=3 independent biological repeats.

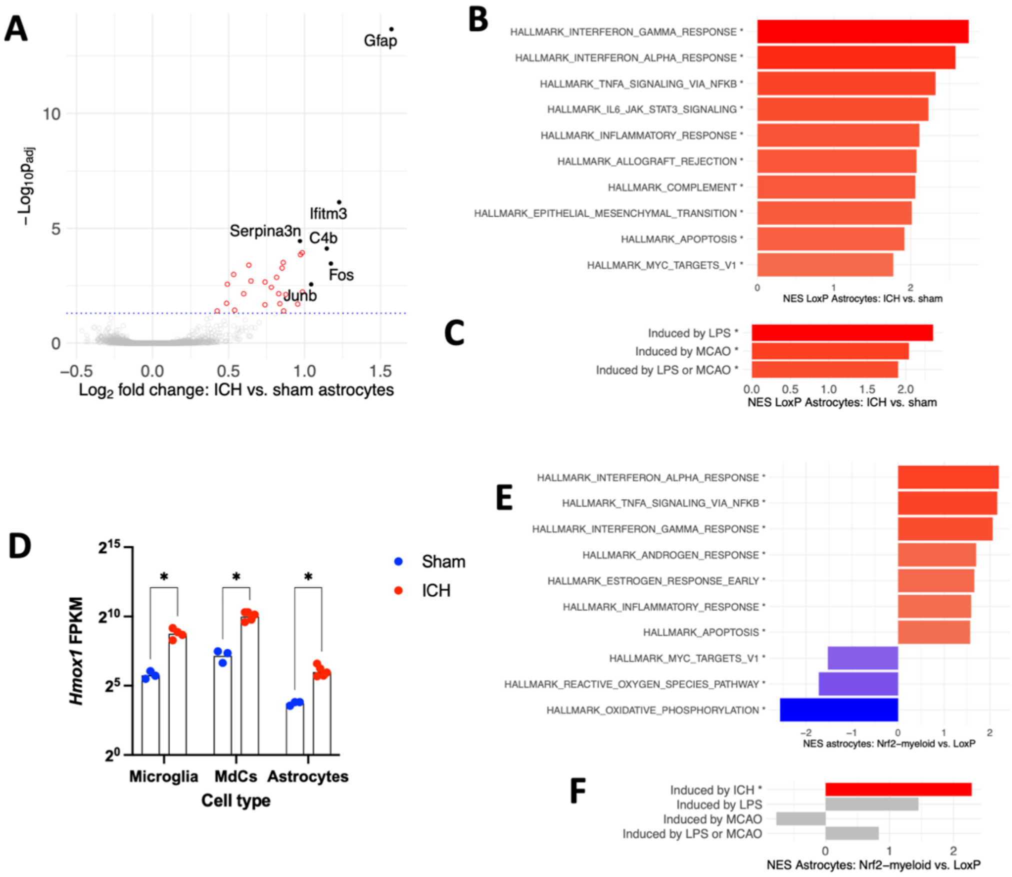

**Supplementary Figure 23: Effects of ICH and MMC specific Nrf2 deletion on astrocyte gene expression in vivo.**

Related to figure 8. **A** Volcano plot of DESeq2 log2 fold changes in astrocyte gene expression of FACS sorted astrocytes of LoxP mice 3 days after ICH or sham surgery, with genes filtered to remove those potentially influenced by MMC contamination (see methods). Red points indicate genes with increased expression after ICH and grey genes which are not differentially expressed (padj>0.05). No genes were significantly reduced in expression after ICH. **B-C** Bar plot of Gene Set Enrichment Analysis (GSEA) Normalised Enrichment Scores (NES) in ICH compared with sham astrocytes. **B** Top 10 Hallmark gene sets that are significantly positively or negatively enriched (*padj<0.05). **C** Previously curated[1,2]. **D** Expression of *Hmox1* in sorted astrocytes Microglia and MdCs of sham and ICH mice measured in fragments per kilobase of transcript per million mapped reads (FPKM) demonstrates induction of *Hmox1* by astrocytes at a lower expression level than in MMCs, concordant with our analyses of human post-mortem tissue. Geisser-Greenhouse corrected repeated-measures mixed-effects REML model: cell-type $F(1.2\text{-}7.5)=70.4$, $p<0.0001$; ICH $F(1,7)=52.9$, $p=0.0002$; cell-type x ICH $F(1.2\text{-}7.5)=40.3$, $p=0.0002$. Post-hoc comparison of ICH effect per cell-type using the Holm-Šidák method; Microglia $p=0.0015$; MdCs $p=0.0002$; Astrocytes $p=0.0015$. **E-F** Bar plot of GSEA NES in astrocytes from $Nrf2^{\Delta MMC}$ or LoxP mice with ICH. **E** Top 10 Hallmark gene sets that are significantly positively or negatively enriched (*padj<0.05). **F** Genes induced by ICH in **A** and the same previously curated gene sets from **C**. This indicates that Nrf2 deficiency drives a higher enrichment of genes that are induced in astrocytes specifically by ICH. Red indicates positive enrichment; blue indicates negative enrichment; grey indicates no statistically significant enrichment following adjustment for multiple testing. n=3 LoxP sham, n=6 LoxP ICH astrocytes and MdCs, n=5 LoxP ICH microglia, n=6 KO ICH.

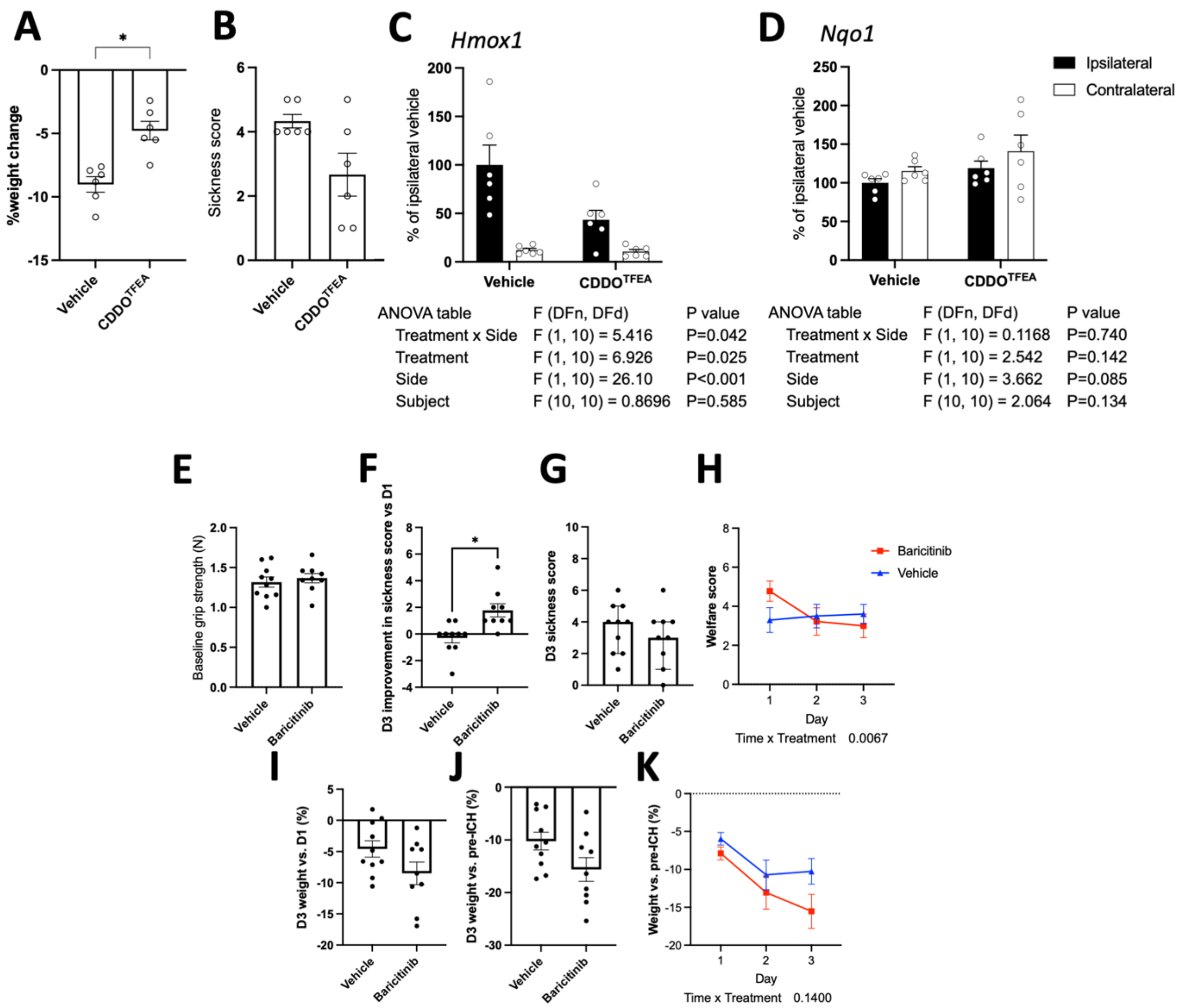

**Supplementary Figure 24: Additional outcomes of mice with ICH treated with CDDO-TFEA, Baricitinib or vehicle following ICH.**

Related to figure 9. **A** Weight change at 24h following ICH compared with pre-ICH in mice treated with CDDO-TFEA or vehicle. Two-tailed t-test with Welch's correction t=4.5, df=9.7, p=0.0013. **B** Sickness score at 24h following ICH where higher score indicates greater sickness. Mann-Whitney U test U=7; p=0.097. **C-D** Bar chart of expression of *Hmox1* (**C**) and *Nqo1* (**D**) measured by PCR and values normalised to the mean expression in ipsilateral samples of mice treated with vehicle are shown. n=6/group. Two-way repeated measures ANOVA tables are shown **E-J** Supplementary behavioural outcomes for KO mice with mononuclear myeloid cell Nrf2 deficiency treated with enteral baricitinib or vehicle at ICH onset, day 1 and 2 post-ICH. **E** Baseline pre-ICH grip strength (N). **F** Improvement in sickness score at day 3 from day 1 after ICH Mann-Whitney U test U= 8.5; p=0.0014. **G** sickness score at day 3 Mann-Whitney U test U= 35.5; p=0.45. **H** Daily sickness score after ICH. **I** Percentage weight change at day 3 compared with day 1 after ICH Welch's t-test t=1.7, df=15.0, p=0.10. **J** Percentage weight change at day 3 compared with pre-ICH Welch's t-test t=1.92, df=15.1, p=0.073. **K** Daily change in weight after ICH. n=10 vehicle, n=9 Baricitinib. Nine KO mice were culled following complications of the gavage procedure prior to outcome data collection, and so their data are not presented (n=4 veh, n=5 baricitinib). Two-tailed t-tests with Welch's correction (**E, F, H, I**). All error bars +/- SEM.

**Supplementary Methods: Quantification of DAB and Fast Red-stained areas in dual stained tissue.**

Related to figure 2. The unprocessed original image (**A**) is first used to generate a binary feature map (**B**). Next, the original image is then split into DAB (**C**) and Fast Red (**D**) channels. These are then converted to binary images (**E, F**). A composite overlay of these (**G**; Fast Red stain coloured red, DAB stain coloured blue) shows strong fidelity with the original image. Overlaying of the feature map on to this image (**H**) shows accurate segmentation of individual cells. Scale bars 30µm.

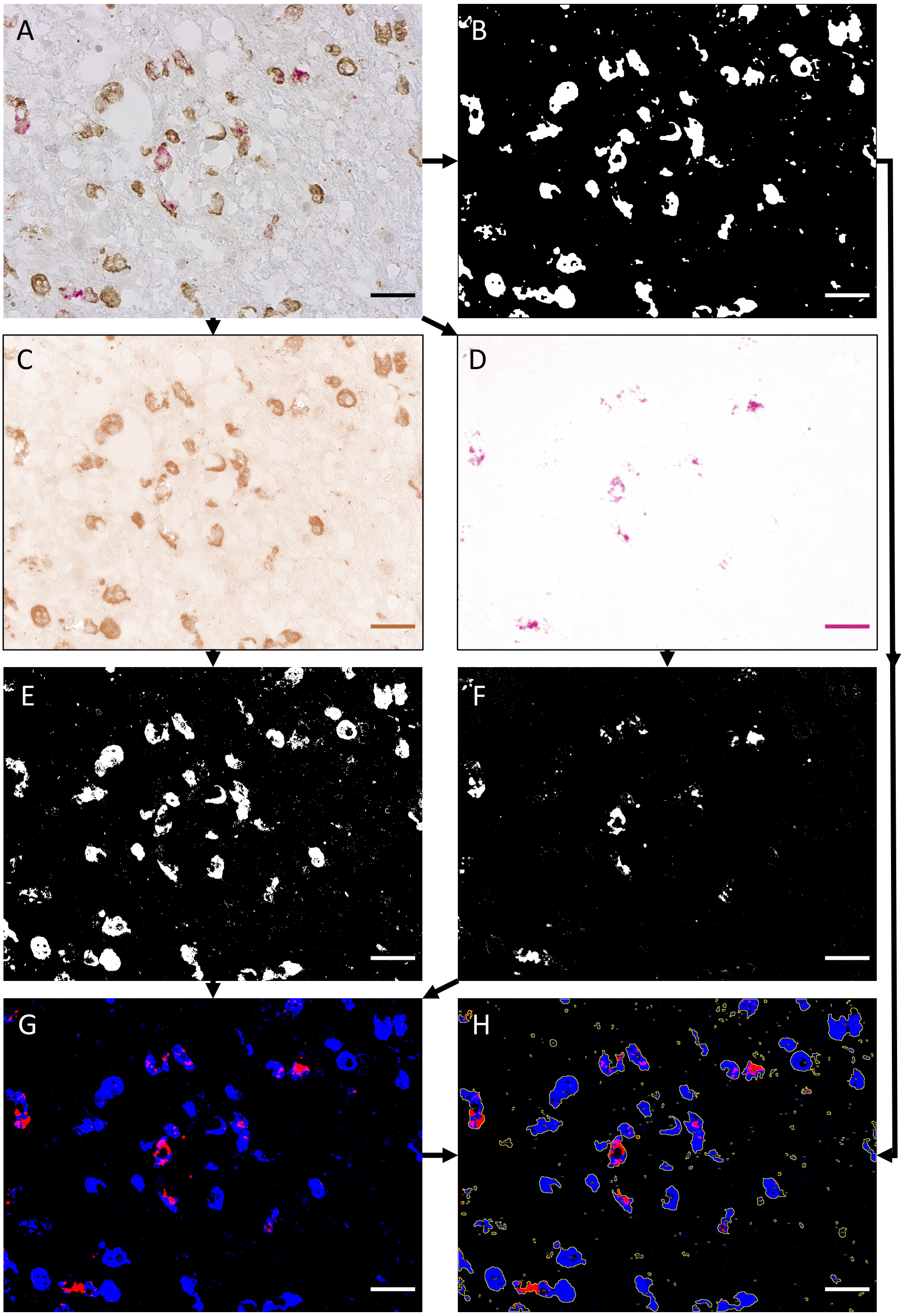
